# Quantitative fluorescence lifetime imaging of protein microenvironments by genetically encoded compact benzodiazole probes

**DOI:** 10.64898/2026.10.13.758698

**Authors:** Haocheng Qianzhu, Stephen M. Butler, Edan Habel, Elwy H. Abdelkader, Nathan Paul, Ryan Woolsey, Xuanhui Zhou, Rory M. Martin, Katrina A. Jolliffe, Gottfried Otting, Thomas Huber

## Abstract

Quantitative fluorescence lifetime imaging microscopy (FLIM) is limited by a lack of probes that combine single-residue specificity, minimal steric perturbation, visible-light excitation, and strong environment-dependent lifetime contrast. We introduce benzodiazole-based non-canonical amino acids (BD-ncAAs) as compact, genetically encoded sensors for site-specific FLIM. Using an engineered pyrrolysyl-tRNA synthetase pair, we incorporate three different BD-ncAAs into proteins in living cells. Site-specific incorporation of BD-ncAAs into a nanobody at different positions preserved native structure and antigen binding. The lead probe, cyano-substituted CBDS, exhibits visible-light excitation, large Stokes shifts, and pronounced environment-dependent fluorescence lifetimes spanning 1 to 24 ns. Under wash-free, live-cell FLIM conditions, CBDS quantitatively resolves localized structural microenvironments and transmembrane hydration gradients, while also distinguishing distinct states during protein secretion in eukaryotes. BD-ncAAs present genetically encoded, minimally perturbing reporters for tracking target proteins, detecting interactions with binding partners, and quantifying site-specific solvent accessibility in living cells by quantitative FLIM.

## Introduction

Fluorescence lifetime imaging microscopy (FLIM) provides a robust, concentration-independent readout that is intrinsically sensitive to the local molecular environment.^1–3^ Time-correlated single-photon counting (TCSPC)-FLIM has been shown to resolve nanosecond-scale changes in excited-state dynamics with high precision, enabling the mapping of temperature, pH, local polarity, viscosity, and other biochemical parameters.^4–7^ In practice, however, much of this resolving power remains underutilized due to limitations of the available probes. Existing lifetime reporters either exhibit modest lifetime variations that are difficult to interpret quantitatively, or produce complex multiexponential decays arising from multiple emissive states, complicating their use for environmental sensing.^8^ In addition, short-wavelength excitation and emission limit compatibility with commonly used confocal platforms and increase background from autofluorescence. The implementation of quantitative FLIM in live cells is further challenged by a trade-off between targeting specificity and photophysical performance. Small-molecule dyes can report bulk cellular conditions but generally lack the ability to interrogate specific protein sites or transient interactions. Conversely, fluorescent proteins offer genetically programmable specificity but are subject to photophysical constraints and add substantial steric bulk that can alter the local environment being probed.

Genetically encoded fluorescent non-canonical amino acids (ncAAs) combine site-specific targeting with the environmental sensitivity of small-molecule fluorophores.^9–17^ Recent rotor-based ncAAs, such as AnapTh, enable real-time imaging of protein microenvironment changes through intensity and spectral responses, but the method is not directly suited to fluorescence-lifetime imaging.^16^ Other fluorescent ncAAs—especially DanAla and Acd—exhibit environment-dependent fluorescence lifetimes, but DanAla requires short-wavelength excitation,^10^ while the long lifetime of Acd in aqueous solution can be substantially shortened in hydrophobic protein environments,^12, 14^ increasing overlap with endogenous fluorescence backgrounds. Moreover, the relatively bulky fluorophores of these ncAAs compromise their compatibility with functionally constrained protein sites, limiting the range of native microenvironments accessible to probing. Their side-chain volumes are substantially larger than those of canonical aromatic residues such as Trp (Fig. 1b).

**Fig. 1.**
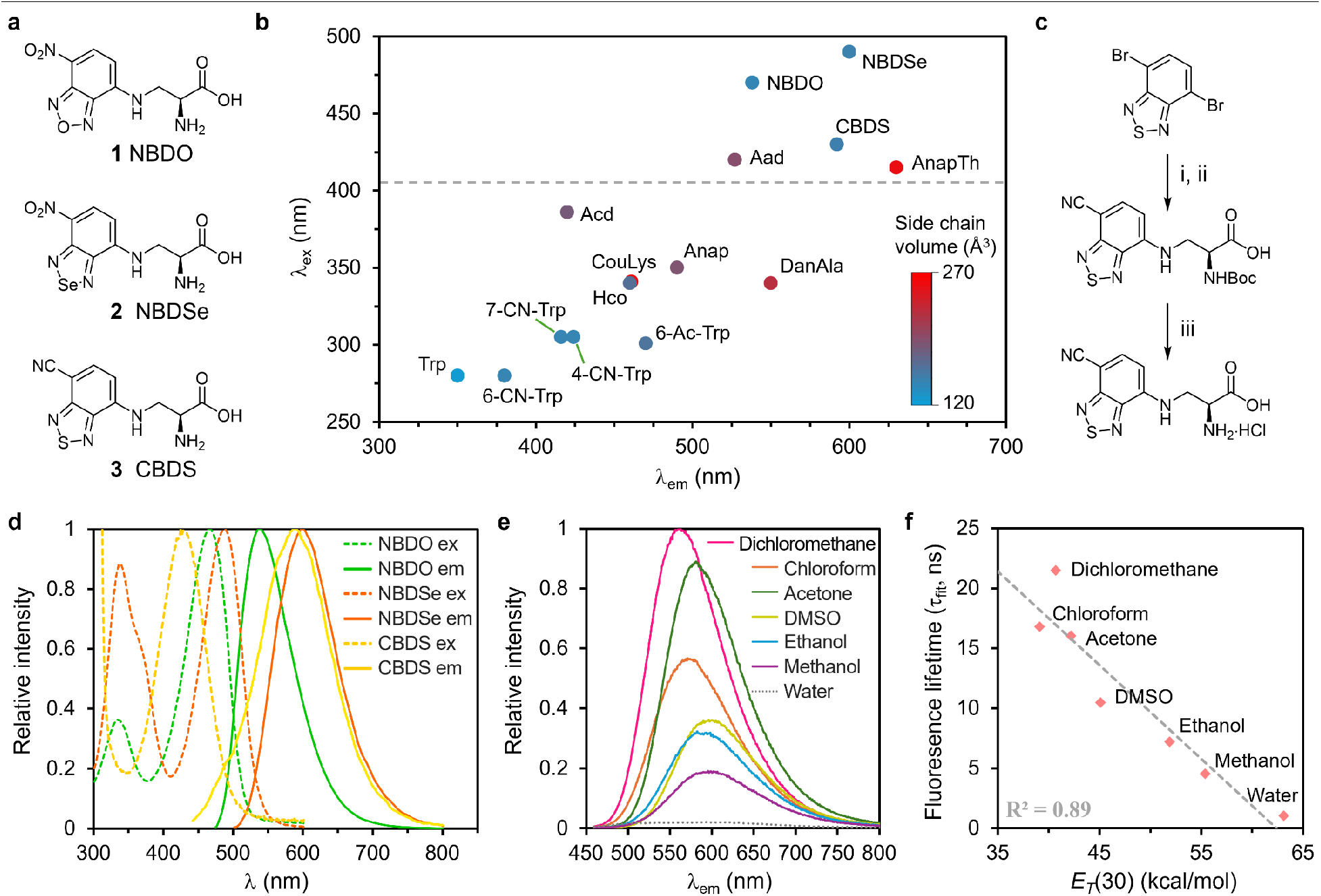
Design and photophysical properties of benzodiazole (BD)-bearing probes. **a**, Chemical structures of the non-canonical amino acids (ncAAs) used in this study: *N*-(7-nitro-2,1,3-benzoxadiazol-4-yl)-L-2,3-diaminopropionic acid (NBDO, **1**); *N*-(7-nitro-2,1,3-benzoselenadiazol-4-yl)-L-2,3-diaminopropionic acid (NBDSe, **2**); *N*-(7-cyano-2,1,3-benzothiadiazol-4-yl)-L-2,3-diaminopropionic acid (CBDS, **3**). **b**, Comparison of genetically encoded fluorescent amino acids by excitation maximum, emission maximum and side chain volume (calculated using the Slonimskii method,^18, 19^ represented by dot color along a blue-to-red color spectrum). Structures of compared ncAAs are provided in Extended Data Fig. 1. The gray dashed line indicates 405 nm, representing the shortest commonly available excitation wavelength on modern confocal microscopes. **c**, Synthetic route to CBDS: (i) CuCN, *N,N-*Dimethylacetamide, 200 °C, mW; ii) Boc-Dap-OH, DIPEA, DMSO, 100 °C, 12% over two steps; iii) 4M HCl in dioxane, THF, Δ, *quant*. **d**, Excitation and emission spectra of free CBDS, NBDO and NBDSe ncAAs in PBS. All three BD-ncAAs are excitable in the visible range and display large Stokes shifts. Emission spectra were recorded at the excitation maximum of each probe (470 nm for NBDO, 490 nm for NBDSe, and 430 nm for CBDS). **e**, Solvent-dependent emission spectra of Boc-protected CBDS. **f**, Fluorescence lifetime of Boc-protected CBDS as a function of solvent polarity, expressed as *E_T_*(30).

To fully exploit the resolving power of TCSPC-FLIM, an ideal genetically encoded probe should satisfy three criteria: First, it should produce large, reproducible and preferably monotonic lifetime shifts with a well-defined and quantitatively interpretable response to environmental changes; second, it should operate entirely within the visible spectral window to minimize both phototoxicity and cellular autofluorescence; and third, it should be sufficiently small to fit into buried protein sites without substantially perturbing their local conformation, thereby preserving native structure and function.

The benzodiazole-bearing fluorescent ncAAs (BD-ncAAs) of the present work constitute a class of compact, genetically encodable amino acids that enable quantitative TCSPC-FLIM. These probes can be excited using standard visible laser lines above 440 nm and exhibit large Stokes shifts (68–162 nm), which sets the emission wavelengths apart from the spectral regions dominated by cellular autofluorescence. The fluorescence lifetime of the leading BD-ncAA is extraordinarily sensitive to the polarity of the local environment, yielding well-separated lifetime distributions spanning 1.0 ns to 23.6 ns. Crucially, we report genetic encoding systems for incorporating the BD-ncAAs at defined sites in diverse protein scaffolds, enabling site-specific lifetime sensing and quantitative discrimination of biochemical states in living cells.

## Results

### Benzodiazole-based ncAAs

To develop new probes for site-specific FLIM sensing, we focused on ncAAs bearing the benzodiazole (BD) scaffold, a class of compact, “push-pull” fluorophores.^20^ We initially targeted previously reported NBDO (*N*-(7-nitro-2,1,3-benzoxadiazol-4-yl)-L-2,3-diaminopropionic acid; Fig. 1a),^21^ as the nitrobenzoxadiazole (NBD) fluorophore has a long and successful history stretching from its first report by Ghosh and Whitehouse in 1968.^22^ Remarkably, despite its small size, NBD is characterized by excitation and emission wavelengths that are comparable with much larger fluorophores (λ_ex_ = 486 nm, λ_em_ = 542 nm; EtOH),^23^ a high quantum yield (ϕ = 0.55),^23^ minimal cytotoxicity, and stability in biological systems.^20, 24–26^ We also chose NBDSe (*N*-(7-nitro-2,1,3-benzoselenadiazol-4-yl)-L-2,3-diaminopropionic acid; Fig. 1a), in which the ring oxygen is replaced by selenium. NBDSe has recently been reported as a sensitive fluorogenic sensor in synthetic peptides^21^ but its utility in live-cell protein imaging has remained unexplored.

To further maximize environmental sensitivity, we also designed CBDS (*N*-(7-cyano-2,1,3-benzothiadiazol-4-yl)-L-2,3-diaminopropionic acid; Fig. 1a), a novel ncAA featuring a 7-cyano-2,1,3-benzothiadiazol-4-yl moiety. Previous structure-activity studies on small push-pull fluorophores identified the combination of a cyano-substitution and a sulfur heteroatom in the benzodiazole ring to offer superior sensitivity to local polarity and hydrogen-bonding compared to nitro-substituted analogs.^27, 28^ By transforming this optimized scaffold into an ncAA side chain, we created a highly responsive member of the BD toolset. The synthesis of this new amino acid, CBDS, is reported for the first time. Similar to NBDO and NBDSe, CBDS was synthesized via nucleophilic aromatic substitution of the corresponding BD-halide with Boc-protected 2,3-diaminopropionic acid, followed by acidic removal of the Boc protecting group to afford the free amino acid (Fig. 1c).

In PBS buffer, the free amino acids NBDO, NBDSe, and CBDS exhibited excitation/emission maxima at 470/538 nm, 490/600 nm, and 430/592 nm, with Stokes shifts of 68 nm, 110 nm, and 162 nm respectively (Fig. 1d), and showed moderate-to-high molar extinction coefficients (Extended Data Fig. 1b, c). To characterize the intrinsic sensitivity of the BD scaffold to different chemical environments, we measured the fluorescence emissions and lifetimes of Boc-protected CBDS in solvents with a wide range of polarities. CBDS exhibited pronounced solvatochromism with progressively red-shifted emission maxima and diminished fluorescence intensity in more polar environments (Fig. 1e). The fluorescence lifetimes of CBDS showed marked dependence on solvent polarity, with a dynamic range spanning <2 ns (water) to >20 ns (dichloromethane (DCM)), and a linear correlation with the microscopic Dimroth–Reichardt empirical solvent polarity parameter *E_T_*(30) (fitted fluorescence lifetime (τ_fit_) = −0.78 × *E_T_*(30) + 49, R^2^ = 0.89; Fig. 1f; Extended Data Fig. 1d). The lifetime weakly correlated with the macroscopic bulk solvent dielectric constant (ε; R^2^ = 0.65; Supplementary Fig. 3) and the superior fit with *E_T_*(30) indicates that the excited-state relaxation of CBDS is predominantly governed by short-range solute-solvent interactions rather than bulk dielectric effects.

To further dissect the solvent contributions, we performed a multiple linear regression using the Kamlet−Taft solvent parameters α (hydrogen bond donor), β (hydrogen bond acceptor), and π* (polarizability; Extended Data Fig. 1d).^29, 30^ The resulting model provided a strong empirical description of the observed lifetimes (R^2^ = 0.97, adjusted R^2^ = 0.94), with negative coefficients for both α and β, indicating that increasing solvent hydrogen-bonding capacity is generally associated with a reduction in fluorescence lifetime. In particular, the marked decrease in fluorescence lifetimes observed in protic solvents (water, methanol, ethanol) compared to highly polar but aprotic solvents (DMSO) suggests that hydrogen-bond donation from the solvent to the charge-separated intramolecular charge-transfer (ICT) state facilitates highly efficient non-radiative decay pathways.^31^ In contrast, CBDS exhibited relatively short fluorescence lifetimes in ethylene glycol (EG)/water mixtures, with lifetimes of 1.89 ns in 1:1 EG/water and 3.11 ns in 9:1 EG/water (Extended Data Fig. 1e), suggesting that its lifetime is not strongly determined by bulk solvent viscosity. This confirms that the non-radiative decay from the ICT state of CBDS is strictly coupled to solvent interactions rather than diffusion-limited processes. This behavior is consistent with the classical push–pull architecture of benzodiazole fluorophores.^20, 32^ Consequently, we reasoned that the exquisite sensitivity of CBDS to its immediate environment makes it an ideal ncAA for quantitatively probing the local protein architecture and dynamic solvent accessibility of specific protein microenvironments.

### Installing BD-ncAAs in proteins

Previous studies have shown that BD fluorophores such as NBDO and NBDSe can probe protein environments, but only through synthetic or semi-synthetic approaches unsuitable for in-cell protein imaging.^21, 33, 34^ To address this, we established in vivo incorporation platforms for these BD-ncAAs based on our previous success in identifying aminoacyl-tRNA synthetase (RS) variants capable of incorporating tryptophan analogues into proteins.^15, 35, 36^ Specifically, we employed RS mutants derived from the ISO4-G1 pyrrolysyl-tRNA synthetase^37^ to direct their incorporation in live cells in response to a programmed amber stop codon during protein translation (Fig. 2a). This genetic code expansion system reliably installs the ncAAs in proteins at specific sites both in bacteria and eukaryotic cells.^38, 39^

**Fig. 2.**
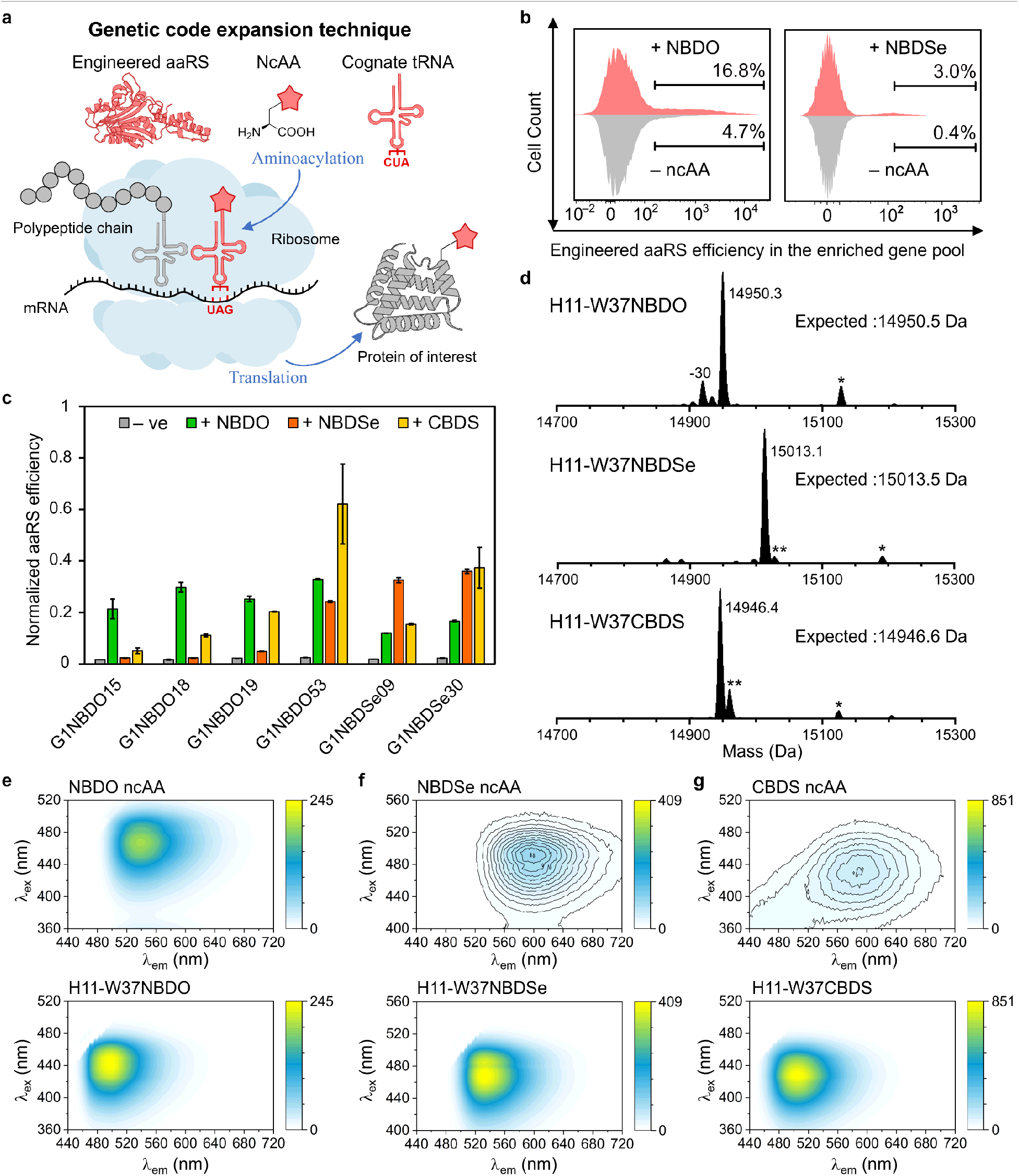
Site-specific incorporation of BD-ncAAs into proteins. **a**, Schematic of genetic code expansion used to incorporate BD-ncAAs in response to an amber stop codon. Engineered G1PylRS variants load the cognate G1-tRNA^Pyl^ with the ncAA, enabling site-specific incorporation during translation. **b**, Fluorescence-activated cell sorting (FACS) histograms from the final selection rounds used to enrich active and orthogonal G1PylRS variants for the incorporation of NBDO or NBDSe in *E. coli*. **c**, Activity and substrate scope of selected G1PylRS variants toward NBDO, NBDSe, and CBDS, assessed by amber suppression efficiency in the RFP reporter assay. **d**, Intact protein mass spectrometry of purified H11 nanobody variants confirms incorporation of BD-ncAAs at the amber-mutated Trp37 position. Major peaks match the expected masses of full-length proteins containing the corresponding ncAA. Peaks marked with one asterisk (*) indicate N-terminal gluconoylation (+178 Da) and two asterisks (**) indicate oxidation (+16 Da) under mass spectrometry conditions. Extended Data Fig. 2 shows clean incorporation without detectable modifications. **e**, **f**, **g**, Two-dimensional fluorescence excitation–emission spectra of BD-ncAAs recorded for the free amino acids (top) and after incorporation in the hydrophobic core of H11 (bottom), all measured at 10 μM. Contour lines in **f** and **g** are plotted at 1 fluorescence-unit intervals. The hydrophobic environment enhances the maximum emission intensity of NBDSe and CBDS approximately 30-fold and 100-fold, respectively.

We used a G1PylRS library previously shown to accept a broad range of structurally diverse ncAAs.^40^ Selection of suitable RS variants was performed in *E. coli* DH10B cells co-transformed with pBK-G1RS, encoding the G1PylRS mutant library, and pBAD-H6RFP, a reporter plasmid expressing red fluorescent protein mRFP1 with an amber codon inserted at the N terminus. Alternating positive and negative selection rounds employed fluorescence-activated cell sorting (FACS) to enrich RS variants that were both active and orthogonal in the presence of NBDO or NBDSe (Fig. 2b; Supplementary Fig. 4–6). Following multiple rounds of selection, 60 colonies for each ncAA were isolated, and RFP expression levels were evaluated (Supplementary Fig. 7, 8). Sequencing of the top-performing clones identified six G1PylRS variants that efficiently incorporated NBDO or NBDSe (Supplementary Table 3), establishing the first genetic code expansion system for these BD-ncAAs.

Sequence analysis of the top variants selected independently for NBDO and NBDSe revealed strong convergence, suggesting a shared mode of substrate recognition and possible polyspecificity within this benzodiazole scaffold class. To test this, we examined the activity of these variants toward NBDO, NBDSe, and CBDS. One variant isolated from the NBDO selection, G1NBDO53 (L124M, A221G, W237Y), efficiently incorporated NBDO, NBDSe, and CBDS (Fig. 2c). Notably, G1NBDO53 differs from the far more selective variant G1NBDO19 by only a single hydroxyl group (phenylalanine versus tyrosine at position 125).

To confirm site-specific incorporation of the three BD-based ncAAs, we expressed variants of the peptidyl–prolyl *cis*–*trans* isomerase FKBP1 A (FKBP12) by replacing the conserved Trp59 residue in the ligand-binding pocket and purified the proteins for intact mass spectrometry. To improve amber suppression efficiency for preparative protein production, the RS mutants G1NBDO53 and G1NBDSe09 were subcloned into pRSF plasmids, yielding pRSF-G1NBDO53 for NBDO and CBDS incorporation and pRSF-G1NBDSe09 for NBDSe. The plasmids were co-transformed with pCDF plasmids encoding the amber-interrupted reporter gene into *E. coli* B95.ΔAΔ*fabR*.^41^ Mass spectrometry of the purified FKBP12 variants showed that, for all three ncAAs, the main products matched the expected masses of the full-length proteins containing the corresponding ncAAs (Extended Data Fig. 2a–c). For NBDO, a minor peak about 30 Da below the expected mass is indicative of in vivo reduction of the nitro group to an amino group consistent with previous reports of nitroreductase-mediated reduction of aromatic nitro groups in *E. coli*, particularly for nitrotyrosine analogues.^42^ In contrast, the corresponding NBDSe variant showed only minimal evidence of nitro group reduction. CBDS remained fully stable in the purified protein, with no detectable chemical modification.

### Nanobody with site-specific BD-ncAAs

A stringent test of the structural compatibility of these ncAAs is their incorporation into nanobodies, which are small, single-domain antibody fragments (∼15 kDa) widely used for high-affinity intracellular targeting.^44^ Fusion of nanobodies with a protein fluorophore or chemical ligation with conventional tags present major modifications that easily interferes with folding and antigen binding.^45, 46^ As a model scaffold, we chose the H11 anti-ArcNL nanobody, a structurally well-characterized binding partner of the ligand-binding pocket in the N-lobe of human Arc (ArcNL).^44^ Substituting the buried core residue Trp37 with each of the BD-ncAAs enabled their fluorescence properties to be evaluated in a defined hydrophobic protein environment. Mass spectrometric analysis of the purified H11 variants confirmed clean incorporation of the corresponding BD-ncAAs at the intended positions (Fig. 2d; Extended Data Fig. 2). CBDS showed the strongest environmental response and was additionally used to probe the semi-exposed sites of Phe48 and Tyr60, as well as Trp54 in the antigen-binding loop CDR2.

At the site of Trp37 in H11, all three BD-ncAAs exhibited the pronounced solvatochromic response expected for the hydrophobic protein interior. Two-dimensional excitation–emission spectra revealed significant blue shifts in both the excitation and emission maxima compared to the free amino acids in aqueous buffer (PBS), confirming the location of the probes in a low-dielectric environment (Fig. 2e–g, Supplementary Fig. 9–11). The magnitude of fluorescence enhancement differed significantly between the three derivatives. NBDO showed only a marginal 1.2-fold increase in maximum emission intensity whereas NBDSe and CBDS demonstrated 30-to 120-fold signal enhancement, respectively. This strong response to environmental polarity combined with low background fluorescence in aqueous solution (Extended Data Fig. 2g) renders NBDSe and CBDS attractive as “wash-free” fluorescence probes that can be monitored in the presence of an excess of the free amino acids.

Incorporation of BD-ncAAs into H11 yielded variants with native-like structure and activity. Crystal structures of H11-W37NBDO (PDB ID: 9PZZ) and H11-W37NBDSe (PDB ID: 9Q00) in complex with ArcNL superimpose to the WT structure (PDB ID: 8QF4)^43^ with C^α^ RMSDs below 0.5 Å (Fig. 3a,b) and show preserved buried surface area and key polar contacts in the nanobody-epitope interface (Extended Data Fig. 3). Biolayer interferometry (BLI) confirmed that the W37-BD-ncAA mutants retained nanomolar binding affinity for ArcNL (Fig. 3c), with binding kinetics closely matching those of wild-type H11 (Extended Data Fig. 3f).^47^ Interestingly, the structural overlays reveal that BD-ncAAs, owing to their amine linker, are pseudoisosteric to tryptophan (Fig. 3a; Extended Data Fig. 3b): the six-membered BD ring takes the site of the six-membered tryptophan ring, orienting the nitro group toward the buried residue Glu6 and subtly shifting its position. The five-membered BD ring lies above the Trp *N*^ε^, changing the χ_1_ angle of the adjacent residue Val81. Far-UV circular dichroism (CD) spectra of H11-W37NBDO and H11-W37NBDSe were similar to the wild-type spectrum (Fig. 3d) and the melting temperatures differed by <2°C from the wild-type protein. Incorporation of CBDS into H11 at Phe48, Tyr60, and Trp54 likewise maintained the function of the nanobody. BLI analysis of these H11-CBDS mutants also revealed fully retained nanomolar binding affinity for ArcNL (Fig. 3e; Extended Data Fig. 3g).

**Fig. 3.**
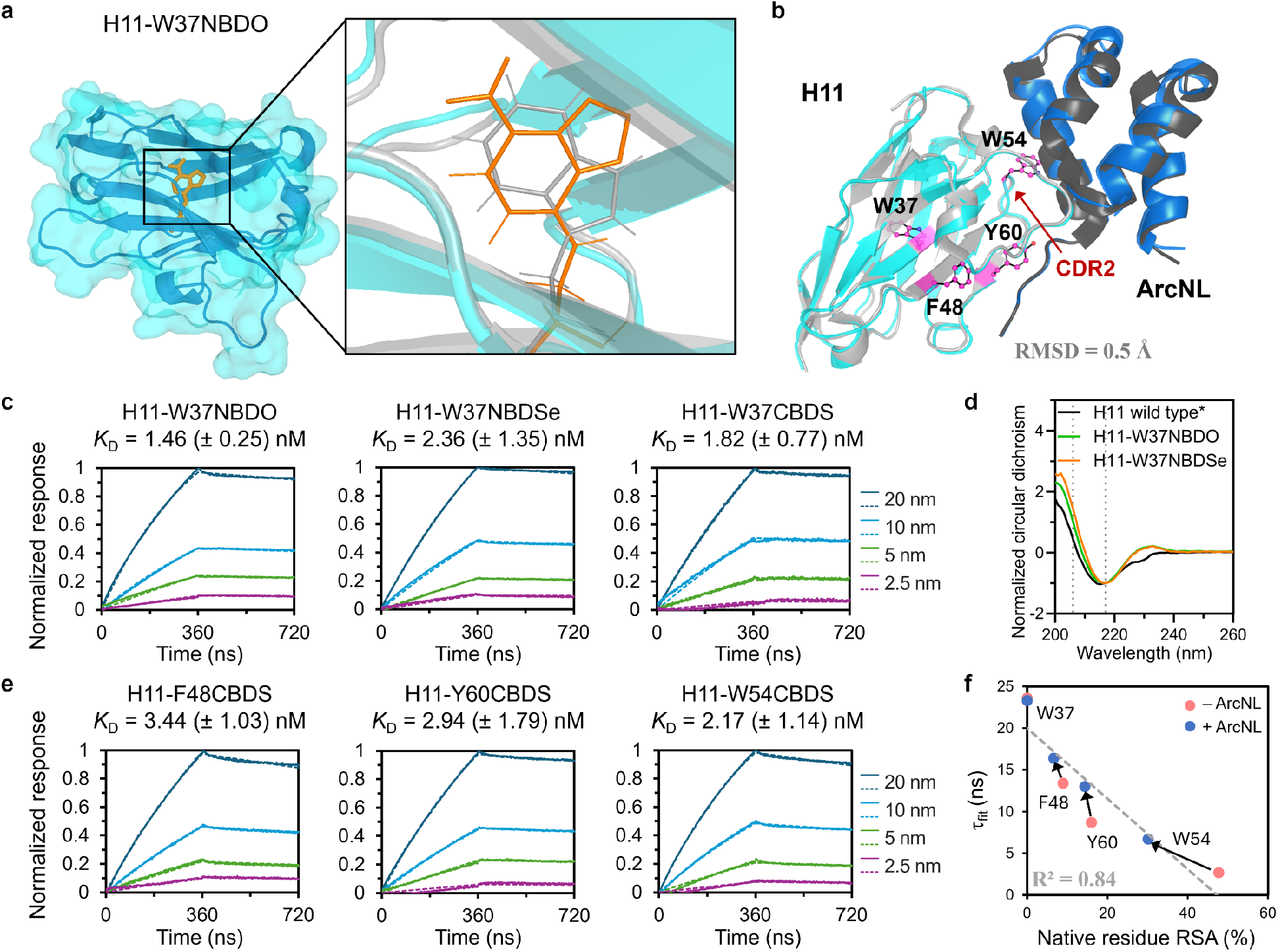
BD-ncAAs preserve nanobody structure and antigen binding after site-specific incorporation. **a**, Crystal structure of the nanobody H11 bearing NBDO at position 37, in complex with ArcNL (PDB 9PZZ). Enlarged view of the superimposition with wild-type H11 illustrates the structural match of the substituted BD-ring (orange sticks) with the native indole ring (gray sticks). **b**, Superposition of the ArcNL-bound H11-W37NBDO complex (blue/cyan) with ArcNL-bound wild-type H11 (black/gray; PDB 8QF4)^43^ shows close structural conservation. Chosen sites for additional environment probing with CBDS are shown in magenta. **c**, Biolayer interferometry analysis of H11 variants carrying BD-ncAAs shows retention of nanomolar binding affinity to ArcNL. **d**, Far-UV circular dichroism spectra of wild-type and BD-labeled H11 variants indicate preservation of the nanobody fold. Wild-type H11 treated with TCEP (*) was included as a reference control to account for the previously reported contribution of aromatic residue–disulfide interactions to the atypical ellipticity above 210 nm.^43^ **e**, Biolayer interferometry analysis of H11 mutants with CBDS incorporated at different locations confirms conserved binding affinity to ArcNL. **f**, Fluorescence lifetimes of purified H11-CBDS mutants measured in PBS buffer (red) and after addition of purified ArcNL (blue) obtained by reconvolution fitting of the TCSPC decay (τ_fit_), plotted against the relative solvent-accessible surface area (RSA) of each labeled residue, calculated from crystal structures (PDB 8QF3 and 8QF4) as a proxy for the local microenvironment polarity.

We next examined how the photophysical response of CBDS relates to the structural environment of the labeled residue. To quantify local solvent exposure, we calculated the relative solvent-accessible surface area (RSA) of each labeling site using crystal structures (PDB 8QF3 and 8QF4; Fig. 4c). For the purified H11-CBDS mutants measured in PBS buffer, both in the absence and presence of purified ArcNL, fluorescence lifetimes (τ_fit_) were obtained by reconvolution fitting of the TCSPC decay curves. We observed a strong negative correlation between the average fluorescence lifetime and the RSA (τ_fit_ = −0.42 × RSA + 20, R^2^ = 0.84; Extended Data Fig. 3). This robust inverse relationship corroborates our fundamental photophysical model that residues with greater solvent exposure are subject to more efficient non-radiative decay and correspondingly shorter lifetimes.

**Fig. 4.**
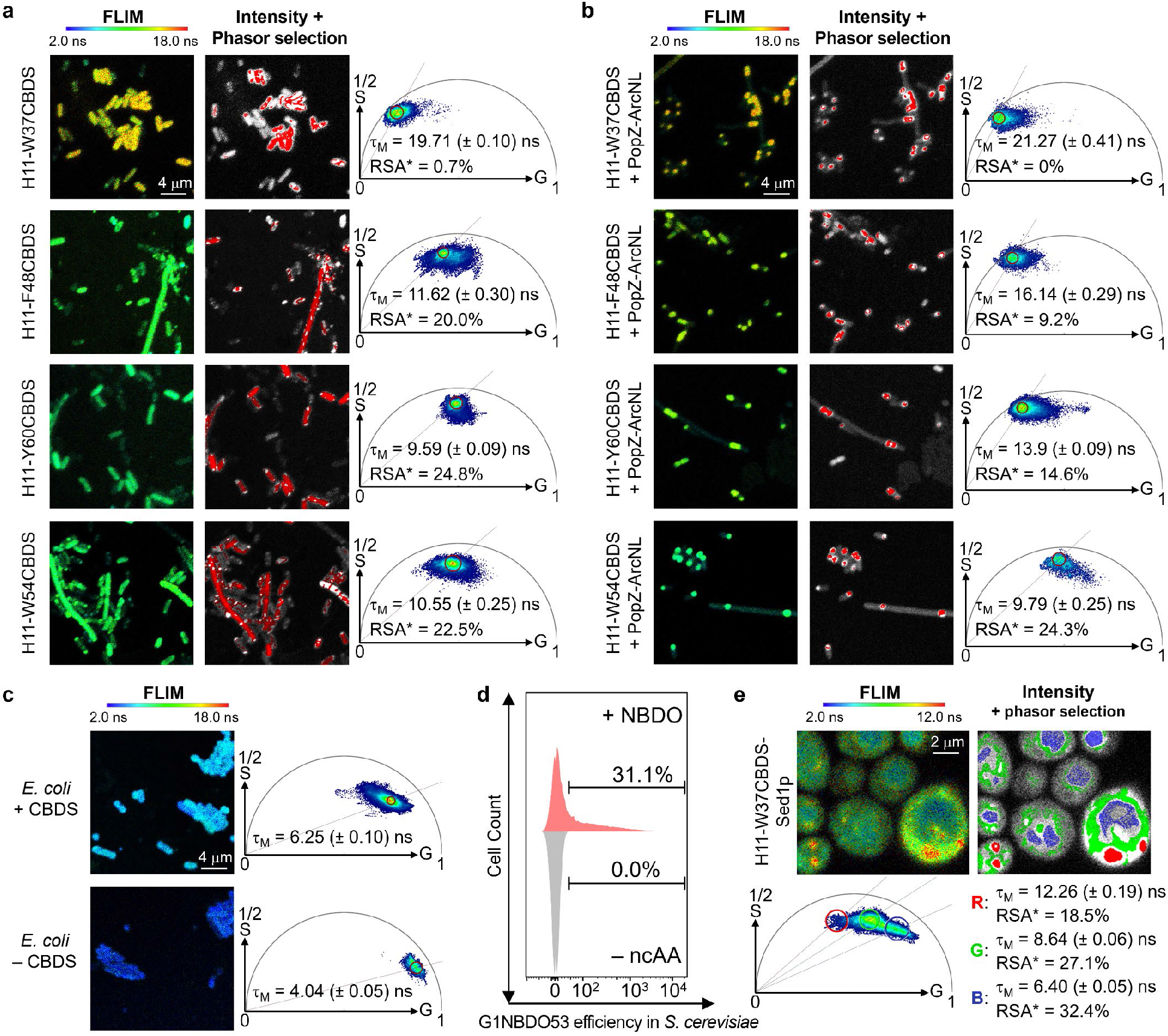
Wash-free FLIM imaging resolves nanobody microenvironments using genetically encoded BD-ncAAs. **a**, FLIM analysis of *E. coli* cells cytosolically expressing H11 variants containing BD-ncAAs at defined positions. Lifetime maps are shown in pseudocolor. Corresponding phasor plots show fluorescence lifetime distributions across the field of view. The corresponding fluorescence intensity image in gray scale is shown with the phasor-selected population overlaid in red. For determination of the modulation lifetime (τ_M_), a fluorescence population was selected directly on the phasor plot. The photons associated with the selected phasor population were then mapped back to their spatial origins, with the corresponding pixels highlighted in red over the fluorescence intensity image shown in grayscale. Reported τ_M_ values are mean ± s.d. from n = 3 independent imaging experiments. The estimated relative solvent-accessible surface area (RSA*) was back-calculated from the fitted relationship between τ_fit_ and RSA. All images are shown at the same magnification. **b**, Co-expression of ArcNL fused to the localization tag PopZ recruits the protein complex to the two poles of *E. coli* cells. H11 mutants carrying CBDS are captured at these locations, confirming that ArcNL binding occurs inside the cell. **c**, FLIM analysis of *E. coli* cells in the presence and absence of CBDS incubation demonstrates clear separation of the free CBDS and cellular autofluorescence from the CBDS-labeled protein signals. **d**, Histogram for the Electra-TAG–mScarlet readthrough assay demonstrating NBDO incorporation in *S. cerevisiae*. **e**, FLIM imaging of H11-W37CBDS displayed on yeast cell surfaces via a Sed1 glycosylphosphatidylinositol anchor (Sed1p) identifies three major lifetime populations.

### In-cell resolution of nanobody microenvironments via wash-free FLIM

The characterized H11 mutants were used as model proteins for live-cell imaging to demonstrate that the environmental sensitivity of CBDS in particular is compatible with FLIM. We first expressed four H11-CBDS variants in *E. coli*. Incorporated CBDS produced robust intracellular fluorescence signals in H11-expressing cells without the need for extensive washing. FLIM images were acquired with 440 nm excitation at a laser repetition rate of 20 MHz, with the excitation power adjusted for each sample to maximize photon counts without saturating the FLIM detector (Fig. 4a), with lifetime maps and phasor plots revealing distinct lifetime distributions that varied according to the specific incorporation site. FLIM data were analyzed using phasor plot analysis. The dominant population was selected directly on the phasor plot, and the photons associated with this population were mapped back to their spatial origins, with the corresponding pixels highlighted in the fluorescence image (Fig. 4a). Due to the presence of multiple fluorescent species and heterogeneous decay kinetics, direct fitting of the FLIM data was difficult. Instead, the modulation lifetime (τ_M_) of all photon events within the selected phasor region was used to provide a model-independent descriptor of the average fluorescence decay behavior (Fig. 4a). Although τ_M_ is not strictly equivalent to the τ_fit_ obtained from time-domain analysis, it provides a practical metric for comparing relative lifetime trends across the mutant series under heterogeneous intracellular conditions.

To verify that the nanobodies remained functional and capable of engaging their targets in vivo, we also co-expressed ArcNL fused to a PopZ localization tag, which locates to the bacterial cell poles. Upon co-expression, the H11-CBDS mutants were observed at the cell poles, confirming that high-affinity binding to ArcNL is maintained in the intracellular environment (Fig. 4b; Extended Data Fig. 4b). τ_M_ values were calculated from all photon events within the selected phasor region.

The in vitro RSA–τ_fit_ relationship established using purified H11-CBDS mutants was then used as a calibration to translate the intracellular τ_M_ measurements into estimated relative solvent-accessibility values (RSA*) for the corresponding sites. For most incorporation sites, the resulting RSA* values were closely comparable to the solvent accessibility expected from the corresponding structural sites measured in vitro, indicating that the site-dependent environmental response of CBDS is largely preserved in the bacterial cytosol. The major exception was W54 in the absence of ArcNL, which displayed a substantially lower RSA* than expected from its structural accessibility. This deviation suggests that CBDS at W54 experiences additional nonspecific interactions in the bacterial cytosol that restrict probe accessibility, an effect that is relieved upon ArcNL binding. Consistent with this interpretation, the direct correlation between intracellular τ_M_ and structural RSA was weaker than that observed in vitro (R^2^ = 0.55), but improved substantially when the unbound W54 mutant was excluded (R^2^ = 0.73), with the slope approaching the in vitro relationship (Supplementary Fig. 16). Together, these results indicate that CBDS retains its solvent-dependent lifetime response in the intracellular environment and that the FLIM-derived τ_M_ values provide biologically informative estimates of local solvent accessibility. Deviations from the structural RSA–lifetime relationship can further reveal site-specific effects of the intracellular environment, such as nonspecific interactions that locally shield the fluorophore from solvent.

For comparison, we also examined other BD-ncAAs. Incorporation of NBDSe in the buried W37 site of H11-W37 yielded τ_M_ = 4.5 ns, which is substantially longer than for the free amino acid (1.3 ns) and consistent with reduced solvent exposure in the protein core (Supplementary Fig. 22).

The orthogonality of the G1Pyl-based genetic encoding system has previously been demonstrated in eukaryotic hosts.^48^ To evaluate the performance of CBDS for FLIM in yeast, we constructed a G1NBDO53/G1-tRNA knock-in strain of *S. cerevisiae* BY4741^49^ (Extended Data Fig. 4e). Using an amber readthrough reporter, 31.1% of the total cell population showed successful NBDO incorporation, corresponding to a translational efficiency of 35.9% (Fig. 4d; Extended Data Fig. 4e, 4f). We next examined whether CBDS could report on folding-associated microenvironments during protein secretion and display in yeast. CBDS was incorporated either at W37, which provides a folding-sensitive reporter, or at the N terminus of H11, and the proteins were expressed as Sed1p^50^ fusions in *S. cerevisiae* (Fig. 4e; Extended Data Fig. 4e, 4h). FLIM resolved three major lifetime populations for H11-W37CBDS-Sed1p (τ_M_ = ∼12, 9, and 6 ns), while H11-Nt-CBDS-Sed1p showed closely corresponding ∼9- and ∼6-ns populations. The ∼9-ns population was broadly distributed around the cell periphery and intracellular regions, consistent with CBDS experiencing a relatively solvent-accessible protein-associated environment, although the images alone do not distinguish successfully surface-displayed protein from material retained within the secretory pathway. By contrast, the longer ∼12-ns population was substantially more prominent for W37-labeled H11 than for the N-terminally labeled construct, indicating that burial at W37 can generate a more protected folding-associated environment; however, its lifetime remained well below the ∼20-ns value of correctly folded H11-W37CBDS, consistent with a non-native or partially folded state. The shortest-lifetime ∼6-ns population localized predominantly to the vacuole-like compartment and was also detectable, but much less prominently, in cells cultured with free CBDS alone. Its strong enrichment when CBDS was incorporated into proteins therefore suggests that a substantial fraction of the labeled protein is trafficked to the vacuole, likely for degradation.

### Quantitative mapping of membrane hydration gradients

To demonstrate the broader utility of CBDS for interrogating complex cellular environments, we targeted the integral outer membrane protein OmpX. An artificial surface-display loop (SDL) consisting of seven alanine residues was inserted between S53 and S54,^52^ with the fifth alanine residue of the SDL substituted with CBDS. We also incorporated CBDS at two native sites, including the periplasm-facing W76 and a site more deeply embedded in the hydrophobic lipid environment, W140.^51^ These three sites were selected to sample distinct physical environments across the transmembrane protein (Fig. 5a).

**Fig. 5.**
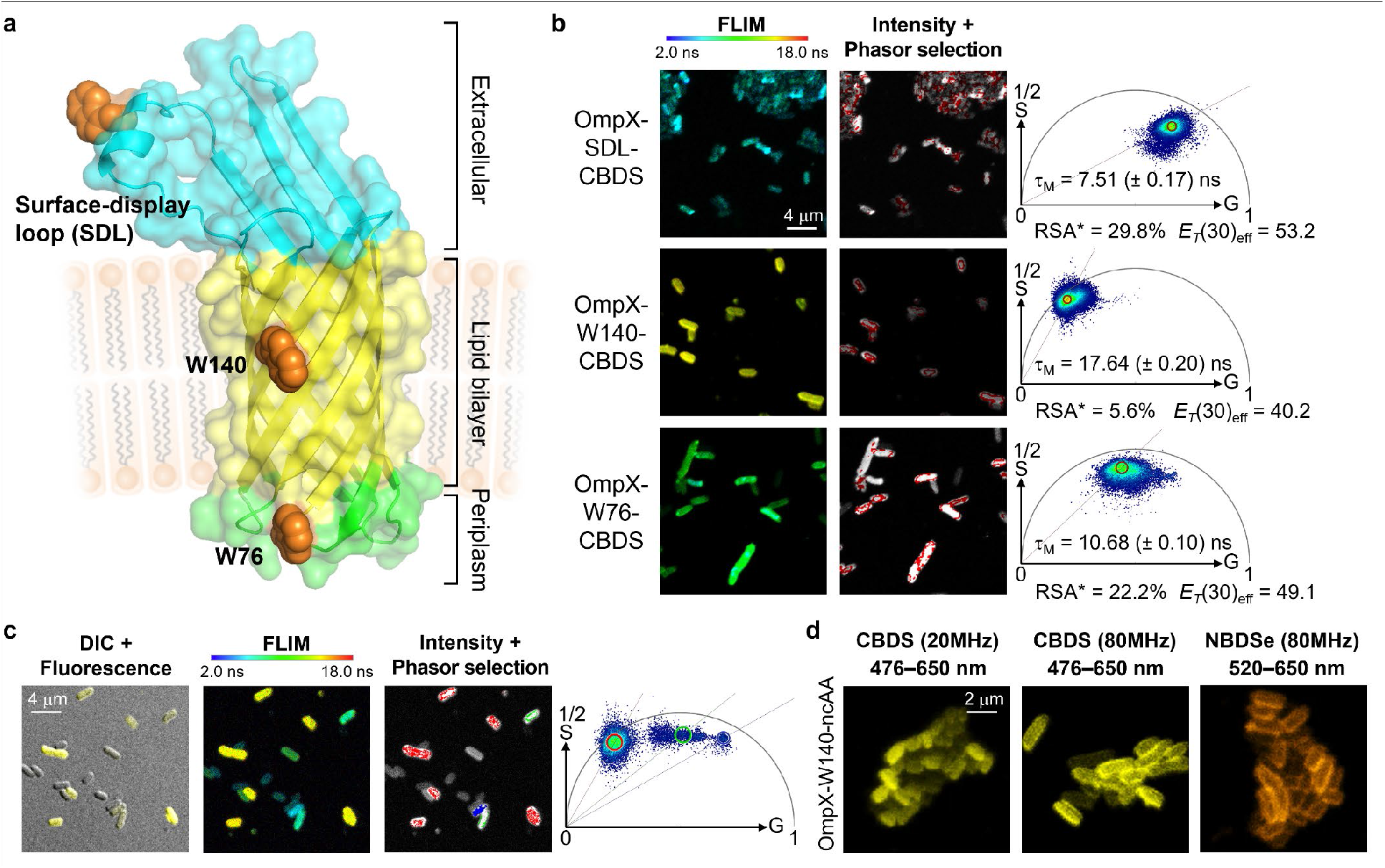
Quantitative FLIM of transmembrane hydration gradients using the genetically encoded CBDS probe. **a**, Structural model of *E. coli* OmpX embedded in a lipid bilayer, generated using AlphaFold2 based on PDB 1QJB.^51^ An artificial surface-display loop (SDL) consisting of seven alanine residues was inserted between S53 and S54. The substituted sites include the fifth alanine residue in the SDL (shown as a tryptophan side chain for space-filling representation), W140, which is exposed to the hydrophobic region of the lipid bilayer, and W76, which is located at the interface with the periplasm. **b**, FLIM analysis of *E. coli* cells expressing the respective CBDS-labeled OmpX variants. The calculated τ_M_ values of the major lifetime populations reflect the steep hydration gradients across the cell membrane. **c**, Differential interference contrast (DIC) images overlaid with the fluorescence channel, together with FLIM analysis of a co-culture of *E. coli* B95 cells expressing three different OmpX-CBDS variants. Differences in both fluorescence intensity and τ_M_ enable discrimination between the different variants. **d**, Fluorescence images of *E. coli* cells expressing OmpX-W140CBDS/NBDSe mutants acquired at laser repetition rates of 20 or 80 MHz. Imaging at 80 MHz for both CBDS and NBDSe preserved the spatial localization of the labeled membrane proteins, with magnified views reveal clear fluorescence at the bacterial cell surface.

This environmental gradient produced a pronounced change in CBDS fluorescence lifetime (Fig. 5b; Extended Data Fig. 5). Moving from the extracellular loop of OmpX (τ_M_ = 7.5 ns) to the site of W76 at the interface to the periplasm (10.7 ns) and finally to the deeply buried W140 (17.6 ns), CBDS exhibited a 2.3-fold lifetime increase. Using the empirical solvent calibration established above (Fig. 1f), the τ_M_ values can be interpreted in terms of effective local polarity (*E*_T_(30)_eff_) at each site: the extracellular loop experiences an environment between bulk-water-like and partially dehydrated (*E*_T_(30)_eff_ = 53 kcal/mol), W76 resides in a more dehydrated interfacial zone (*E*_T_(30)_eff_ = 49 kcal/mol), and W140 is shielded in a highly apolar, aprotic microenvironment (*E*_T_(30)_eff_ = 40 kcal/mol).

The lipid bilayer core has a low effective dielectric constant (ε ≈ 2–3.2), corresponding to weakly polar non-hydrogen-bonding solvents with *E*_T_(30) values of approximately 33–35 kcal/mol.^53, 54^ The *E*_T_(30)_eff_ value at W140 is substantially lower than those characteristic of aqueous environments, yet remains above the values expected for a hydrocarbon-like lipid bilayer core. This observation is consistent with the heterogeneous nature of the membrane–protein interface. The membrane interior is not simply a uniform hydrocarbon phase as the local environment of the fluorophore can be influenced by nearby protein residues, lipid functional groups, and water molecules associated with the protein-lipid interface. Previous site-directed fluorescence studies of membrane proteins using BADAN have shown that fluorophore polarity and hydration can vary substantially with membrane depth.^55^ An older fluorescence study likewise provided evidence for water at protein-lipid interfaces, including around hydrophobic protein surfaces.^56^ Thus, the *E*_T_(30)_eff_ value at the site of W140 reflects the composite microenvironment directly surrounding the CBDS side chain rather than the polarity of the bulk lipid phase. This interpretation highlights an important advantage of site-specifically encoded CBDS over measurements reporting on bulk properties.

The long lifetime of CBDS, extending to nearly 20 ns in hydrophobic environments, necessitates a reduced laser repetition rate of 20 MHz to provide sufficient temporal separation between successive excitation pulses for accurate lifetime determination. Despite the lower number of excitation cycles per unit time, CBDS retained sufficient fluorescence contrast and spatial information. Indeed, when *E. coli* cells expressing three different OmpX-CBDS variants were mixed in a single co-culture, the populations could be distinguished by differences in both fluorescence intensity and τ_M_ (Fig. 5c). Increasing the repetition rate to 80 MHz increased photon sampling per unit acquisition time and improved the spatial definition of OmpX-W140CBDS, revealing clearer membrane localization (Fig. 5d). We therefore investigated NBDSe as a complementary FLIM probe whose shorter lifetime is compatible with higher repetition-rate excitation. NBDSe-labeled OmpX showed clear membrane localization at 80 MHz laser repetition rate (500 nm excitation; Fig. 5c) and retained the same environment-dependent lifetime trend across the OmpX incorporation sites as CBDS, although over a narrower dynamic range (τ_M_ = 1.3, 2.6 and 4.0 ns; Extended Data Fig. 5c).

## Discussion

The use of quantitative TCSPC-FLIM in living cells has historically been constrained by the lack of suitable genetically encoded probes. By establishing a robust genetic code expansion platform for the site-specific incorporation of a family of BD-ncAAs into proteins, the present work successfully encoded NBDO, NBDSe, and the novel cyano-substituted CBDS, providing a suite of minimally perturbing sensors for excitation with standard visible laser lines. CBDS stands out as a genetically encodable, compact fluorophore that combines exceptionally long fluorescence lifetimes with pronounced environmental sensitivity, enabling precise and quantitative mapping of local protein microenvironments and transmembrane hydration gradients in living cells using wash-free FLIM. Beyond demonstrating eukaryotic compatibility, the yeast experiments illustrate how site-specific lifetime sensing can distinguish heterogeneous protein microenvironments encountered during protein trafficking and processing. To facilitate adoption of this technology, the plasmid required for bacterial incorporation of all three BD-ncAAs has been deposited with Addgene (Watertown, MA; #225292).

### Advantages

CBDS combines several features that are advantageous for quantitative FLIM. While existing fluorescent ncAAs demonstrate environmental sensitivity or long-lived fluorescence, CBDS combines a long fluorescence lifetime with a broad, polarity-dependent lifetime response, providing a larger dynamic range for resolving changes in the local protein environment. Its long lifetime also facilitates discrimination from short-lived cellular autofluorescence, while its excitation and emission wavelengths in the visible range are compatible with conventional confocal FLIM platforms.

With a side chain volume comparable to that of tryptophan, CBDS can replace individual aromatic residues and minimize any perturbation of the target protein and its local chemical environment. CBDS meets the three criteria of small size, visible-range fluorescence wavelengths and sensitive fluorescence lifetime changes for an ideal genetically encoded reporter for reporting changes in the local protein microenvironment by a quantitative fluorescence-lifetime readout.

The successful genetic encoding and FLIM imaging of CBDS in both bacterial and yeast cells shows that this environmentally sensitive fluorophore can readily be applied in prokaryotes and eukaryotes. As the G1-PylRS platform has also been established in mammalian cells,^48^ our results provide a strong foundation for extending CBDS-based fluorescence lifetime imaging to more complex eukaryotic systems and in vivo applications.

BD-ncAAs have shown scaffold-dependent photostability, with NBDSe exhibiting increased resistance to photobleaching relative to NBDO.^21^ Under the imaging conditions used here, CBDS maintained robust fluorescence signals during repeated raster scanning, including 80 MHz acquisitions with a ∼2.0 μs pixel dwell time, six line accumulations and acquisition periods of ∼11 s, supporting sufficient photophysical stability for quantitative fluorescence imaging. Systematic photobleaching measurements will nevertheless be required to establish its photostability under prolonged or repeated imaging.

### Limitations

High-yield protein expressions in the *E. coli* B95 ΔAΔ*fabR* strain occasionally exhibit signs of glutamine incorporation at the targeted amber codon, which can be attributed to the similarity between the respective codons (Extended Data Fig. 2a–f). This effect can be observed with any ncAA and calls for further engineering of the translation machinery.

Chemical reduction of nitro to amino groups is possible in the reducing environment of the bacterial cytosol. In *E. coli*, the reduction is likely mediated by endogenous nitroreductases. We noted the effect with NBDO, whereas the NBDSe and CBDS probes proved stable. The chemical stability of these probes may need reassessment in uncharacterized expression hosts or novel biological contexts. Similarly, potential ncAA-associated cytotoxicity should be evaluated when these probes are applied in new cell types or at substantially different concentrations or exposure times.

### Outlook

The present results invite several directions for exploration. Extending BD-ncAA incorporation to mammalian cells would broaden the biological contexts accessible to lifetime-based measurements. The ability to probe local environmental changes at defined protein sites opens opportunities for studies of dynamic processes such as post-translational modification-dependent protein association, conformational transitions, folding and allostery. The approach could detect site-specific protein interactions in soluble and membrane-associated proteins in living cells. An expanded palette of BD-ncAAs with systematically tuned photophysical properties could enable multiplexed measurements. In combination, these developments could extend the system to a general strategy for quantitatively mapping dynamic protein microenvironments in living systems.

## Methods

### Synthesis of BD-ncAAs

All chemicals and solvents were of reagent grade (>95%) and used as received unless otherwise noted. Chemicals were obtained from Ambeed or from Merck. Unless otherwise specified, reactions were carried out in oven-dried glassware, under an atmosphere of nitrogen. Please refer to the Supplementary Information for full details for synthesis of NBDO and NBDSe amino acids.

(*S*)-2-((*tert*-Butoxycarbonyl)amino)-3-((7-cyanobenzo[c][1,2,5]thiadiazol-4-yl)amino)propanoic acid (Boc-CBDS)

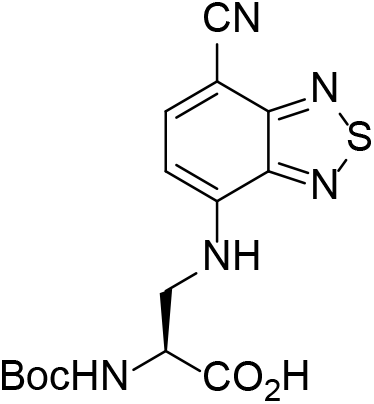

A microwave vial was charged with 4,7-dibromobenzo[*c*][1,2,5]thiadiazole (991 mg, 3.37 mmol), CuCN (301 mg, 3.36 mmol) and *N,N*-dimethylacetamide (11 mL). The resulting mixture was sparged with argon for 10 min, then heated under microwave conditions to 200 °C for 10 min. The reaction mixture was cooled, diluted with saturated aqueous NaHCO3 solution, and extracted with ethyl acetate (3x 200 mL). The organic extracts were combined, washed with brine, dried (Na_2_SO_4_), and the solvent removed *in vacuo*. The crude mixture of starting material, mono-cyanated, and di-cyanated products, was then dissolved in DMSO (11 mL). Boc-Dap-OH (274 mg, 1.34 mmol) and DIPEA (580 µL, 3.32 mmol) were added, and the resulting solution was heated to 100 °C for 17 h. The reaction mixture was cooled to room temperature, diluted with ethyl acetate, and washed with 1H aqueous HCl solution (5x 50 mL) and brine (50 mL), then dried (Na_2_SO_4_) and the solvent removed *in vacuo*. Flash column chromatography, eluting with a gradient from 100% CH_2_Cl_2_ → 15% MeOH/84% CH_2_Cl_2_/1% AcOH, afforded the desired compound as an orange-brown solid (145 mg, 12%). Mp: 180 °C (decomp.); ^1^H NMR (300 MHz, MeOD): δ 7.90 (d, *J* = 7.9 Hz, 1H), 6.60 (d, *J* = 7.9 Hz, 1H), 4.46 – 4.37 (m, 1H), 3.88 (dd, *J* = 13.8, 4.6 Hz, 1H), 3.76 – 3.67 (m, 1H), 1.38 (s, 9H); ^13^C{^1^H} NMR (MeOD, 126 MHz): δ 157.9, 155.4, 147.6, 146.4, 141.1, 118.4, 102.0, 89.5, 80.7, 55.6 (broad), 46.3, 28.7; HRMS (ESI): [M+H]^+^ calc. for C_15_H_18_N_5_O_4_S, 364.10740; found 364.10765; IR (ATIR, solid): 3341, 2928, 2211, 1698, 1562, 1158 cm^-1^.

(S)-2-Amino-3-((7-cyanobenzo[c][1,2,5]thiadiazol-4-yl)amino)propanoic acid hydrochloride (CBDS·HCl)

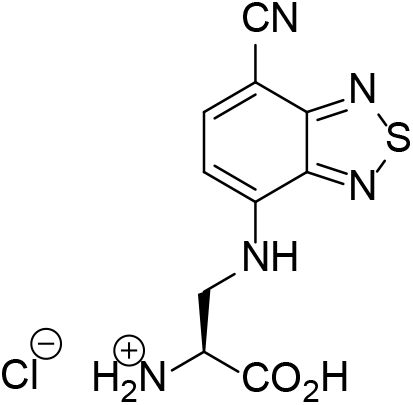

HCl in dioxane (1.7 mL, 4M solution) was added to a solution of Boc-CBDS (135 mg, 0.371 mmol) in anhydrous THF (4 mL) at room temperature. The resulting solution was heated at reflux for 2.5 h. The solvent was removed under a flow of nitrogen, and the resulting solid triturated with dichloromethane to afford the desired compound as an orange-brown solid (110 mg, *quant.*). m.p. 170°C (decomp.); ^1^H NMR (500 MHz, DMSO-*d6*): δ 8.09 (d, *J* = 8.2 Hz, 1H), 7.95 (t, *J* = 5.8 Hz, 1H), 6.69 (d, *J* = 8.2 Hz, 1H), 4.10 – 4.05 (m, 1H), 3.97 – 3.83; ^13^C{^1^H} NMR (126 MHz, DMSO-*d6*): δ 169.0, 153.8, 145.9, 144.3, 139.9, 117.5, 101.5, 88.1, 51.5, 42.8; HRMS (ESI): [M+Na]^+^ calc. for C_10_H_9_N_5_NaO_2_S, 286.03692; found 286.03658; IR (ATIR, solid): 2846 (broad), 2210, 1728, 1557, 879, 514 cm^-1^.

### Screening of functional G1PylRS for BD-ncAAs

The identification of functional G1PylRS enzymes to recognize BD-ncAAs used a previously established library of G1PylRS mutants encoded on the pBK-G1RS plasmid.^15^ The plasmid library was transformed into *E. coli* DH10B cells harboring the pBAD-H6RFP reporter plasmid encoding mRFP1 with an amber stop codon following an N-terminal His_6_ tag. After transformation, the culture was directly inoculated into 25 mL LB medium supplemented with 100 mg/L carbenicillin, 50 mg/L kanamycin, 0.4% L-arabinose, and 0.5 mM target ncAA. This culture served as the sample for the first round of positive selection (1P+). The same cells grown without ncAA supplementation were used as a control sample (1P–). Overnight expression at 37 °C led to readily detectable level of RFP expression, and the cells were then harvested, resuspended in 5 mL of PBS buffer (137 mM NaCl, 2.7 mM KCl, 10 mM Na₂HPO₄, 1.8 mM KH₂PO₄, pH 7.4), and diluted 20-fold. FACS was performed using a FACSAria Fusion cell sorter (BD Biosciences, USA). Cells with high red fluorescence levels were collected from the 1P+ sample (indicated by violet shades in Supplementary Fig. 4, 5) and subjected to a subsequent round of negative selection (2N–) in the absence of ncAA. Cells exhibiting low RFP expression were collected from the 2N– sample and aliquoted to inoculate media under positive (3P+, with ncAA) and negative (3P–, without ncAA) conditions. The selection experiments continued with iterative positive and negative rounds following the same strategy. The 5P+ sample for NBDO demonstrated a clear response to the presence of ncAA, and the top 3.3% of RFP-fluorescent cells from the 5P+ sample were collected. The selection experiments for NBDSe continued to the seventh round to reach enrichment on the target population. The top 3% of RFP-fluorescent cells from the 7P+ samples were collected.

About 2,000 cells collected from each final round were recovered by plating on LB agar plates containing 100 mg/L carbenicillin and 50 mg/L kanamycin. Isolated colonies were analyzed in 96-well plates. Sixty enzyme candidates for each selection experiment were inoculated into media under both positive (with 0.5 mM ncAA) and negative (without ncAA) conditions. The red fluorescence intensity was measured as an indicator of RFP expression and normalized to the OD_600_ of the cell culture using a TECAN Infinite 200 Pro M Plex plate reader (Tecan, Switzerland; Supplementary Fig. 6, 7). DNA sequence analysis of the pBK-G1RS plasmids identified four and two individually different candidates to incorporate NBDO and NBDSe, respectively. The amino acid mutations of these candidates are listed in Supplementary Table 3.

To assess the substrate cross-specificity, the tRNA synthetase plasmids pBK-G1RS carrying each of the six selected RS mutants were individually co-transformed with the reporter plasmid pBAD-H6RFP into *E. coli* DH10B cells. Cells were grown in LB medium containing 100 mg/L carbenicillin and 50 mg/L kanamycin at 37 °C. 5 μL of each overnight culture were used to inoculate 150 μL LB medium grown on 96-well plates, supplemented with the same antibiotic, along with 0.4% L-arabinose under four different conditions of ncAA supplementation: without any ncAA; with 0.5 mM of NBDO, NBDSe, or CBDS. Each culture was prepared in triplicate. The plates were shaken at 400 rpm for 12 h at 37 °C and for 24 h at 25 °C. The average of the triplicates was calculated according to the measured intensity of red fluorescence and normalized by the OD_600_ of the cell culture using a TECAN Infinite 200 Pro M Plex plate reader (Tecan, Switzerland). Based on the cross-specific reactivity, G1NBDO53 was subsequently used to incorporate NBDO and CBDS, and G1NBDSe09 was used to incorporate NBDSe.

### BD-ncAA incorporation in *E. coli*

For preparative protein production or imaging in *E. coli*, the G1NBDO53 and G1NBDSe09 synthetase genes were subcloned into pRSF vector.^57^ *E. coli* B-95.ΔAΔ*fabR* cells were co-transformed with the synthetase plasmid pRSF-G1NBDO53/G1NBDSe09 and the pCDF plasmid containing the reporter gene, where the target residue was replaced by an amber codon (Twist Bioscience, USA). The transformed cells were grown at 37 °C in LB medium containing 25 mg/L kanamycin and 25 mg/L spectinomycin. A 1 mL aliquot of the overnight culture was used to inoculate 100 mL LB medium supplemented with 25 mg/L kanamycin, 25 mg/L spectinomycin. Cells were grown at 37 °C to an OD_600_ of 0.4, at which point the corresponding BD-ncAA was added to a final concentration of 0.5 mM. The culture was then continued at 37 °C until an OD_600_ of 0.6 was reached. The temperature was subsequently reduced to 25 °C, and protein expression was induced by the addition of IPTG a final concentration of 1mM.

For protein purification, after expression for 16 h, the cells were harvested by centrifugation (4,000 *g*, 4 °C, 15 min). Following resuspension in buffer A (50 mM Tris-HCl, pH 7.5, 300 mM NaCl, 5% glycerol, 10 mM imidazole), the cells were lysed by sonication as described above. The cell lysate was centrifuged (30,000 *g*, 4 °C, 1 h) and the cleared supernatant was loaded onto a 1 mL His GraviTrap column (Cytiva, USA). The column was washed with 30 column volumes buffer A and the protein was eluted with 5 column volumes buffer B (same as buffer A but with 500 mM imidazole). Afterwards, the buffer was exchanged to PBS buffer using an Amicon ultrafiltration centrifugal tube (Merck Millipore, Germany) with a molecular weight cut-off of 10 kDa.

### BD-ncAA incorporation in *S. cerevisiae*

For the construction of *S. cerevisiae* capable of genetic code expansion for BD-ncAA incorporation, BY4741 was used as the parental strain. The G1NBDO53 synthetase/tRNA expression cassette was assembled using the Yeast Toolkit MoClo system. Briefly, the G1NBDO53 gene was cloned under control of the constitutive TDH3 promoter and ENO1 terminator, together with the cognate G1-tRNA^Pyl^ expressed from a hybrid SCR1′-tRNA^Gly^ dicistronic RNA polymerase III promoter and SUP4 terminator.^58^ The complete expression cassette was assembled into the pYTK096 integration vector, which contains homology arms for genome integration at the URA3 locus.

Reporter constructs were maintained on a centromeric low-copy number CEN6/ARS4 plasmid carrying LEU2 as the selectable marker. Reporter expression was controlled by the GAL1 promoter and SSA1 terminator. For assessment of amber suppression efficiency, the reporter encoded an Electra2-mScarlet3 fusion separated by a flexible linker containing a TAG amber stop codon.

For yeast surface-display experiments, the reporter construct encoded an N-terminal MFα pre-pro secretion signal (MFapp8),^59^ followed by H11-W37TAG/H11-Nt-TAG, and a linker to the C-terminal anchoring region of Sed1p for cell-surface display.^50^

The G1NBDO53/tRNA integration cassette was linearized and co-transformed with the appropriate reporter plasmid into competent BY4741 cells. Transformed cells were plated on SD-SCAA agar lacking, tryptophan, leucine and uracil at 30 °C for 2-4 days. Individual colonies were subsequently streaked onto new selective agar plates.

For flow cytometry and fluorescence microscopy experiments, three independent colonies were inoculated into 500 μL SD-SCAA medium lacking tryptophan, leucine and uracil in 96-deep-well plates and grown to saturation for 1-2 days at 30 °C. Cultures were diluted 1:5 into 500 μL SG-SCAA induction medium, to a starting OD_600_ of ∼0.3-0.5. Cells were supplemented with 1 mM BD-ncAA or no ncAA and incubated for 20 h at 30 °C. Cells were harvested by centrifugation (1,000 *g*, 4 °C, 5 min), washed twice with PBS and resuspended in PBS. For fluorescence microscopy, cells were immobilized on 1% (w/v) agarose pads.

SD-SCAA propagation medium contained 2% (w/v) glucose, 0.67% (w/v) yeast nitrogen base without amino acids, 0.146% (w/v) yeast synthetic dropout medium lacking uracil, leucine and tryptophan. SG-SCAA induction medium contained 2% (w/v) galactose, 0.67% (w/v) yeast nitrogen base without amino acids), 0.146% (w/v) yeast synthetic dropout medium lacking uracil, leucine and tryptophan, 0.54% (w/v) anhydrous Na₂HPO₄ and 0.856% (w/v) NaH_2_PO_4_, pH 6.

### Spectroscopic measurement

Solvents used in all photophysical measurements were of spectroscopic grade, and obtained from Merck.

For fluorescence spectroscopy, both the free BD-ncAAs and protein samples were prepared at a concentration of 10 μM. The fluorescence data of Fig. 1d, Fig. 2e–g, and Extended Data Fig. 2g were obtained using a CARY Eclipse fluorescence spectrophotometer (Agilent, USA) with a 3 mL/400 μL quartz cuvette. PBS buffer was used as the blank. The scan rate was 600 nm/min with 1 nm of data interval using the photomultiplier tube (PMT) detector voltage at 600 V. The absorbance data shown in Fig. 1d and Extended Data Fig. 1b and 1c were recorded using a Cary 60 UV–Vis spectrophotometer (Agilent, USA).

Data in Fig. 1e, 1f, 3f, and Extended Data Fig. 1e were collected using a Horiba Fluorolog-QM^TM^ fluorometer with a time-correlated single photon counting (TCSPC) configuration. Samples for solvent screen experiments were prepared from a 5 mM stock solution in DMSO, and diluted to 5 μM concentration with the specified solvent. Protein samples were prepared in PBS buffer, and diluted to 1 – 2 μM concentration of CBDS. An excess of ArcNL was added as a solution in PBS buffer (50 μM). All samples were excited at 405 nm using a DeltaDiode^TM^ laser (Horiba Scientific). Emission at the wavelength of maximum intensity was collected until accumulation of 5,000 counts. The instrument response function (IRF) was collected by measuring the light scattered at 405 nm from a LUDOX^®^ suspension. Lifetimes were calculated by deconvolution of the IRF from the fluorescence decay curve, followed by fitting to a single exponential function:

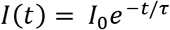

Fitting was performed using the PowerFit-10 module within FelixFL software (Horiba Scientific). The quality of each fit was assessed by chi-squared value (χ^2^) and the residual between the raw data and its fit.

### Intact protein mass spectrometry

Intact protein analysis was performed on an Orbitrap Fusion Tribrid mass spectrometer (Thermo Fisher Scientific, USA) connected to a Thermo Fisher Scientific UltiMate 3000 HPLC system equipped with ZORBAX 300SB-C3, 3.5 μm, 4.6 x 50 mm HPLC column (Agilent Technologies, USA). Approximately 50 pmol of sample was injected using a 500 μL/min linear gradient of solvent A (0.1% (v/v) formic acid in water) and solvent B (0.1% (v/v) formic acid in acetonitrile), ramping solvent B from 5% solvent B at the start to 80% after 12 min. Data were collected using an electrospray ionisation (ESI) source in positive ion mode. Protein intact mass was determined by deconvolution using the program Xcalibur 3.0.63 (Thermo Fisher Scientific, USA).

### Protein crystallization and structure determination

ArcNL was loaded onto a HiLoad 26/600 Superdex 75 pg column equilibrated with PBS. The fractions containing ArcNL were identified using SDS-PAGE and concentrated to a stock solution using a 3 kDa MWCO Amicon ultrafiltration centrifugal tube (Merck Millipore, USA). Stock solutions of H11-W37NBDO and H11-W37NBDSe were created in an identical manner. Equimolar amounts of ArcNL and either H11-W37NBDO or H11-W37NBDSe were combined and then loaded onto the same gel filtration column. The monodisperse peak of the heterodimer was confirmed using SDS-PAGE, and samples were concentrated to a stock solution of 10 mg/mL using a 3 kDa MWCO filter. Heterodimer solutions were then diluted to 5 mg/mL, 2.5 mg/mL and 1.25 mg/mL, and, using vapor diffusion and 2 μL drops of protein:mother liquid on conditions based on the Hampton pre-crystallization test. Clusters of small crystals were identified in condition B1 (0.1 M Tris-HCl, pH 8.5, 1 M ammonium sulfate) for all protein concentrations. After iterative rounds of seeding and screening gradients of ammonium sulfate, single crystals of sufficient size were grown for both samples at 1.25 mg/mL in 2 M ammonium sulfate, 0.1 M Tris-HCl, pH 8.5.

Diffraction data were collected at 100 K at the MX2 beamline at the Australian Synchrotron^60^ and processed using XDS space group.^61^ Data were scaled in Aimless (CCP4^62^) until the data was of sufficient quality in the highest resolution shell. The structure was solved using molecular replacement (Phaser MR^63^ using PDB ID 8QF4^43^ as the search model, followed by remodeling in Coot,^64^ and refinement in Phenix.^65^

CIF restraints for the chromophore were generated by building the molecule in PyMOL (The PyMOL Molecular Graphics System, Version 3.0 Shrödinger, LLC.), generating hydrogens, followed by processing of the resulting PDB file in eLBOW.^66^ The resulting CIF dictionary was then modelled in the structure using the replace residue function in Coot and used as a restraint for refinement. Data collection and refinement statistics are given in Supplementary Table 6.

### Affinity measurements by BLI

Binding kinetics of nanobody H11 mutants with Twin-Strep-tagged ArcNL were measured on a Gator Pivot (Gator Bio) at 30 °C with Strep-Tactin XT probes (Gator Bio). Probes were pre-equilibrated in K-buffer (Gator Bio) prior to loading ArcNL (5 μg/mL) to a response of 2 nm, followed a 60 second baseline stabilization step in K-buffer. The association with H11 was measured by dipping the sensors into wells containing H11 at concentrations of 20–0.625 nM in a two-fold dilution series for 180 seconds, followed by transfer into buffer-only wells for 180 seconds to measure the dissociation. Reference probes (ligand-loaded in buffer-only and unloaded probes dipped into analyte at the highest concentration) were included to account for baseline drift and background subtraction. Data were reference-subtracted, aligned and globally fitted to a 1:1 binding model to determine association (*k*_on_) and dissociation (*k*_off_) rate constants, as well as the equilibrium dissociation constant (*K*_D_). The reported kinetic parameters represent the mean of two independent experiments.

### Protein thermal stability measurements

The stabilities of the protein folds of H11 mutants were assessed by thermal denaturation monitored by circular dichroism (CD) spectroscopy using a Chirascan spectrophotometer (Applied Photophysics, UK) and a 1 mm quartz cuvette. The CD signal was monitored at 218 nm over a temperature range of 24 to 95 °C. In all cases, the protein concentration was 10 μM in PBS and measurements were recorded for 4 s per point in 1 °C increments. Blanks containing PBS measured in the same cuvette were subtracted from the data. As previous work attributed the atypical ellipticity of H11 above 210 nm to aromatic–disulfide interactions, we used TCEP-treated wild-type H11 as a reference control; this treatment did not impair H11 binding.^43^

### Imaging setup

For microscopy, *E. coli* B-95.ΔAΔ*fabR* cells or genome modified *S. cerevisiae* BY4741 were cultured as described above. Control cells lacking the BD-ncAA genetic encoding system were cultured with the same concentration of ncAA to assess background association of free BD-ncAAs. Cells were isolated from the medium by centrifugation at 4,000 × *g* for 2 min. The pellet was resuspended in PBS and incubated for 5 min, followed by centrifugation to remove residual medium. 10 μL of the resuspended cells in PBS (threefold diluted relative to the overnight culture) were then loaded onto 1% agarose pads and covered with coverslips.^67^

Imaging was performed using a Stellaris 8 confocal microscope (Leica, Germany), equipped with a 63×/1.40 oil-immersion objective. Fluorescence was detected using a HyD X detector in photon-counting mode. Images were acquired with a 1 Airy unit pinhole, a scan speed of 400 Hz and a pixel dwell time of approximately 2.01 μs. Laser excitation power was adjusted between samples to obtain sufficient photon counts while avoiding detector saturation, unless otherwise stated.

For CBDS, fluorescence was excited at 440 nm and emission was collected between 476 and 650 nm. FLIM measurements were normally acquired using a laser repetition rate of 20 MHz to accommodate the long fluorescence lifetime of CBDS. Where indicated, CBDS was additionally imaged at 80 MHz for comparison of fluorescence intensity and spatial resolution under higher-repetition-rate acquisition conditions. For NBDSe, fluorescence was excited at 500 nm and emission was collected between 520 and 650 nm using a laser repetition rate of 80 MHz.

FLIM data were processed in LAS X using phasor analysis with the first harmonic, a phasor threshold of 15, pixel binning of 1, and the wavelet filter enabled.^68^ Analysis was performed over the entire acquired field without applying additional fluorescence-intensity gating or temporal gating to the TCSPC data. For each region of interest (ROI), the fluorescence decay was transformed into its phasor coordinates, G and S, corresponding to the cosine and sine components of the normalized Fourier transform of the fluorescence decay, respectively. The phasor modulation was calculated as:

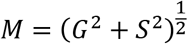

and the modulation lifetime (τ_M_) was calculated according to:

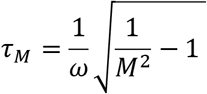

where ω = 2πf and f is the laser repetition frequency used for the corresponding acquisition. For each imaging experiment, τ_M_ was calculated from the ROI-averaged phasor coordinates G and S, rather than by averaging pixel-wise lifetime values. All measurements were performed in three independent imaging experiments, and the reported τ_M_ values represent the mean ± s.d. of these experiments (n = 3).

## Acknowledgments

We thank Dr Harpreet Vohra, Mr Michael Devoy, and Dr Fei-Ju Li at the John Curtin School of Medical Research, Australian National University for technical support on FACS experiments. We acknowledge Microscopy Australia (ROR: 042mm0k03) at the Centre for Advanced Microscopy of the Australian National University, especially Dr Angus Rae and Daryl Webb for their knowledge and input. We acknowledge the Australian National University Nanobody Facility and Dr. Megan Outram for assistance with BLI experiments. Financial support by the Australian Research Council, including the Centre of Excellence for Innovations in Peptide and Protein Science (CE200100012) and Discovery Projects (DP230100079, DP260100191), is gratefully acknowledged. N.P. thanks the CSIRO Advanced Engineering Biology Future Science Platform for scholarship support.

## Author contributions

H. Q. developed the genetic code expansion systems for BD-ncAAs, performed protein expression, spectroscopy, imaging, and flow cytometry analysis, designed the study, supervised the research, and wrote the manuscript. S. B. synthesized the BD-ncAAs, performed fluorescence spectroscopy, designed the study, supervised the research, and wrote the manuscript. E. H. performed X-ray crystallography and wrote the manuscript. N. P. performed protein expression, sample preparation for crystallography, and BLI and CD measurements. E. A. performed fluorescence spectroscopy and designed constructs for imaging. R. W. established the incorporation system of BD-ncAAs in yeast. X. Z. contributed to the development of genetic code expansion systems. R. M. contributed to the synthesis of BD-ncAAs. K. J., G. O., and T. H. designed the study, supervised the research, and wrote the manuscript.

---

**Extended Data Fig. 1.**
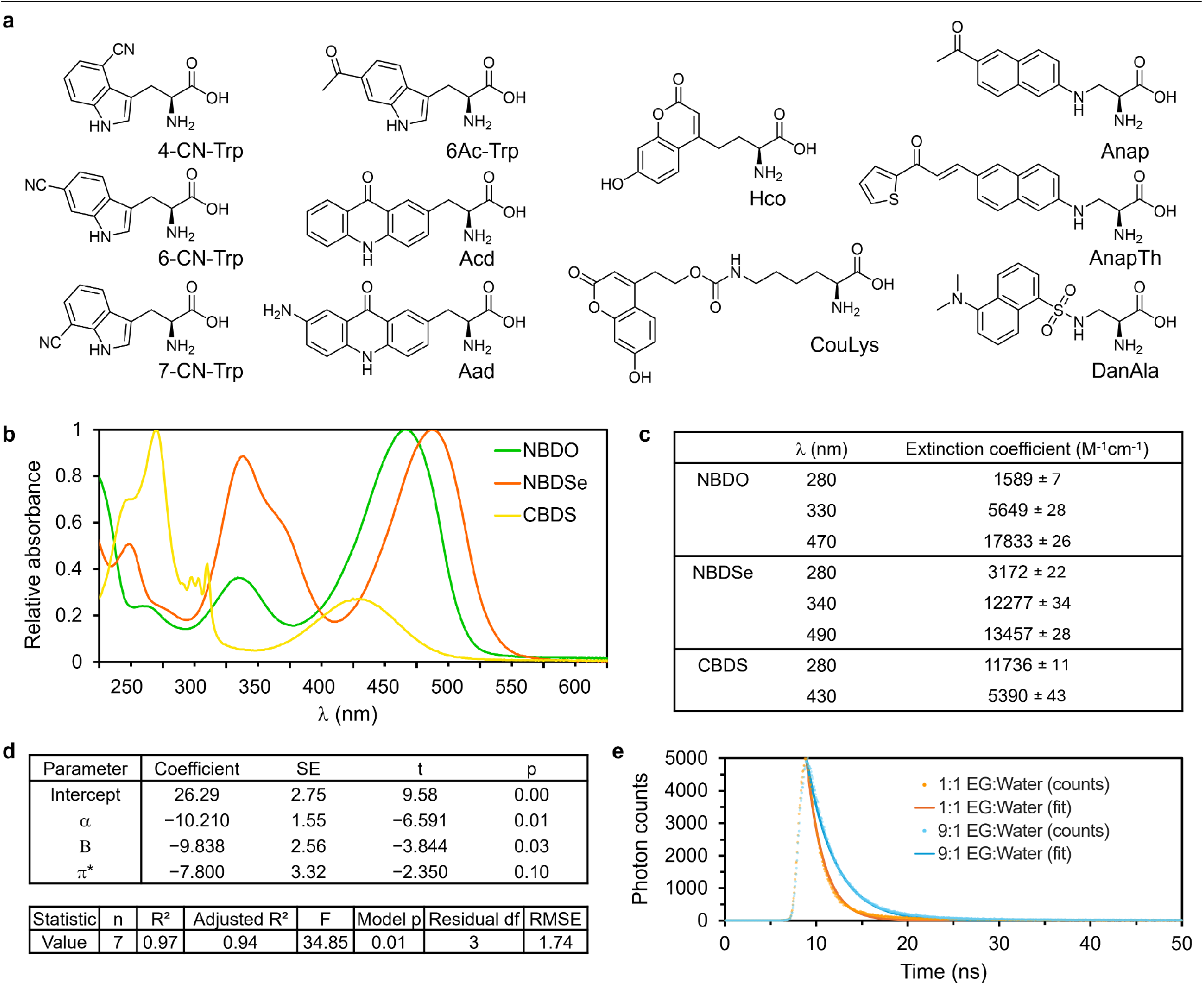
**a**, Chemical structures of the previously reported fluorescent ncAAs. **b**, **c**, Full UV-Vis absorbance spectra of NBDO/NBDSe/CBDS, with measured extinction coefficients noted for each peak. **d**, Multiple linear regression analysis of the solvent dependence of CBDS fluorescence lifetime. CBDS fluorescence lifetimes measured in seven solvents were fitted to the Kamlet–Taft solvent parameters α, β and π* using the model τ = 26.29 − 10.21α − 9.84β − 7.80π*. Regression coefficients, standard errors, t-values and two-sided p-values are shown together with overall model statistics. **e**, Fluorescence decay histograms (dots) and corresponding exponential fits (lines) for CBDS in high-viscosity [9:1] and low-viscosity [1:1] ethylene glycol (EG) to water mixtures. The decay was recorded using TCSPC and fitted using a single-exponential decay model convoluted with the instrument response function (IRF) measured under identical experimental conditions.

**Extended Data Fig. 2.**
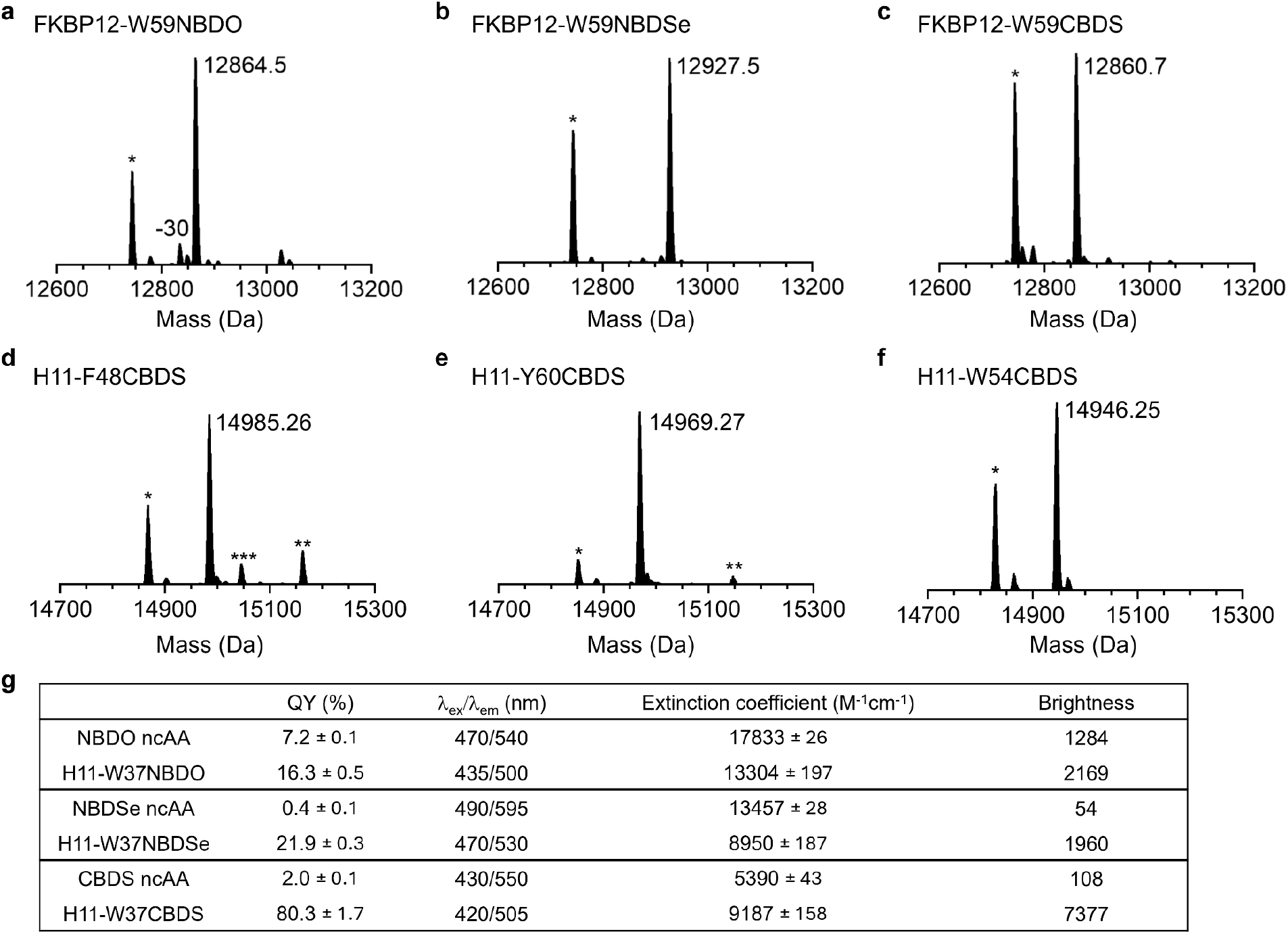
**a, b, c**, Intact protein mass spectrometry of purified FKBP12 variants confirms incorporation of BD-ncAAs at the amber-mutated Trp59 position. Major peaks match the expected masses of full-length proteins containing the corresponding ncAA without detectable modifications. Peaks marked with one asterisk (*) indicate amber stop codon suppression by glutamyl-tRNA which is commonly observed when using B95.ΔAΔ*fabR* strains to implement genetic code expansion. **d**, **e**, **f**, Intact protein mass spectrometry of purified H11 mutants with incorporated. Major peaks correspond to the expected masses. * indicates glutamyl-tRNA suppression, ** indicates the major product with α-N-gluconoylation, and *** indicates glutamine suppression with α-N-gluconoylation. **g**, Comparing the photophysical properties of the BD-ncAAs as free amino acids or incorporated into H11. Significantly enhanced fluorescence intensity was observed when NBDSe and CBDS are used to replace the core position W37 of H11. Quantum yield (QY), excitation and emission maxima, molar extinction coefficients and calculated brightness values are shown.

**Extended Data Fig. 3.**
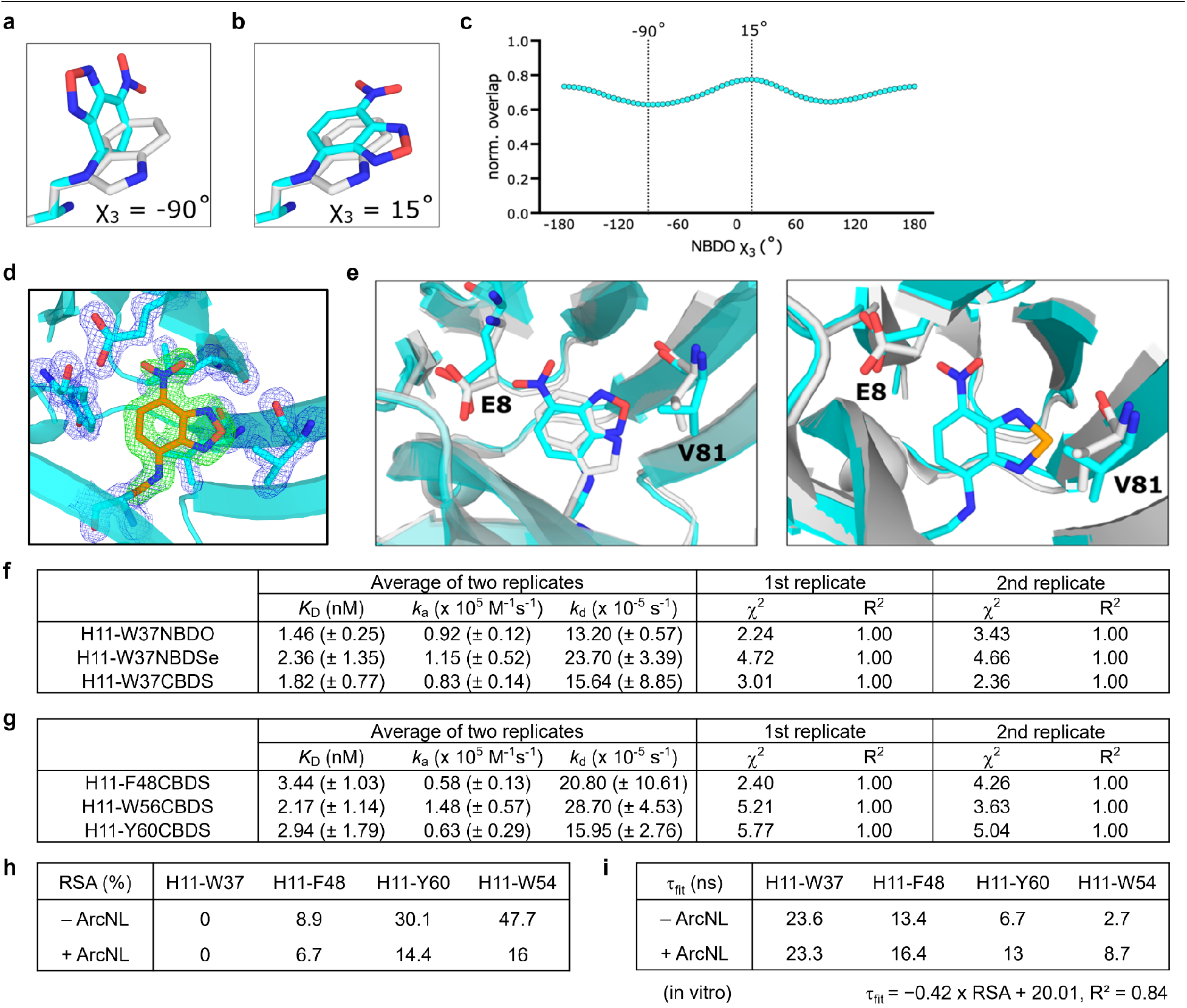
**a, b**, Theoretical overlap between Trp (white) and NBDO (cyan). Stick illustrations are set to match dihedral angles of NBDO (with χ_3_ = -90°/15°). **c**, Normalized overlap calculated by using the overlap command in PyMOL for the superimposed amino acids, for every 5° χ_3_ angle in NBDO, and dividing the result by the self-overlap of Trp. **d**, Omit electron density for the NBDO side chain is shown as a green mesh contoured at 5σ; 2mF_o_-DF_c_ density for surrounding residues is shown as a blue mesh contoured at 2σ. **e**, Environment surrounding BD-AAs in H11 with NBDO/NBDSe replacing position W37. **f**, **g**, BLI kinetic and affinity data from two independent experiments showing the mean and range of dissociation constants, on-rates and off-rates of H11 mutants binding to Arc-NL. **h**, Relative surface area (RSA) of each labelled residue calculated from the crystal structures (PDB 8QF3 and 8QF4). **i**, Correlation between the in vitro τ_fit_ and the corresponding RSA values for the same H11-CBDS mutants in the absence and presence of ArcNL.

**Extended Data Fig. 4.**
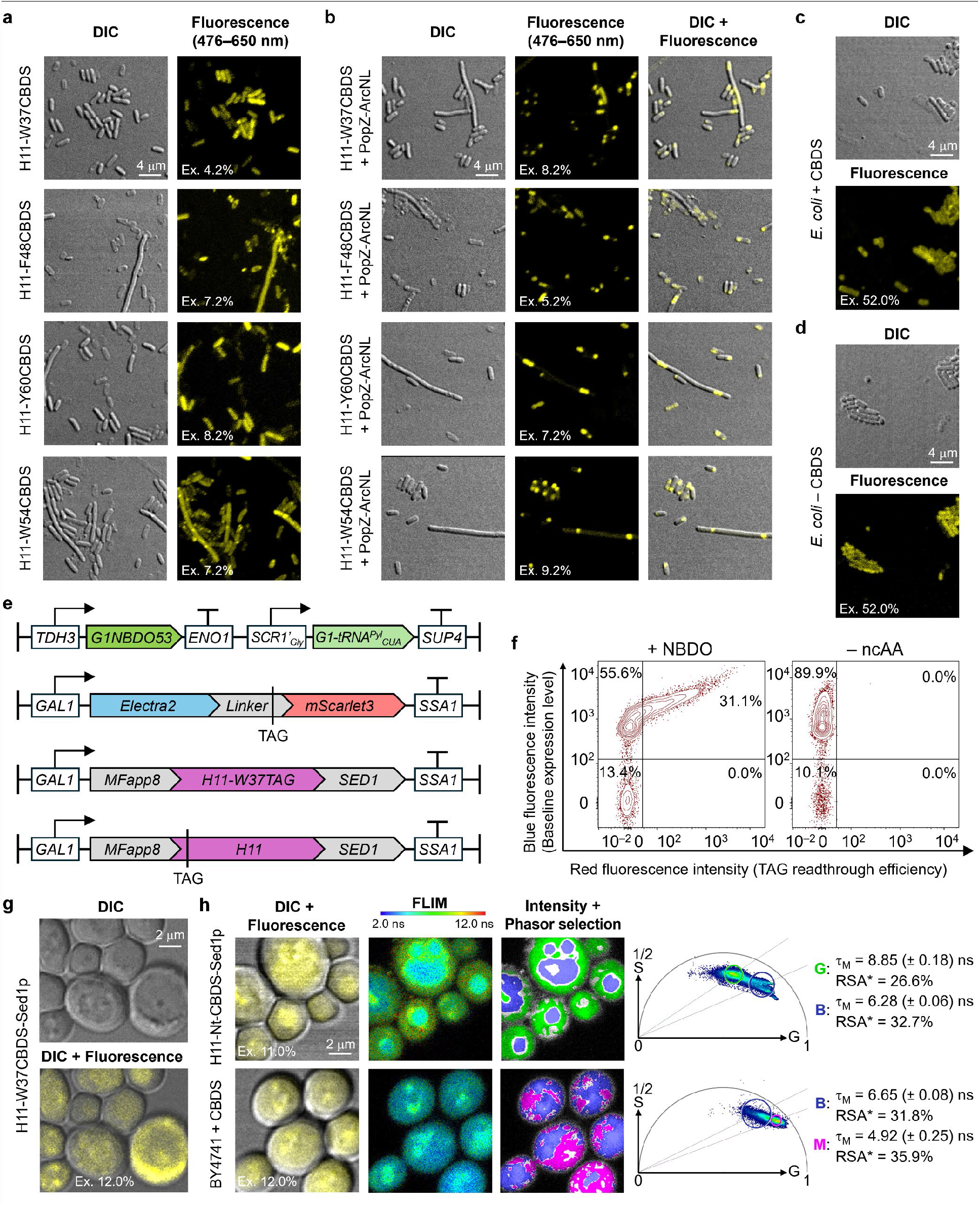
**a**, Corresponding DIC and fluorescence images of the same fields of view used for the FLIM analysis of *E. coli* cells expressing H11-CBDS mutants in Fig. 4a. Transmitted-light images were acquired using differential interference contrast (DIC), and fluorescence images were acquired with 440 nm excitation at a laser repetition rate of 20 MHz, with the excitation power adjusted for each sample to maximize photon counts without saturating the FLIM detector. Fluorescence images were acquired using the excitation powers indicated on each image. For display, the fluorescence intensity scale was adjusted by setting the upper limit of the intensity scale bar to the maximum pixel intensity in each image and were not used for quantitative intensity comparison. **b**, Corresponding DIC and fluorescence images of the same fields of view used for the FLIM analysis of *E. coli* cells expressing H11-CBDS mutants together with PopZ-ArcNL in Fig. 4b. Overlay images show the cellular localization of CBDS-incorporated H11. **c**, **d**, Corresponding DIC and fluorescence images of the same fields of view used as control for the FLIM analysis in Fig. 4c. **e**, Schematic of the yeast expression system used to incorporate BD-ncAAs and for surface display of H11-CBDS. The genomically integrated G1NBDO53RS/G1-tRNA expression cassette is driven by the *TDH3* and *SCR1′* promoters, respectively. The fluorescent reporter construct for monitoring amber suppression encodes a fusion protein comprising Electra2 and mScarlet3, separated by a flexible linker containing an in-frame TAG stop codon. The surface-display reporter construct comprises an MFα pre-pro-app8 secretion signal, H11-W37TAG/H11-Nt-TAG, and the *SED1* gene encoding the Sed1 GPI anchor protein (Sed1p) under the control of the *GAL1* promoter. **f**, Flow cytometry analysis of the readthrough assay demonstrating NBDO incorporation in *S. cerevisiae*. Translation efficiency was calculated as the fraction of cells exhibiting successful amber suppression, identified by mScarlet3 (red) fluorescence, relative to the total number of cells expressing the baseline reporter, identified by Electra2 (blue) fluorescence. **g**, Corresponding DIC images and fluorescence overlay of the same fields of view used for the FLIM analysis of *S. cerevisiae* cells expressing H11-W37CBDS-Sed1p in Fig. 4e. **h**, DIC/fluorescence overlay and FLIM analysis of *S. cerevisiae* cells expressing H11-Nt-CBDS-Sed1p or cultured with CBDS alone as a control. FLIM images are pseudocoloured by fluorescence lifetime. Phasor populations corresponding to representative cellular contexts were selected directly on the phasor plots and mapped back to the images, where the corresponding pixels are highlighted in blue, green or magenta on grayscale images. τ_M_ for each cellular context was determined independently for each image and averaged across three independent imaging experiments per condition.

**Extended Data Fig. 5.**
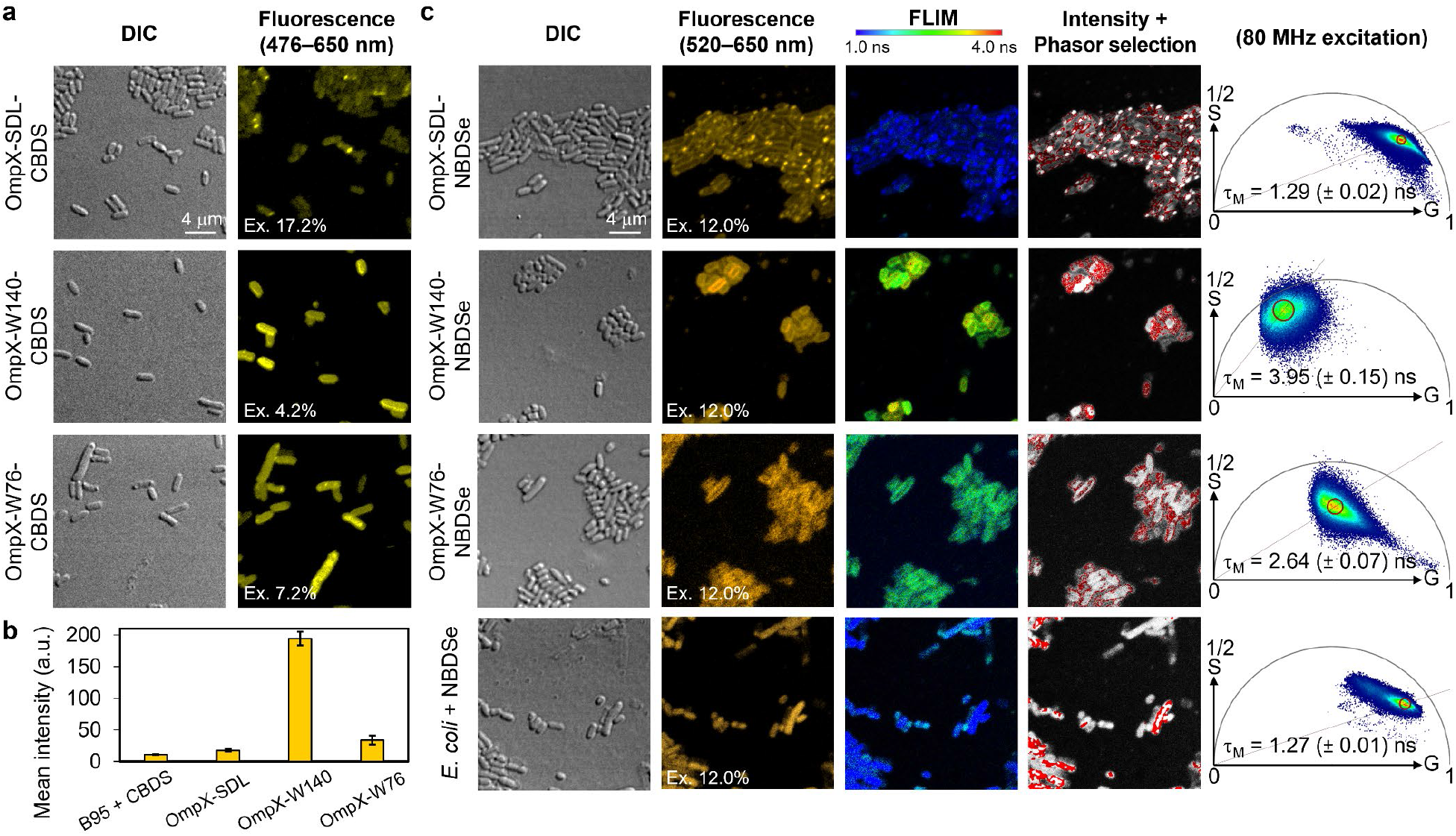
**a**, Corresponding DIC and fluorescence images of the same fields of view used for the FLIM analysis of *E. coli* cells expressing OmpX-CBDS mutants in Fig. 5b. **b**, Fluorescence intensity comparison of E. coli cells expressing three OmpX-CBDS mutants or cultured with CBDS alone, imaged at an 80 MHz laser repetition rate with the excitation power fixed at 9% for all acquisitions (corresponding images in Supplementary Fig. 21). Raw 16-bit fluorescence images were analyzed in Fiji (ImageJ) using an identical intensity threshold for all images. Pixels with intensity values ≥9 were selected, and the mean gray value of the threshold-positive region was measured from the original image. Three independently acquired images were analyzed per condition, and data are presented as mean ± SD. **c**, DIC and fluorescence images and FLIM analysis of *E. coli* cells expressing OmpX-CBDS mutants, acquired at a laser repetition rate of 80 MHz to resolve the shorter fluorescence lifetime of NBDSe. Mean τ_M_ values for each condition showed the same environment-dependent trend as the corresponding OmpX-CBDS mutants, although over a narrower lifetime range.

