## Supplementary Information for "Quantitative fluorescence lifetime imaging of protein microenvironments by genetically encoded compact benzodiazole probes"

**Table of Contents**

**Synthesis**

**Supplementary figures**

Supplementary Fig. 1. BD-ncAAs break the chromophore size–wavelength trade-off.

Supplementary Fig. 2. Fluorescence decay analysis of CBDS in different solvents.

Supplementary Fig. 3. Plot of CBDS fluorescence lifetime versus solvent dielectric constant.

Supplementary Fig. 4. FACS experiments for selection of active and specific G1PylRS enzymes that recognize NBDO.

Supplementary Fig. 5. FACS experiments for selection of active and specific G1PylRS enzymes that recognize NBDSe.

Supplementary Fig. 6. Representative FACS setup for the selection of *E. coli* cells containing the G1RS library.

Supplementary Fig. 7. Activity and specificity screen of G1PylRS variants for NBDO.

Supplementary Fig. 8. Activity and specificity screen of G1PylRS variants for NBDSe.

Supplementary Fig. 9. Two-dimensional excitation–emission spectra of free NBDO and NBDO-incorporated H11.

Supplementary Fig. 10. Two-dimensional excitation–emission spectra of free NBDSe and NBDSe-incorporated H11.

Supplementary Fig. 11. Two-dimensional excitation–emission spectra of free CBDS and CBDS-incorporated H11.

Supplementary Fig. 12. Fluorescence decay analysis of CBDS-containing H11 mutants.

Supplementary Fig. 13. FLIM analysis of *E. coli* B95 cells cultured with or without CBDS.

Supplementary Fig. 14. FLIM analysis of *E. coli* B95 cells expressing H11-CBDS mutants.

Supplementary Fig. 15. FLIM analysis of *E. coli* B95 cells co-expressing H11-CBDS mutants and PopZ-tagged ArcNL.

Supplementary Fig. 16. Relationship between RSA and $\tau_M$ of H11-CBDS mutants in *E. coli*.

Supplementary Fig. 17. FACS setup for amber readthrough efficiency test in *S. cerevisiae*.

Supplementary Fig. 18. FLIM analysis of *S. cerevisiae* BY4741 cells expressing CBDS-incorporated proteins.

Supplementary Fig. 19. FLIM analysis of *E. coli* B95 cells expressing OmpX-CBDS mutants.

Supplementary Fig. 20. FLIM analysis of *E. coli* B95 cells expressing three different OmpX-CBDS mutants in co-culture.

Supplementary Fig. 21. FLIM analysis of *E. coli* B95 cells expressing OmpX-CBDS mutants using an 80 MHz laser repetition rate.

Supplementary Fig. 22. FLIM analysis of *E. coli* B95 cells expressing H11-W37NBDSe using an 80 MHz laser repetition rate.

Supplementary Fig. 23. FLIM analysis of *E. coli* B95 cells expressing OmpX-NBDSe mutants using an 80 MHz laser repetition rate.

**Supplementary tables**

Supplementary Table 1. Characteristic properties of fluorescent amino acids.

Supplementary Table 2. Fluorescence lifetimes of CBDS in different solvents.

Supplementary Table 3. Mutations found in six G1PylRS variants selected for activity with NBDO and NBDSe.

Supplementary Table 4. DNA and corresponding amino acid sequences of the proteins used in the current work.

Supplementary Table 5. Fluorescence emission maxima of CBDS in different solvents and protein environments used for TCSPC measurements.

Supplementary Table 6. Crystallographic statistics of H11-W37NBDO and H11-W37NBDSe.

**References**

### General Synthesis

All chemicals and solvents were of reagent grade (>95%) and used as received unless otherwise noted. Chemicals were obtained from Ambeed or from Merck. Unless otherwise specified, reactions were carried out in oven-dried glassware, under an atmosphere of nitrogen. Nuclear magnetic resonance (NMR) spectra were recorded at 300 K using either a 600 Bruker Avance (equipped with a high resolution cryogenic triple nucleus probehead), 500 Bruker Avance DPX 400 or a Bruker Avance 300 spectrometer. NMR spectra were calibrated to the residual solvent peak in $CDCl_3$ ($\delta$ = 7.26/77.16 ppm), MeOD-*d4* ($\delta$ = 3.3/49.0 ppm) or DMSO-*d6* ($\delta$ = 2.50/39.52 ppm) at 300 K. $^1$H NMR and $^{13}$C{$^1$H} NMR spectra were recorded at the indicated frequencies. Chemical shifts are expressed as parts per million (ppm) and are referenced to solvent residual signals. The data are reported as chemical shift, multiplicity (s = singlet, d = doublet, t = triplet, m = multiplet), coupling constant *J* in Hz and relative integral. High resolution mass spectra were recorded on a Bruker Apex II Fourier Transform Ion Cyclotron Resonance (FTICR) mass spectrometer with a 7.0 T magnet, fitted with an off-axis Analytic electrospray source with quadrupole mass analyser, and are reported as *m/z* (relative intensity).

### Synthesis

(*S*)-2-((*tert*-Butoxycarbonyl)amino)-3-((7-nitrobenzo[*c*][1,2,5]oxadiazol-4-yl)amino)propanoic acid (**Boc-NBDO**)

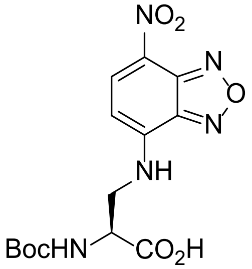

NBD-Cl (2.467 g, 12.4 mmol) was added to a mixture of Boc-Dap-OH (2.691 g, 13.2 mmol) and $K_2CO_3$ (3.41 g, 24.7 mmol) in MeOH (41 mL). The resulting mixture was heated to 50 °C for 70 minutes. Once reaction was complete, the deep red mixture was cooled to room temperature and the solvent removed *in vacuo*. Ethyl acetate (300 mL) and 1 M aqueous HCl solution (200 mL) were added to dissolve the residue. The organic and aqueous phases were partitioned, and the aqueous phase extracted three times with EtOAc (1000 mL total). The combined organic phases, were washed with brine (1x 200 mL), dried ($Na_2SO_4$), and the solvent removed *in vacuo* to afford the product as a deep red solid (4.46 g, 98%). Mp: 160 °C (decomposition); $^1$H NMR (300 MHz, MeOD): $\delta$ 8.54 (d, *J* = 8.8 Hz, 1H), 6.49 (d, *J* = 8.8 Hz, 1H),

4.55 (dd, *J* = 7.9, 5.0 Hz, 1H), 4.08 – 3.79 (m, 2H), 1.37 (s, 9H); $^{13}C\{^1H\}$ NMR (126 MHz, MeOD): δ 173.1, 157.9, 146.6 (broad), 145.9, 145.4, 138.2, 100.3 (broad), 80.9, 53.9, 46.1, 20.6; LRMS calc. for $C_{14}H_{18}N_5O_7^+$ [M + H]$^+$: 368, found 368; IR (ATIR): 3238, 1761, 1674, 1588, 1294, 1129 cm$^{-1}$.

(*S*)-2-Amino-3-((7-nitrobenzo[*c*][1,2,5]oxadiazol-4-yl)amino)propanoic acid hydrochloride (**NBDO·HCl**)

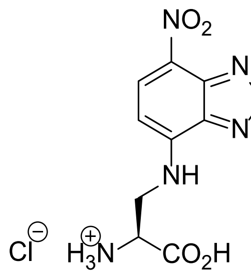

4 M HCl in dioxane (30 mL) was added dropwise to a solution of **Boc-NBDO** (4.46 g, 12.1 mmol) in anhydrous THF (60 mL) at room temperature. The resulting solution was heated to reflux for 75 minutes. The solvent was removed *in vacuo* and the resulting solid triturated repeatedly with $Et_2O$ and dichloromethane until clean. The product was collected as a deep red solid (3.579 g, 97%), with all analytical data matching that previously reported in the literature.[1] Mp: 160 °C (decomposition); $^1H$ NMR (300 MHz, MeOD): δ 8.58 (d, *J* = 8.6 Hz, 1H), 6.53 (d, *J* = 8.6 Hz, 1H), 4.44 – 4.39 (m, 1H), 4.19 (dd, *J* = 15.0 Hz, 4.8 Hz, 1H), 4.09 (dd, *J* = 15.0 Hz, 7.7 Hz, 1H); $^{13}C\{^1H\}$ NMR (126 MHz, MeOD): δ 169.6, 145.9, 145.8, 145.2, 137.9, 125.1, 101.2, 53. 0, 44.2; LRMS calc. for $C_9H_{10}N_5O_5^+$ [M + H]$^+$: 268, found 268; IR (ATIR, solid): 3389, 3066 (broad), 1622, 1488, 1259 cm$^{-1}$.

(*S*)-2-((*tert*-Butoxycarbonyl)amino)-3-((7-nitrobenzo[*c*][1,2,5]selenadiazol-4-yl)amino)propanoic acid (**Boc-NBDSe**)

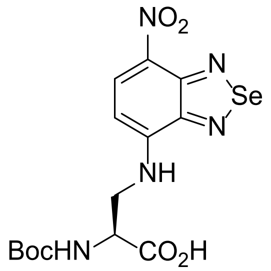

Triethylamine (570 μL, 4.09 mmol) was added to a solution of NBDSe-Cl[2] (357 mg, 1.36 mmol) and Boc-Dap-OH (337 mg, 1.65 mmol) in DMSO (4.5 mL) at room temperature. The resulting solution was heated to 60 °C for 2h. Once complete, the reaction mixture was poured into 50 mL of 0.1 M aqueous HCl solution. The resulting precipitate was collected by vacuum filtration. The precipitate was washed with ethyl acetate to afford the desired product as a deep red solid (297 mg, 51 %). The ethyl acetate wash and aqueous phase were separated, and the aqueous phase extracted once more with ethyl acetate. The combined ethyl acetate layers were washed with brine, and the solvent removed *in vacuo*. Recrystallization of the resulting solid from hot ethyl acetate yielded a further batch of the desired compound (213 mg, 36%, 87% total). m.p.: >200 °C; $^1$H NMR (500 MHz, DMSO-*d6*): δ 8.58 (d, *J* = 8.9 Hz, 1H), 8.19 (t, *J* = 6.0 Hz, 1H), 7.32 (d, *J* = 7.9 Hz, 1H), 6.45 (d, *J* = 8.9 Hz, 1H), 4.37 – 4.30 (m, 1H), 3.86 – 3.70 (m, 1H), 1.30 (s, 9H); $^{13}$C{$^1$H} NMR (126 MHz, DMSO-*d6*): δ 171.9, 155.4, 151.9, 148.1, 135.0, 128.5, 97.6, 78.4, 52.7, 43.6, 28.0; LRMS (ESI): $[M+H]^+$ calc. for $C_{14}H_{18}N_5O_6Se$, 432; found 432. IR (ATIR, solid): 2973, 1687, 1550, 1244, 1156, 1125 $cm^{-1}$.

(*S*)-2-Amino-3-((7-nitrobenzo[*c*][1,2,5]selenadiazol-4-yl)amino)propanoic acid hydrochloride (**NBDSe ·HCl**)

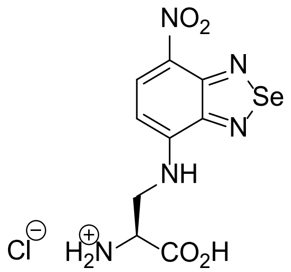

HCl in dioxane (3.5 mL, 4M solution) was added to a solution of **Boc-NBDSe** (297 mg, 0.690 mmol) in anhydrous THF (10 mL) at room temperature. The resulting solution was heated to 60 °C for 4 h. The solvent was removed under a flow of nitrogen, and the resulting solid triturated with dichloromethane to afford the desired compound as a deep red solid (251 mg, *quant.*) with all analytical data matching that previously reported in the literature.[1] m.p.: > 200 °C; $^1$H NMR (300 MHz, DMSO-*d6*): δ 8.61 (d, *J* = 8.9 Hz, 1H), 8.54 (broad s, 3H), 8.25 (t, *J* = 6.2 Hz, 1H), 6.56 (d, *J* = 8.9 Hz, 1H), 4.39 – 4.21 (m, 1H), 4.10 – 3.87 (m, 2H); $^{13}$C{$^1$H} NMR (126 MHz, DMSO-*d6*): δ 169.1, 152.0, 151.7, 147.7, 134.7, 129.2, 98.0, 51.2, 42.7; LRMS (ESI): $[M+H]^+$ calc. for $C_9H_{10}N_5O_4Se$, 332; found 332; IR (ATIR, solid): 3272, 1715, 1555, 1268 $cm^{-1}$.

(*S*)-2-((*tert*-Butoxycarbonyl)amino)-3-((7-cyanobenzo[c][1,2,5]thiadiazol-4-yl)amino)propanoic acid (**Boc-CBDS**)

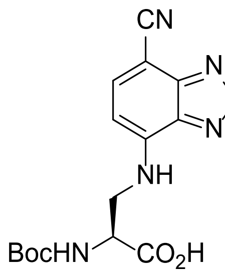

A microwave vial was charged with 4,7-dibromobenzo[*c*][1,2,5]thiadiazole (991 mg, 3.37 mmol), CuCN (301 mg, 3.36 mmol) and N,N-dimethylacetamide (11 mL). The resulting mixture was sparged with argon for 10 min, then heated under microwave conditions to 200 °C for 10 min. The reaction mixture was cooled, diluted with saturated aqueous $NaHCO_3$ solution, and extracted with ethyl acetate (3x 200 mL). The organic extracts were combined, washed with brine, dried ($Na_2SO_4$), and the solvent removed *in vacuo*. The crude mixture of starting material, mono-cyanated, and di-cyanated products, was then dissolved in DMSO (11 mL). Boc-Dap-OH (274 mg, 1.34 mmol) and DIPEA (580 µL, 3.32 mmol) were added, and the resulting solution was heated to 100 °C for 17 h. The reaction mixture was cooled to room temperature, diluted with ethyl acetate, and washed with 1H aqueous HCl solution (5x 50 mL) and brine (50 mL), then dried ($Na_2SO_4$) and the solvent removed *in vacuo*. Flash column chromatography, eluting with a gradient from 100% $CH_2Cl_2$ → 15% MeOH/84% $CH_2Cl_2$/1% AcOH, afforded the desired compound as an orange-brown solid (145 mg, 12%). Mp: 180 °C (decomp.); $^1$H NMR (300 MHz, MeOD): δ 7.90 (d, *J* = 7.9 Hz, 1H), 6.60 (d, *J* = 7.9 Hz, 1H), 4.46 – 4.37 (m, 1H), 3.88 (dd, *J* = 13.8, 4.6 Hz, 1H), 3.76 – 3.67 (m, 1H), 1.38 (s, 9H); $^{13}$C{$^1$H} NMR (MeOD, 126 MHz): δ 157.9, 155.4, 147.6, 146.4, 141.1, 118.4, 102.0, 89.5, 80.7, 55.6 (broad), 46.3, 28.7; HRMS (ESI): $[M+H]^+$ calc. for $C_{15}H_{18}N_5O_4S$, 364.10740; found 364.10765; IR (ATIR, solid): 3341, 2928, 2211, 1698, 1562, 1158 $cm^{-1}$.

(*S*)-2-Amino-3-((7-cyanobenzo[*c*][1,2,5]thiadiazol-4-yl)amino)propanoic acid hydrochloride (**CBDS·HCl**)

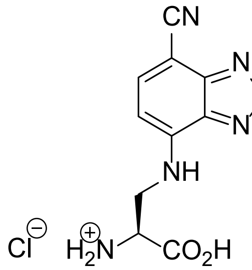

HCl in dioxane (1.7 mL, 4M solution) was added to a solution of **Boc-CBDS** (135 mg, 0.371 mmol) in anhydrous THF (4 mL) at room temperature. The resulting solution was heated at reflux for 2.5 h. The solvent was removed under a flow of nitrogen, and the resulting solid triturated with dichloromethane to afford the desired compound as an orange-brown solid (110 mg, *quant.*). m.p. 170°C (decomp.); $^{1}$H NMR (500 MHz, DMSO-*d6*): δ 8.09 (d, *J* = 8.2 Hz, 1H), 7.95 (t, *J* = 5.8 Hz, 1H), 6.69 (d, *J* = 8.2 Hz, 1H), 4.10 – 4.05 (m, 1H), 3.97 – 3.83; $^{13}$C{$^{1}$H} NMR (126 MHz, DMSO-*d6*): δ 169.0, 153.8, 145.9, 144.3, 139.9, 117.5, 101.5, 88.1, 51.5, 42.8; HRMS (ESI): $[M+Na]^+$ calc. for $C_{10}H_9N_5NaO_2S$, 286.03692; found 286.03658; IR (ATIR, solid): 2846 (broad), 2210, 1728, 1557, 879, 514 cm$^{-1}$.

**Boc-NBDO**: $^{1}$H NMR (500 MHz, MeOD)

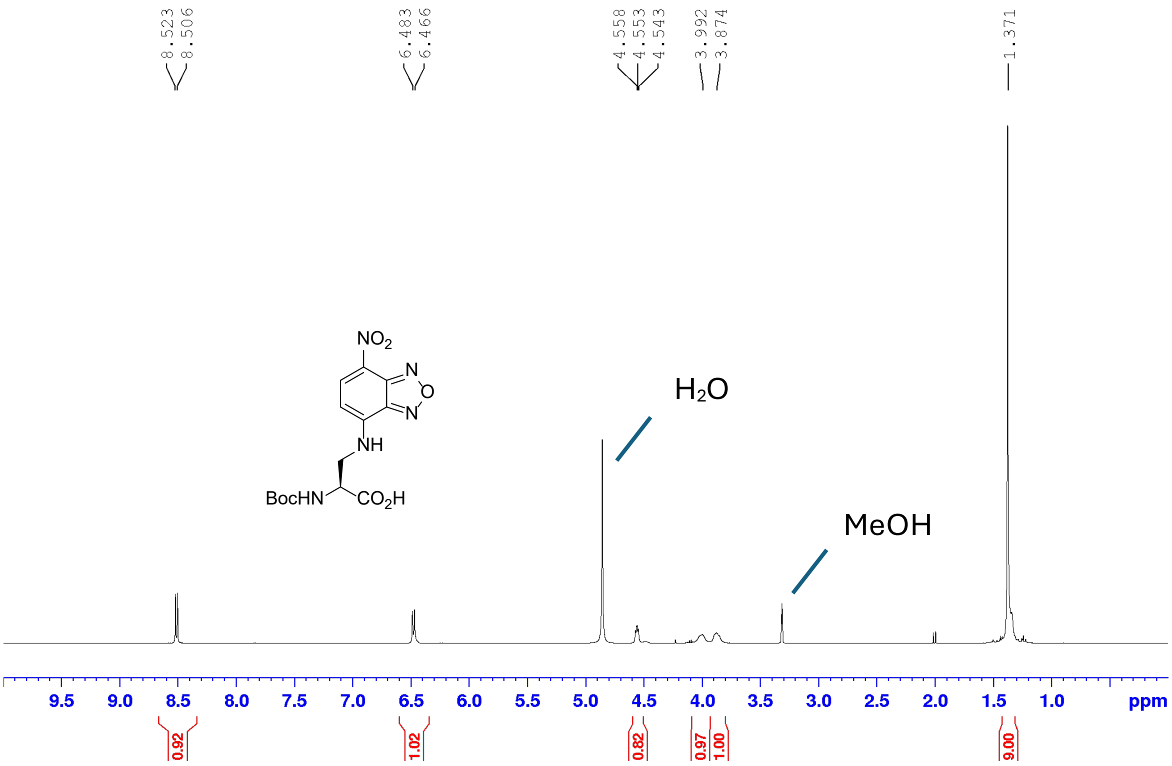

**Boc-NBDO**: $^{13}$C{$^{1}$H} NMR (126 MHz, MeOD)

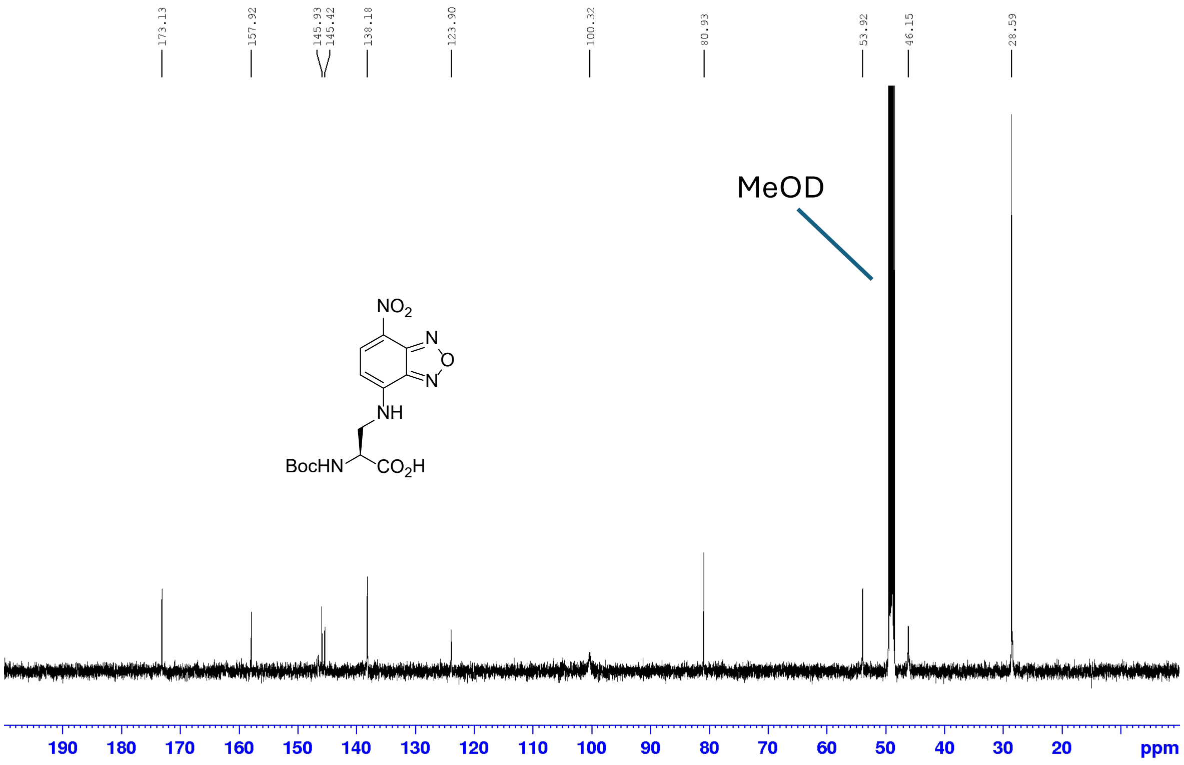

**NBDO·HCl**: $^1$H NMR (500 MHz, MeOD)

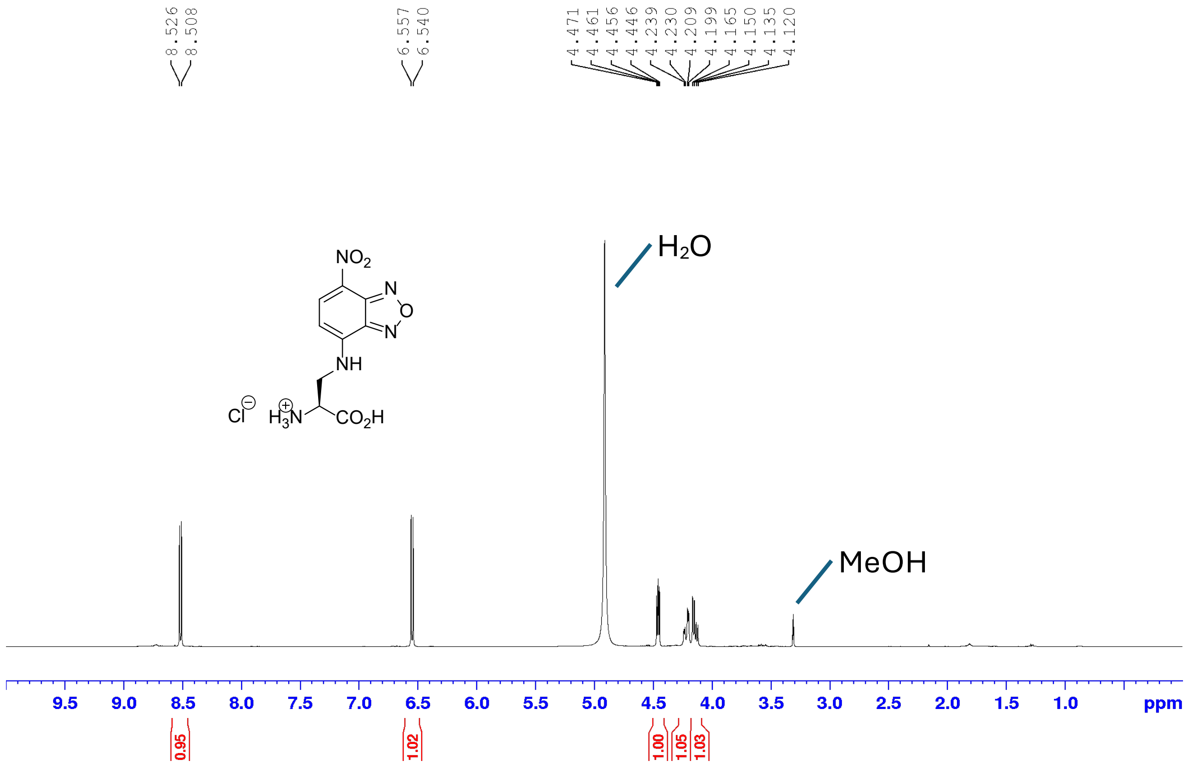

**NBDO·HCl**: $^{13}C\{^1H\}$ NMR (126 MHz, MeOD)

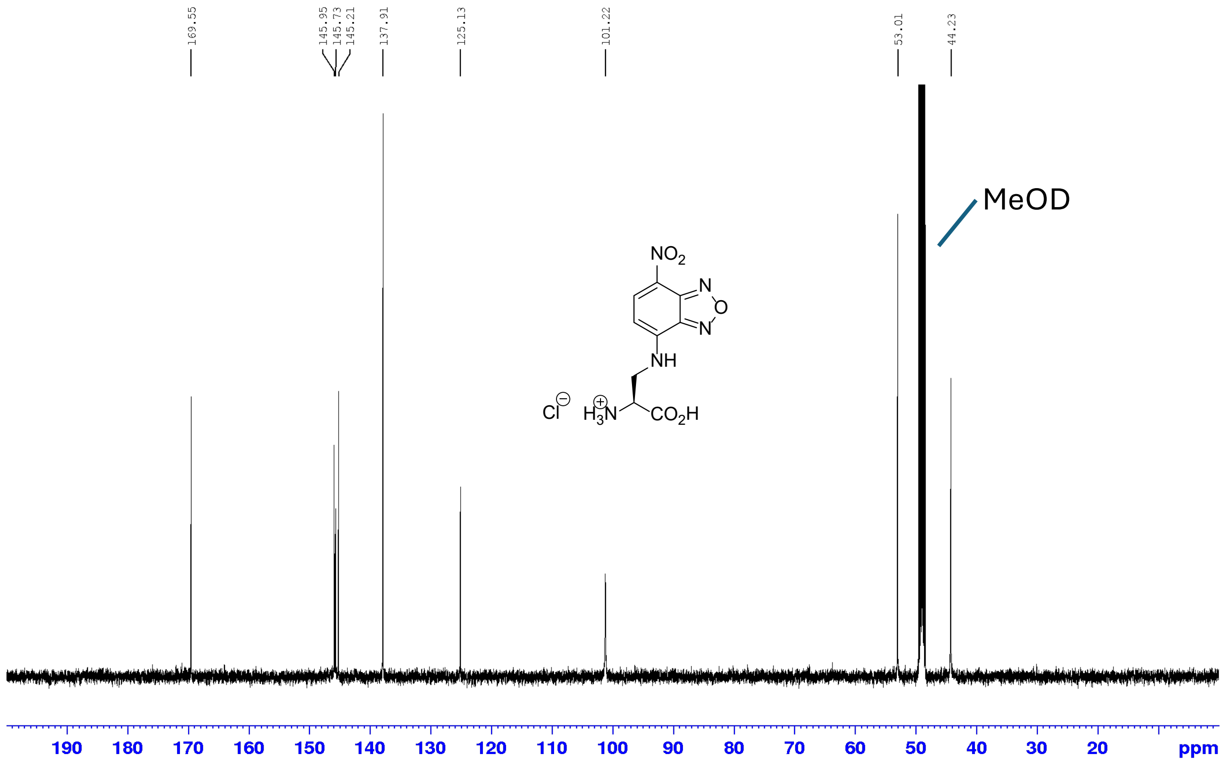

**Boc-NBDSe**: $^{1}H$ NMR (500 MHz, DMSO-*d6*)

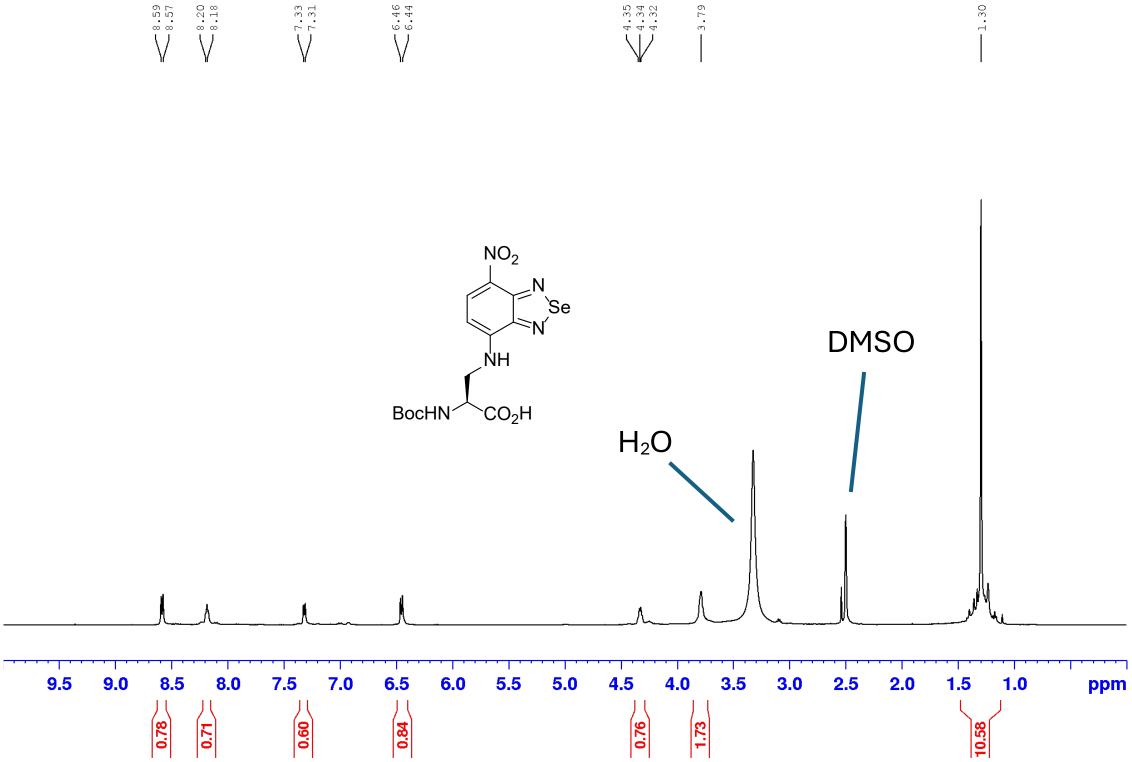

**Boc-NBDSe**: $^{13}C\{^{1}H\}$ NMR (126 MHz, DMSO-*d6*)

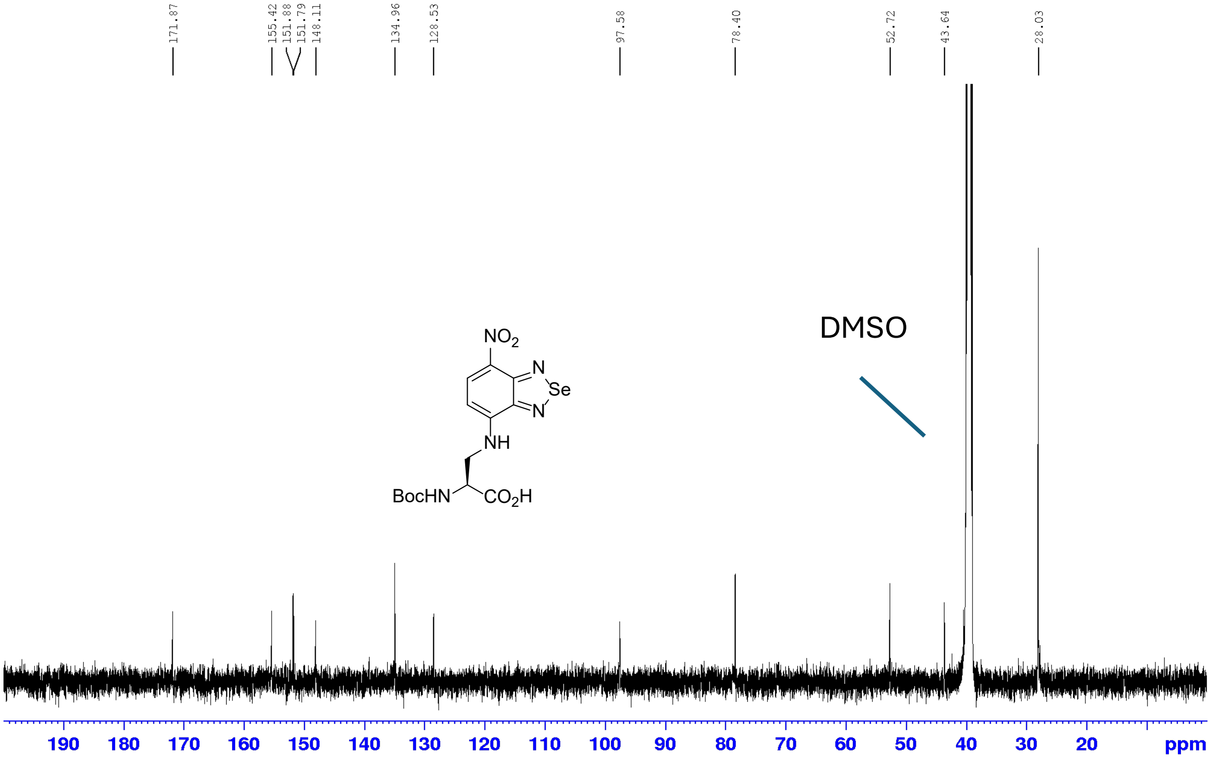

**NBDSe·HCl**: $^{1}$H NMR (500 MHz, DMSO-*d6*)

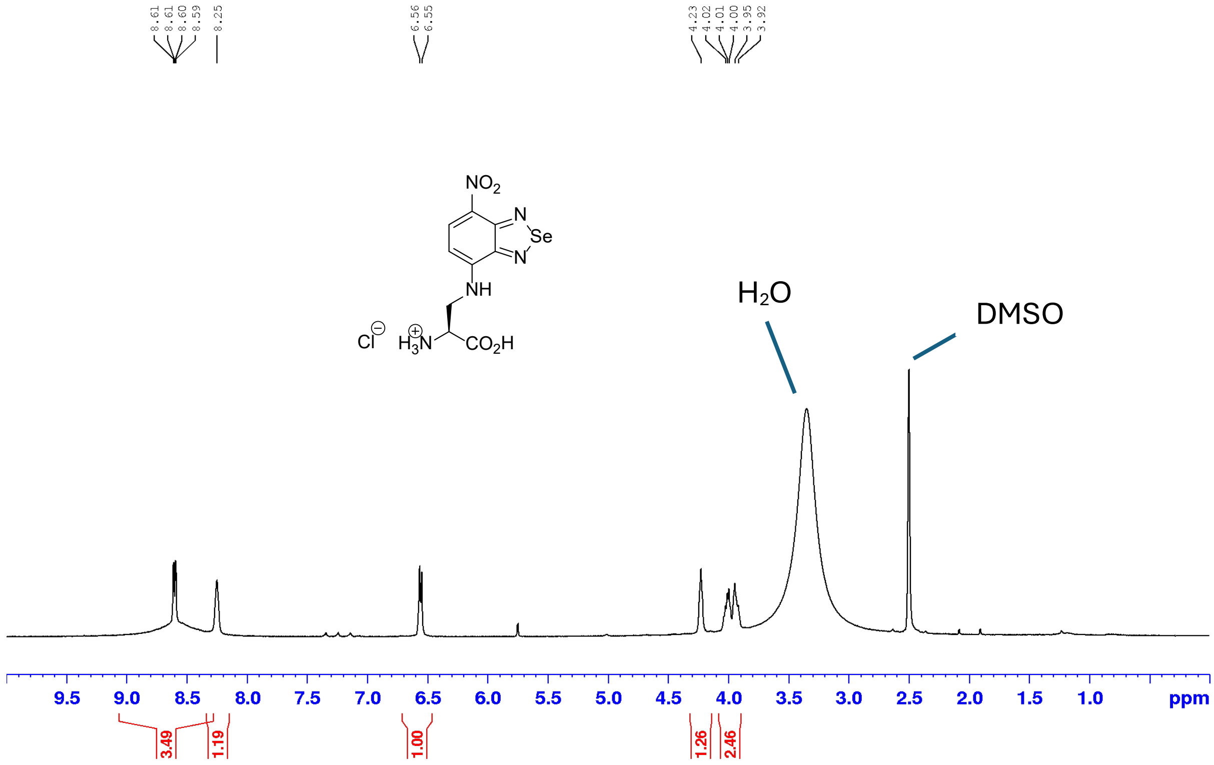

**NBDSe·HCl**: $^{13}$C{$^{1}$H} NMR (126 MHz, DMSO-*d6*)

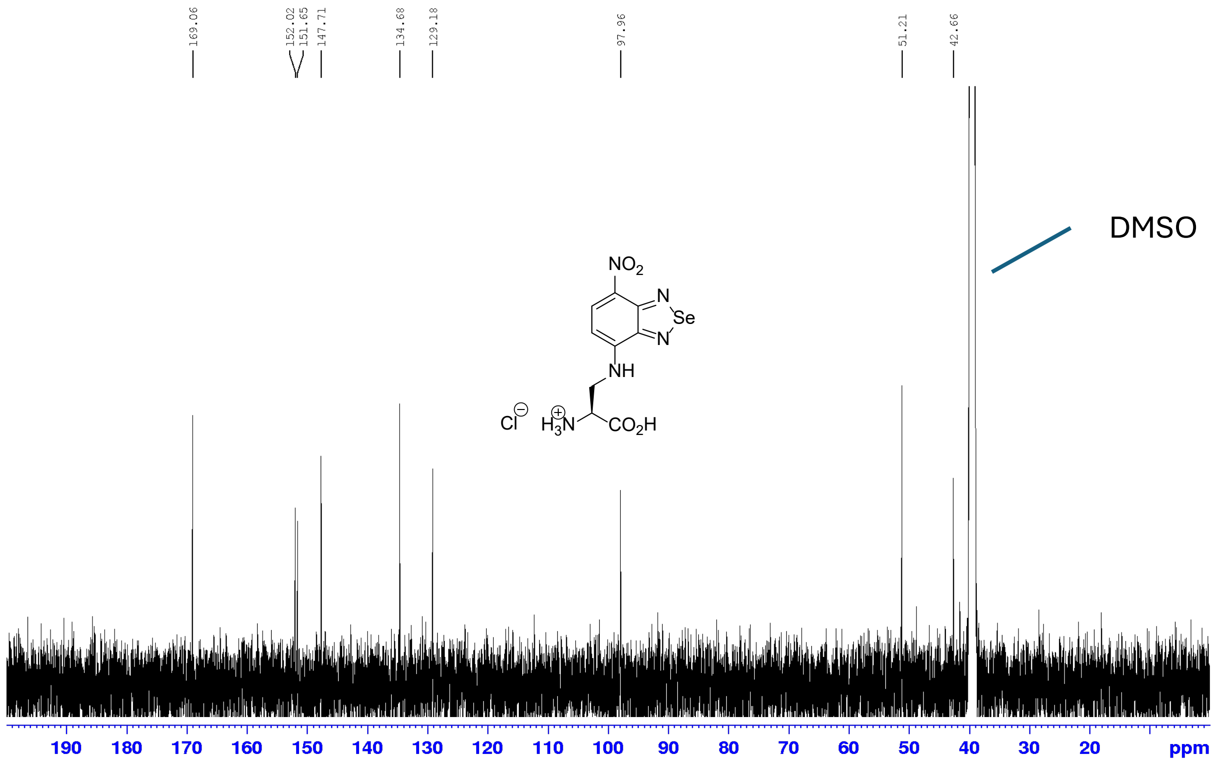

**Boc-CBDS**: $^{1}$H NMR (500 MHz, MeOD)

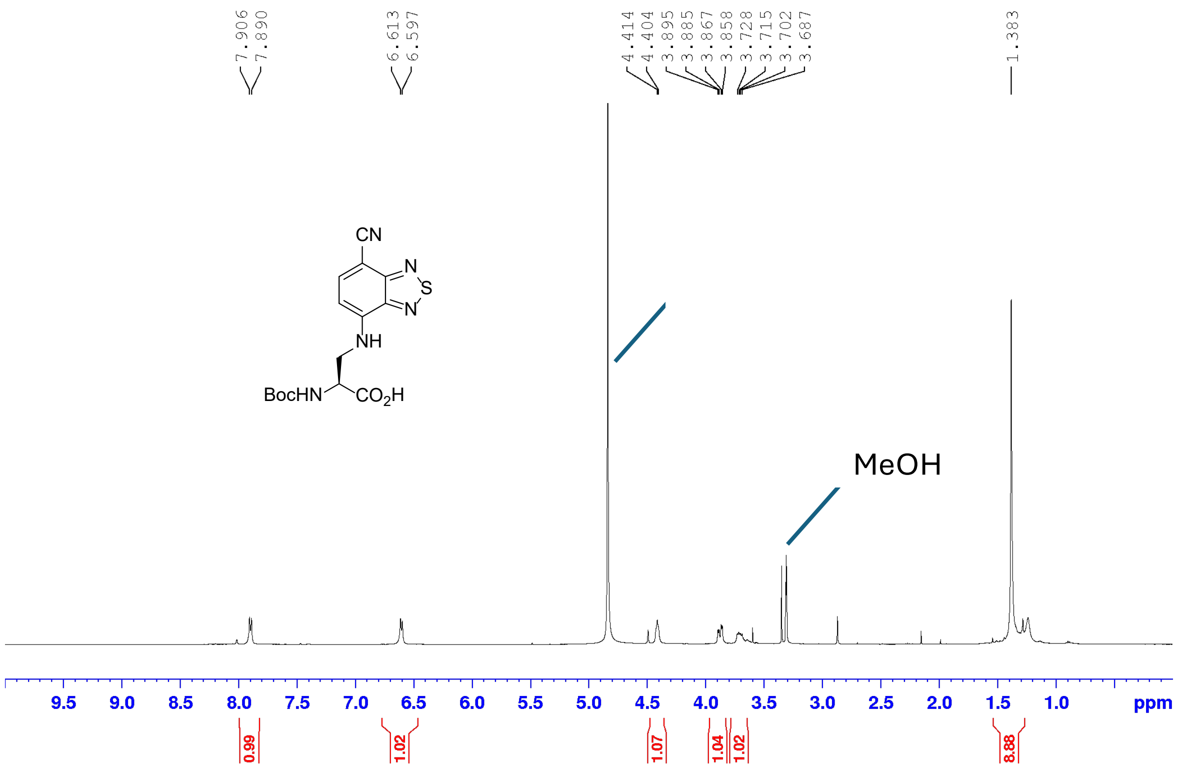

**Boc-CBDS**: $^{13}$C{$^{1}$H} NMR (500 MHz, MeOD)

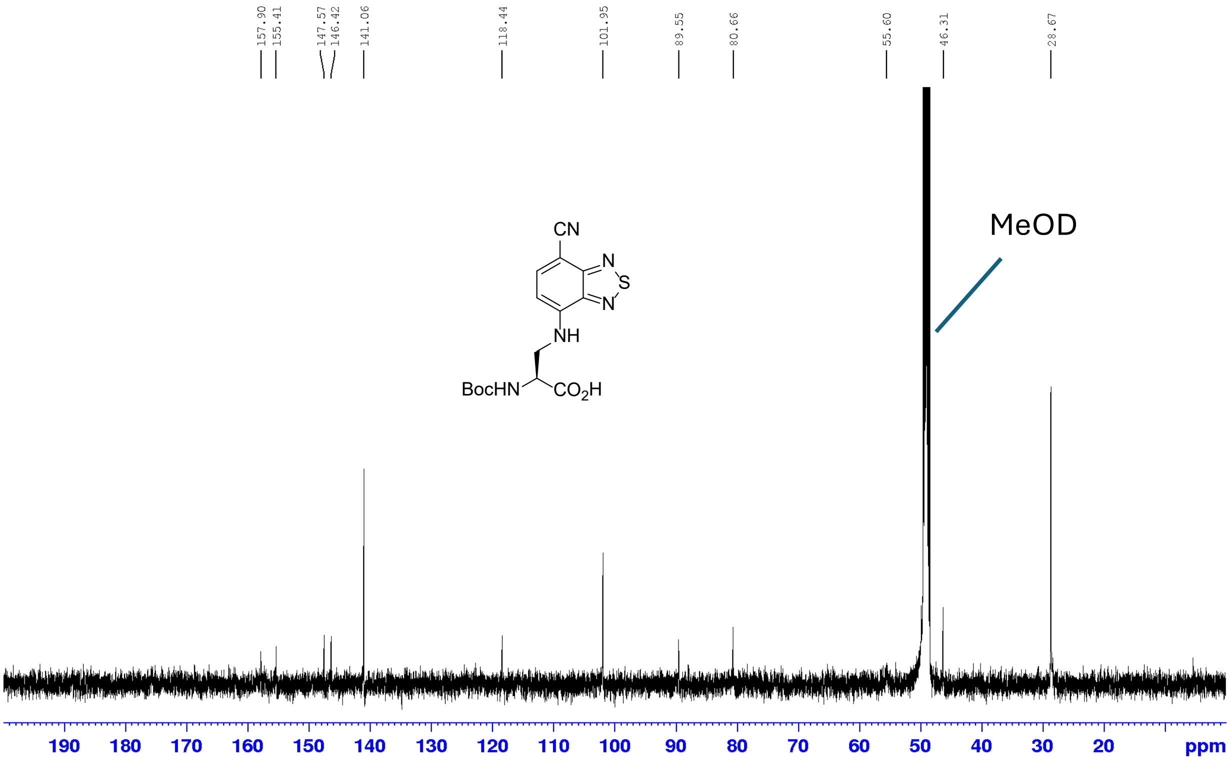

**CBDS·HCl**: $^{1}$H NMR (400 MHz, DMSO-*d6*) (water peak suppressed)

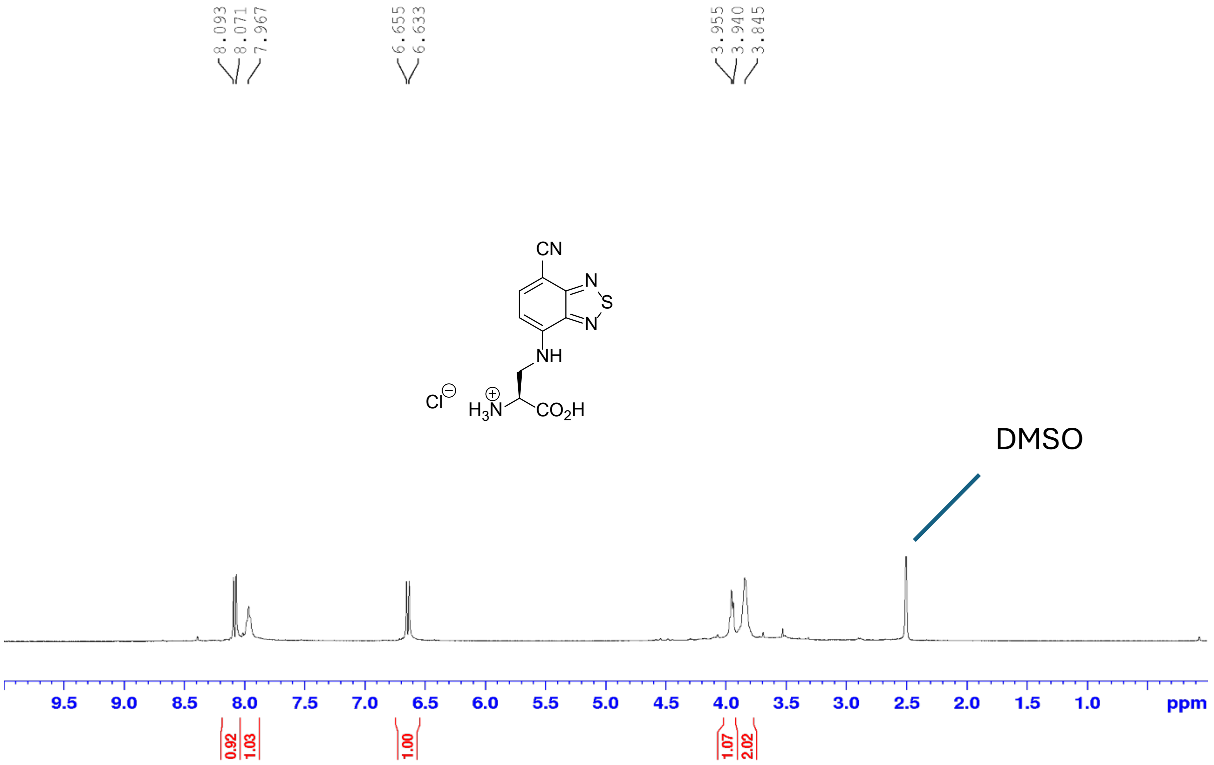

**CBDS·HCl**: $^{13}$C{$^{1}$H} NMR (DMSO-*d6*, 126 MHz)

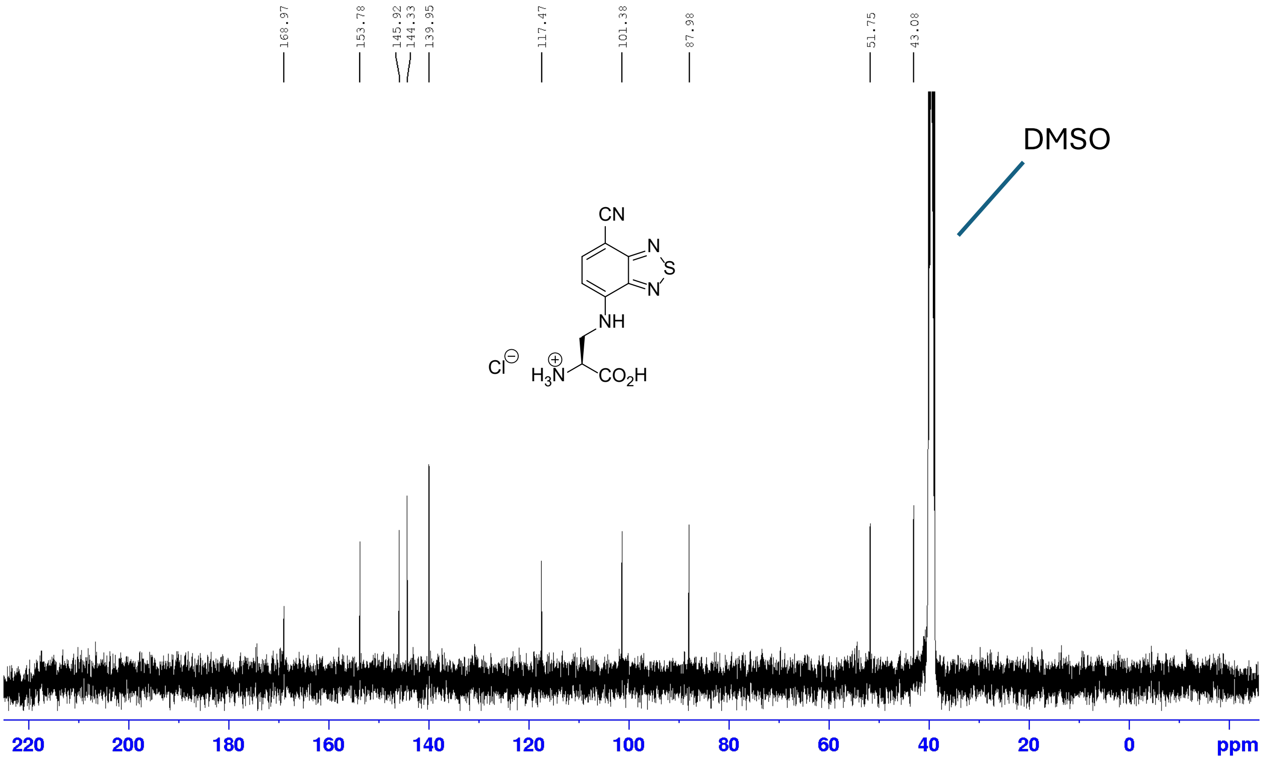

### IR spectra

#### Boc-NBDO

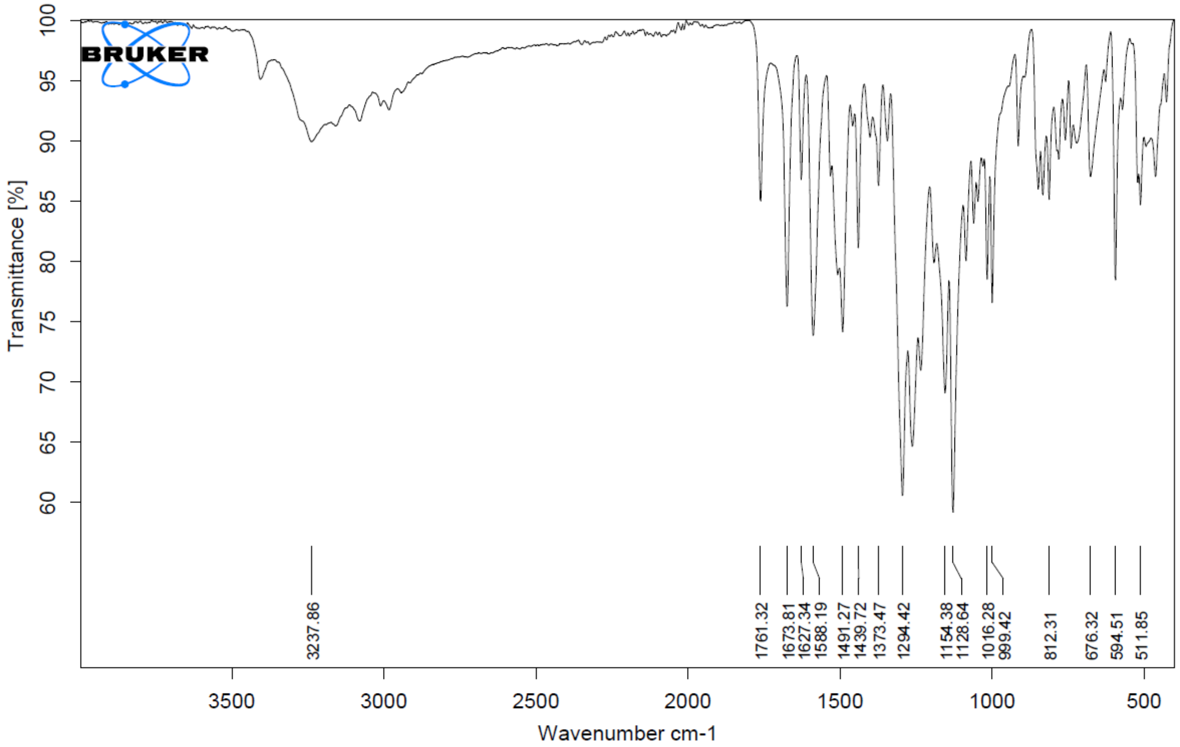

#### NBDO·HCl

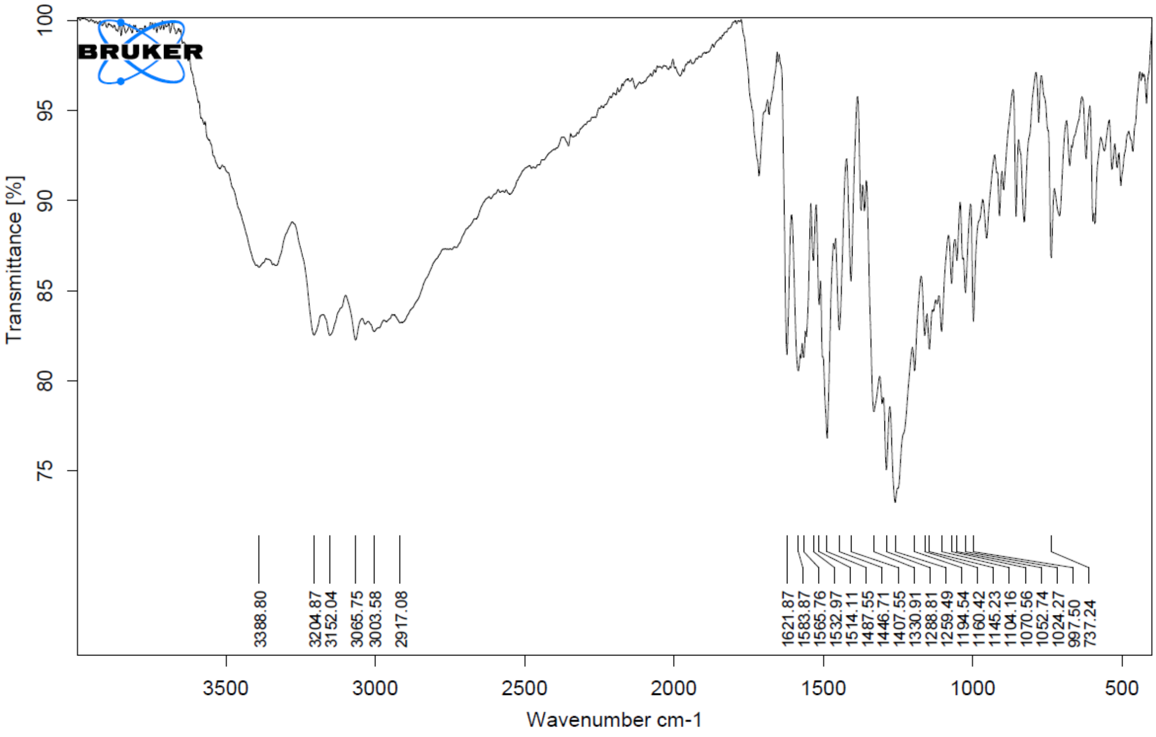

**Boc-NBDSe**

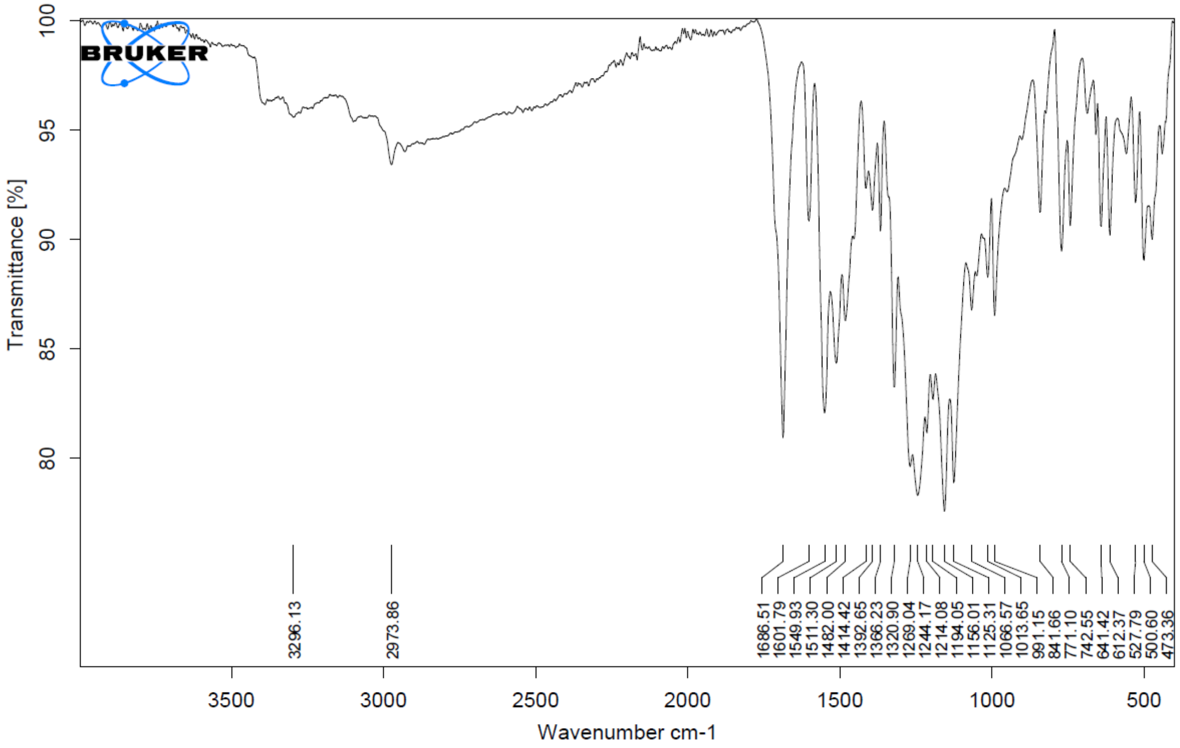

**NBDSe·HCl**

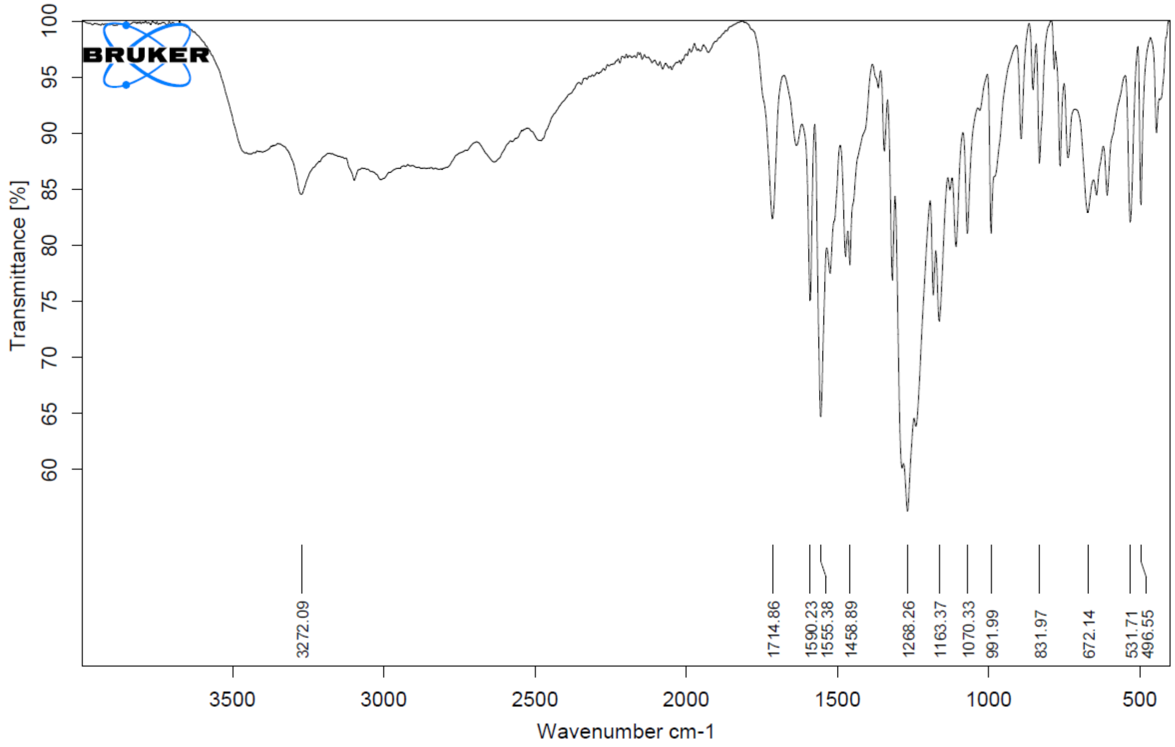

**Boc-CBDS**

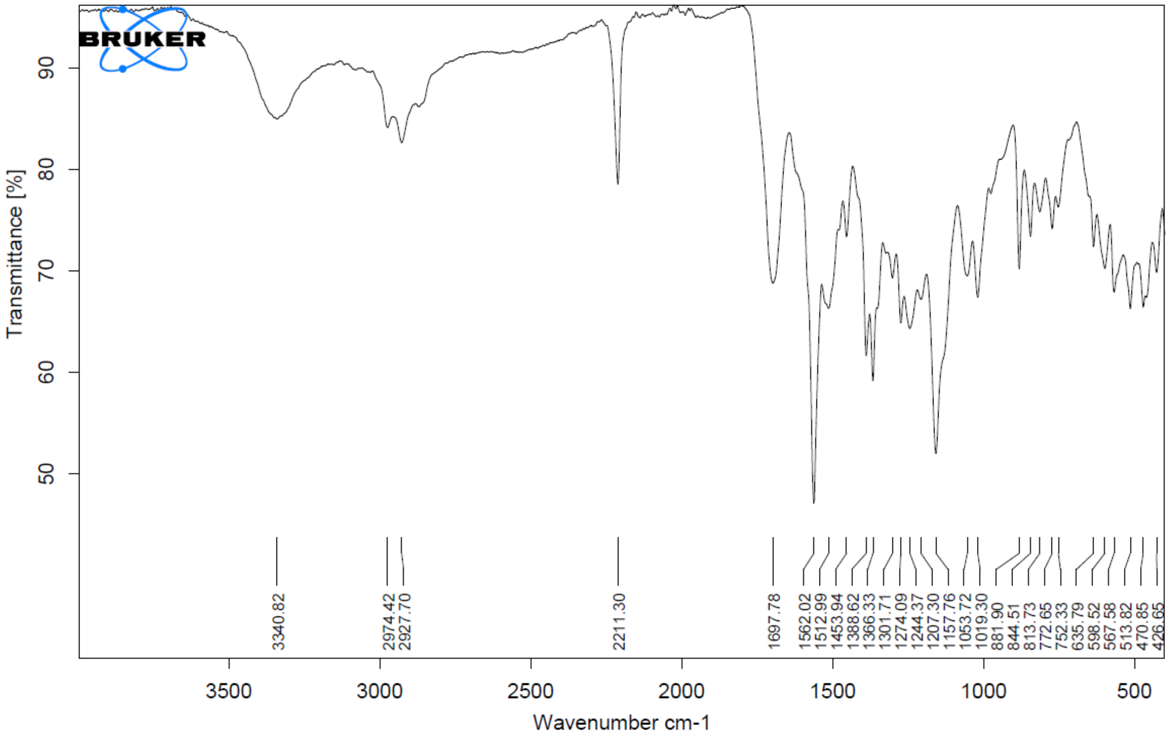

**CBDS·HCl**

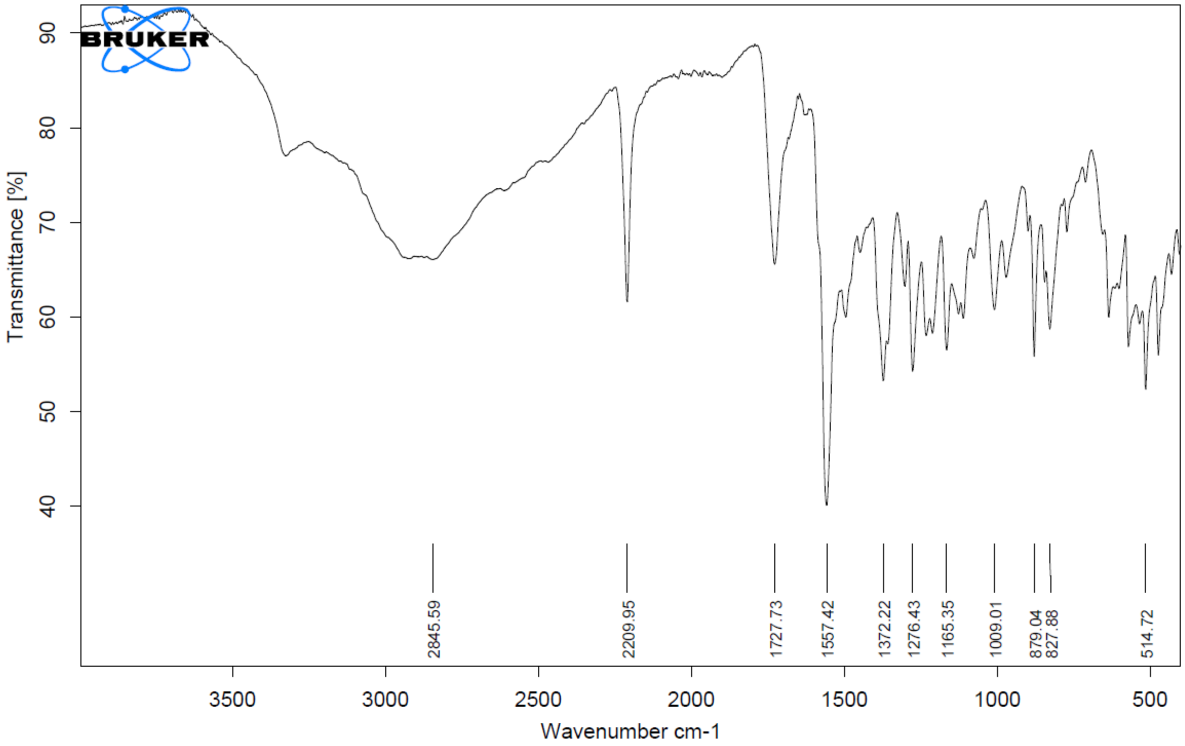

**HRMS**

**Boc-CBDS**

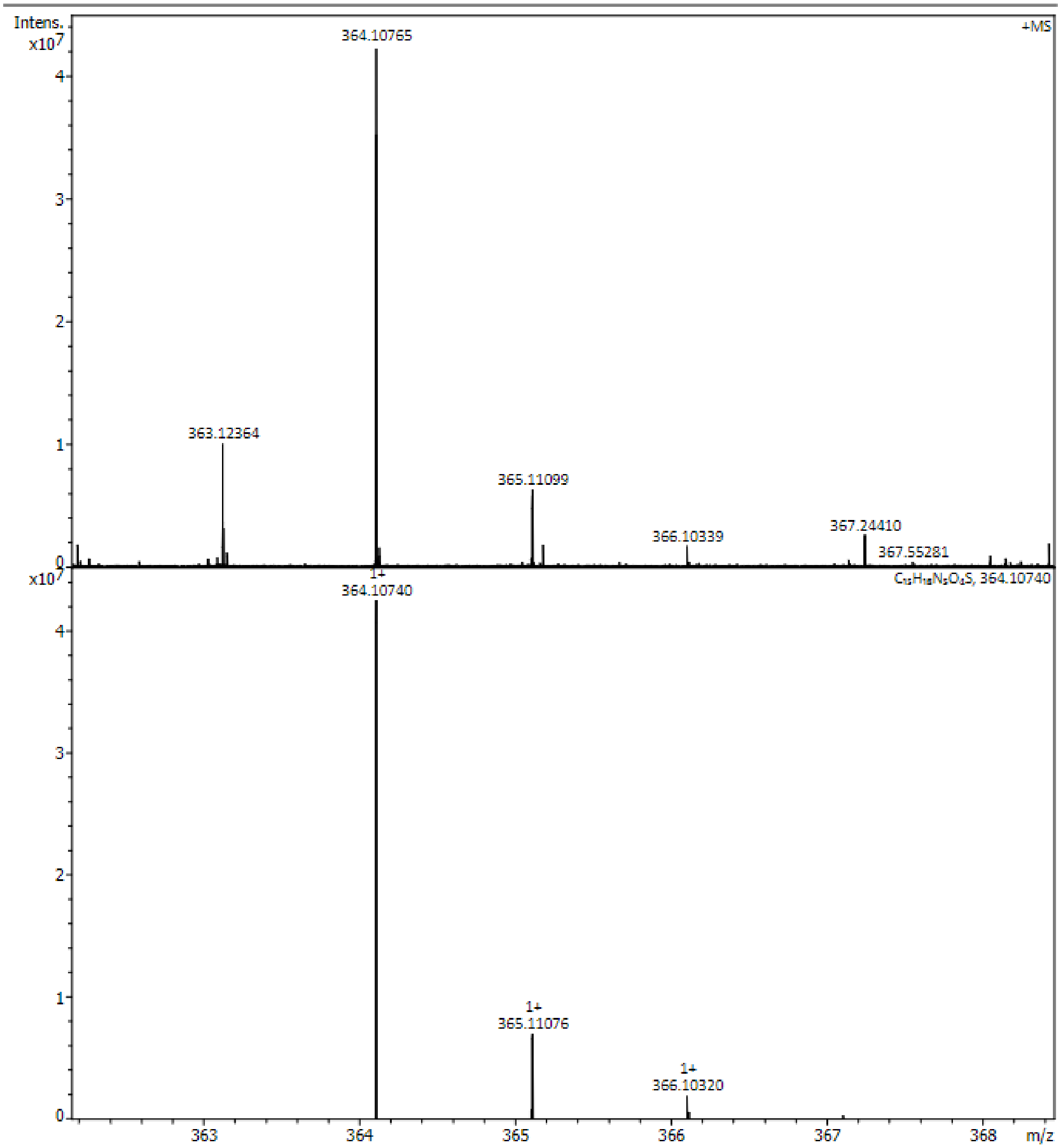

**Top:** nESI HRMS spectrum of Boc-CBDS; **Bottom:** Simulated mass spectrum.

**CBDS·HCl**

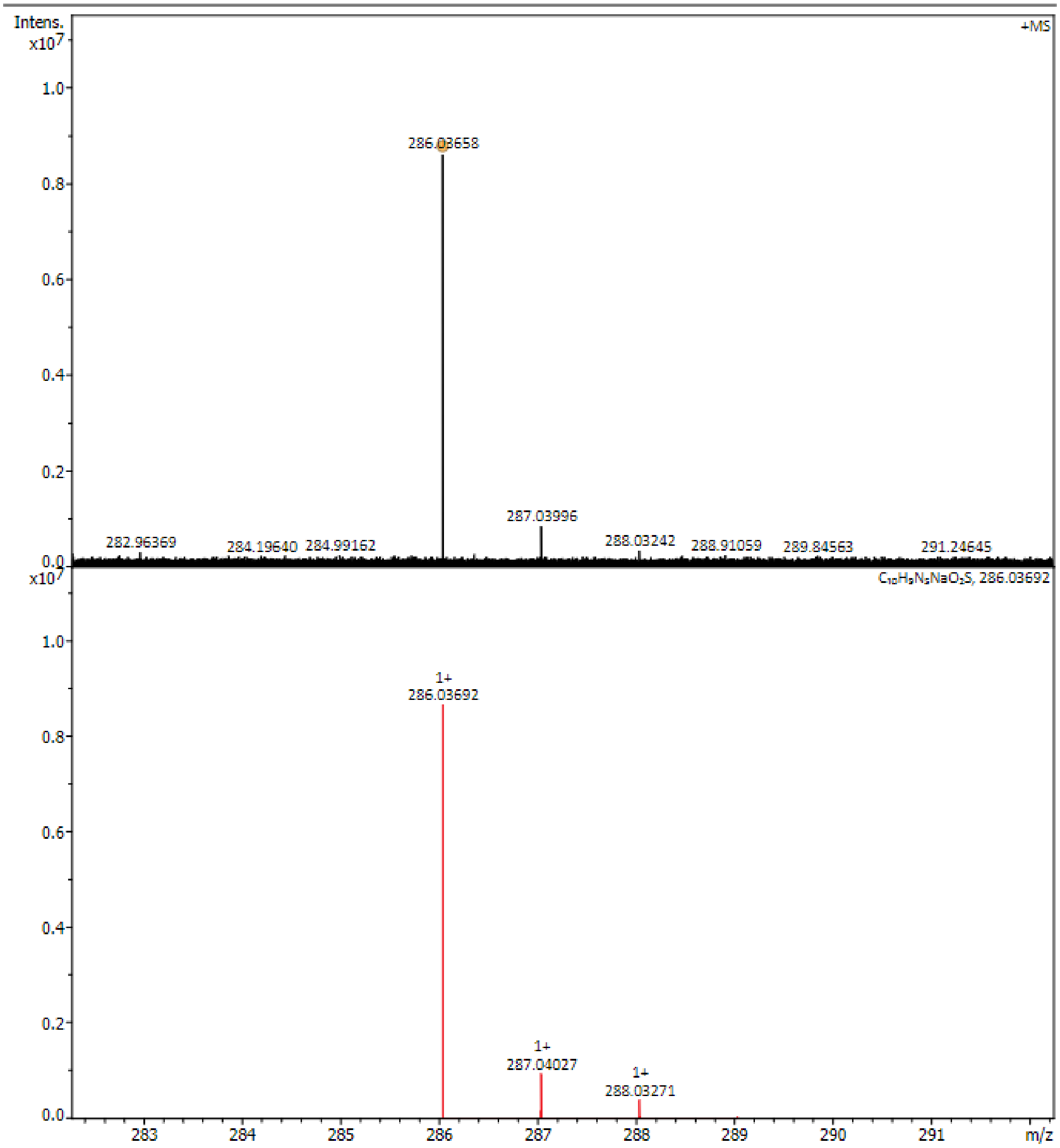

**Top:** nESI HRMS spectrum of CBDS·HCl; **Bottom:** Simulated mass spectrum.

### Supplementary figures

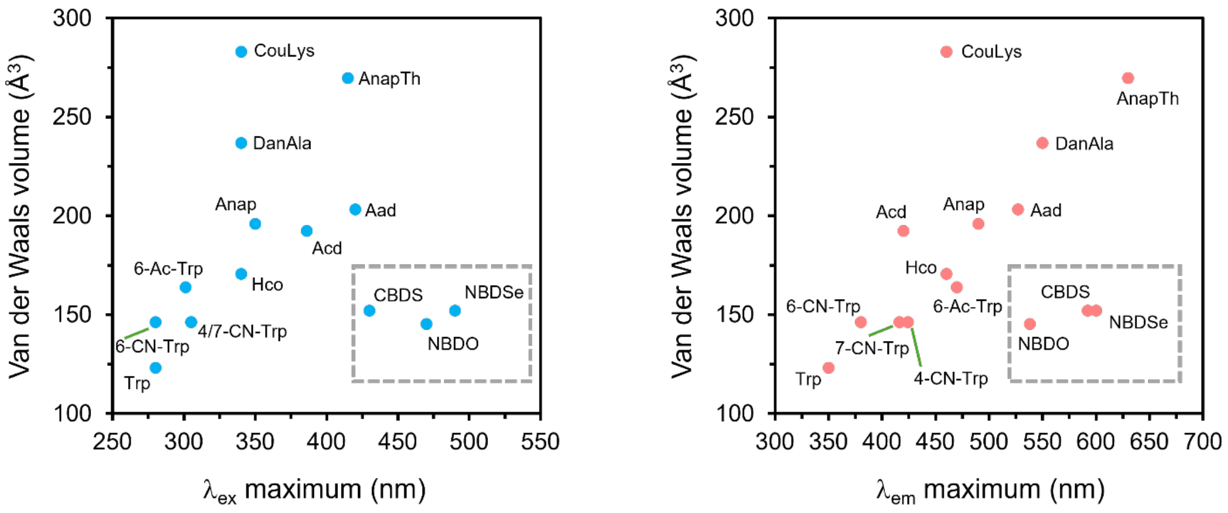

Supplementary Fig. 1. BD-ncAAs break the chromophore size–wavelength trade-off. Alternative representation of Figure 1b, with fluorescent amino acid comparators aligned by excitation or emission maximum wavelength and plotted against the corresponding van der Waals volume calculated from the side chain from the β-position onward. Most fluorescent amino acids show a general trend toward longer excitation or emission wavelengths with increasing chromophore size, whereas the three BD-ncAAs deviate from this trend.

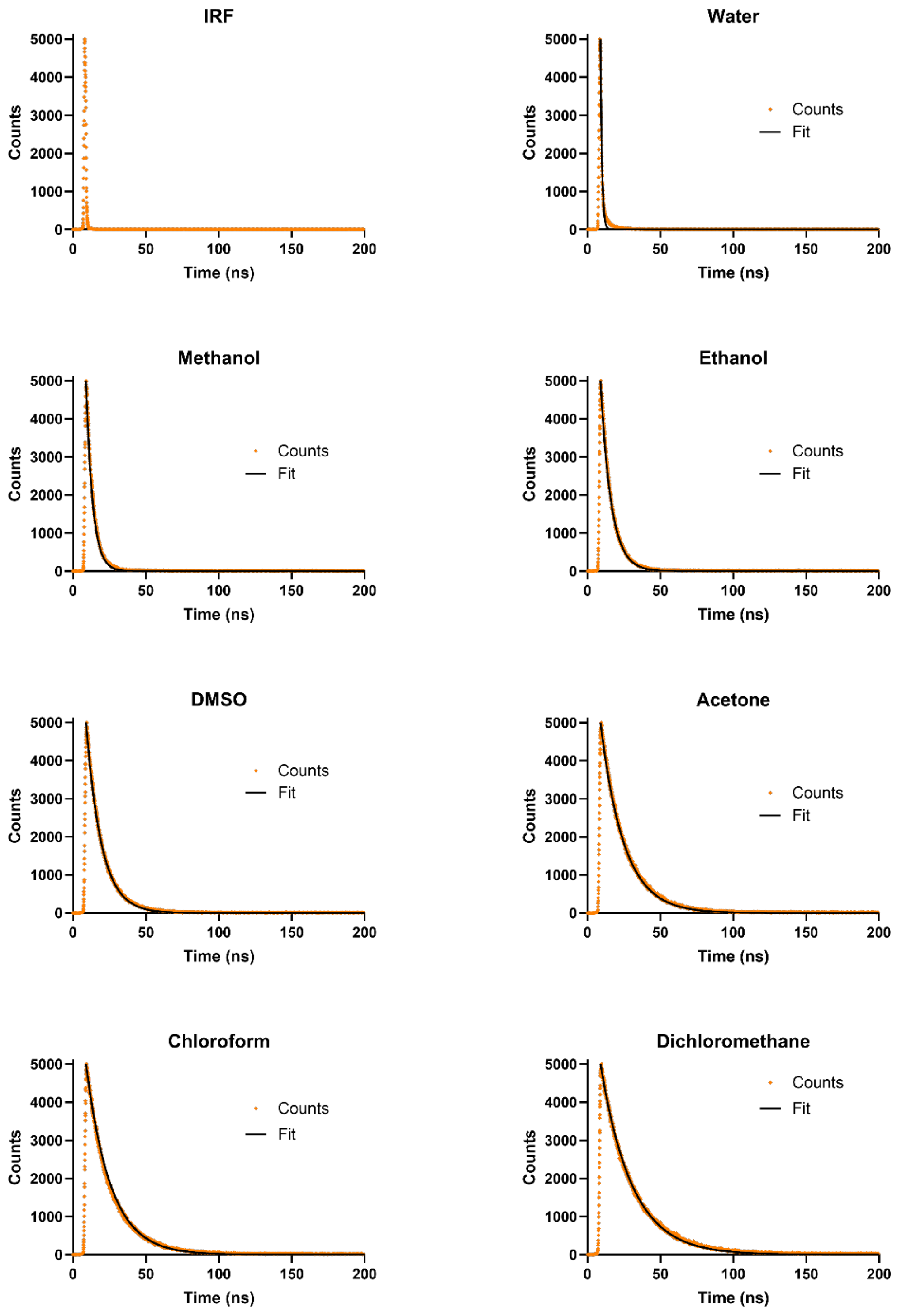

Supplementary Fig. 2. Fluorescence decay histogram of CBDS in different solvents (orange symbols) together with the reconvolution fit (black line). The decay was recorded using TCSPC and fitted using a single-exponential decay model convoluted with the instrument response function (IRF) measured under identical experimental conditions.

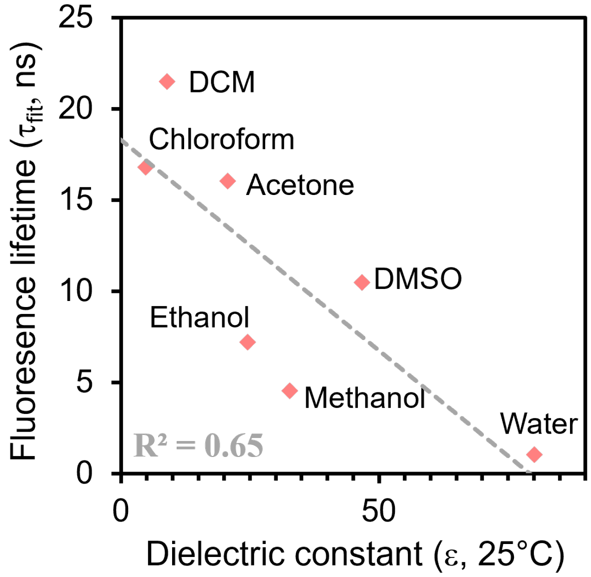

Supplementary Fig. 3. Plot of CBDS fluorescence lifetime versus solvent dielectric constant (at 25°C). The linear regression fit ($R^2$ = 0.65) demonstrates that there is no strong correlation between the fluorescence lifetime of CBDS and the dielectric constant of the solvent.

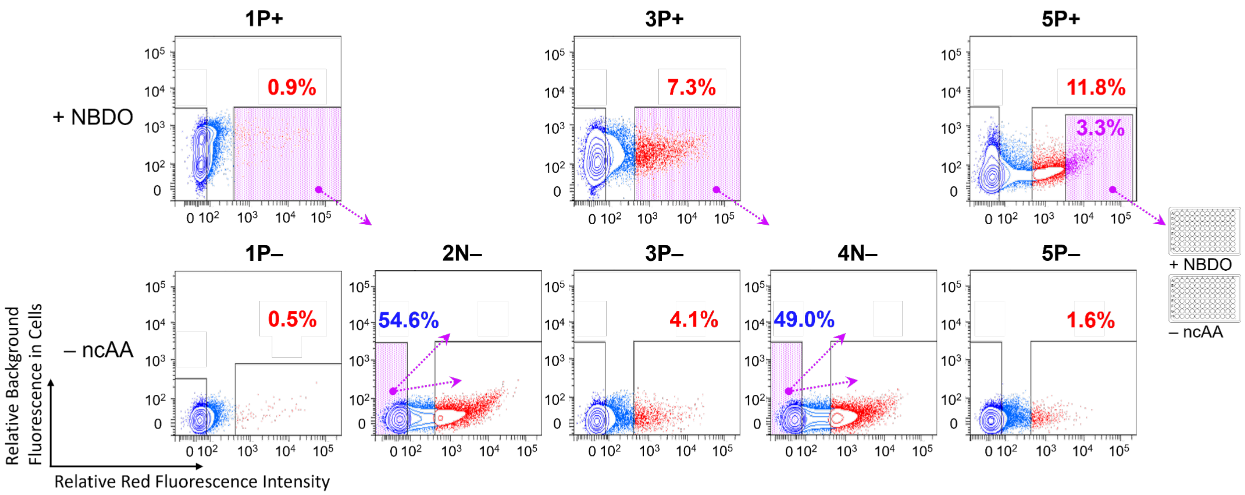

Supplementary Fig. 4. FACS experiments for selection of active and specific G1PylRS enzymes that recognize NBDO. The x-axis indicates red fluorescence intensity (excitation at 560 nm), and the y-axis reflects background fluorescence upon excitation at 488 nm. Positive samples (1P+, 3P+, 5P+) were prepared in the presence of 0.5 mM NBDO, while negative samples (1P–, 2N–, 3P–, 4N–, 5P–) were cultured without ncAA. Populations collected for subsequent rounds are shaded in purple, with percentages denoting the gated fluorescence-positive fraction. Arrows indicate the direction of the iterative enrichment strategy. The top 3.3% from the final round samples were collected for further characterization.

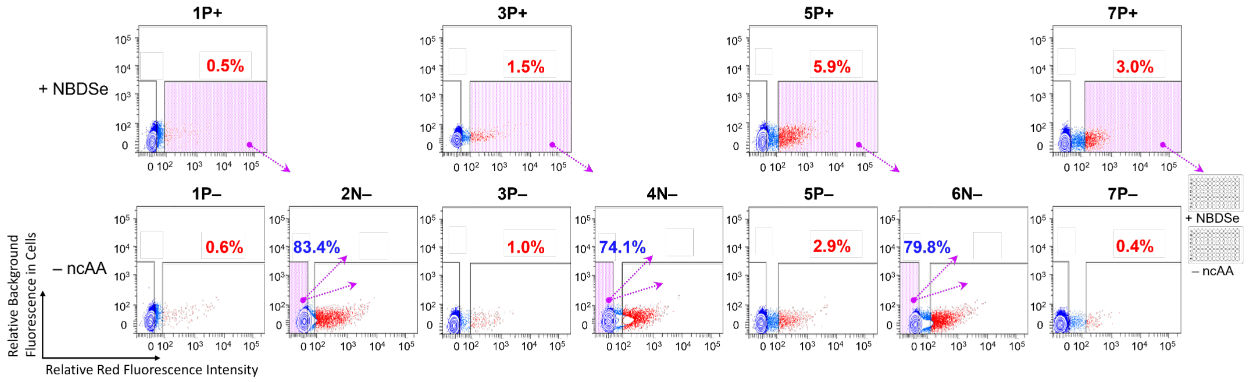

Supplementary Fig. 5. FACS experiments for selection of active and specific G1PylRS enzymes that recognize NBDSe. The axes, arrows, and shades have the same indication as in Supplementary Fig. 4. Positive samples (1P+, 3P+, 5P+, 7P+) were prepared in the presence of 0.5 mM NBDSe, while negative samples (1P–, 2N–, 3P–, 4N–, 5P–, 6N–, 7P–) were cultured without ncAA. The top 3% from the final round samples were collected for further characterization.

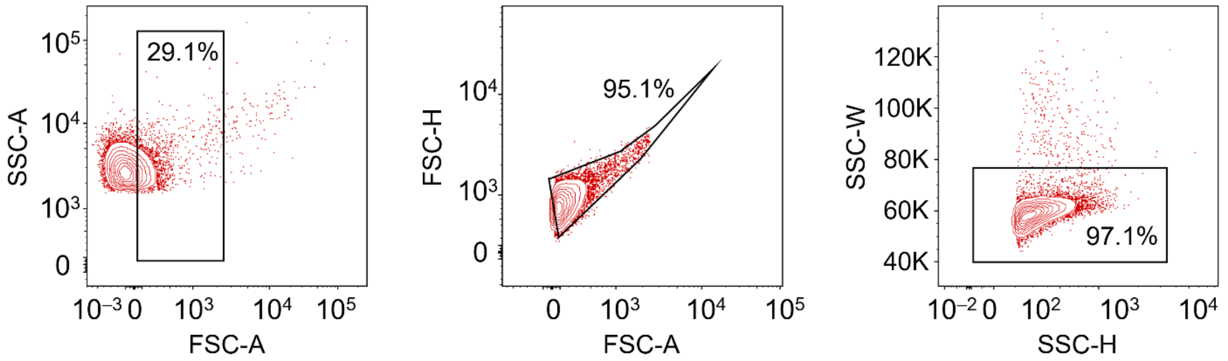

Supplementary Fig. 6. Representative FACS setup (NBDSe 7P+) for the selection of *E. coli* DH10B cells containing the G1RS library. Cell populations were initially identified based on forward scatter (FSC) and side scatter (SSC) to discriminate cells according to size and granularity. Debris was excluded using an SSC area (SSC-A) versus FSC area (FSC-A) density plot, followed by sequential gating using FSC height (FSC-H) versus FSC-A and SSC width (SSC-W) versus SSC height (SSC-H) plots. These morphological assessments revealed no significant differences between the samples.

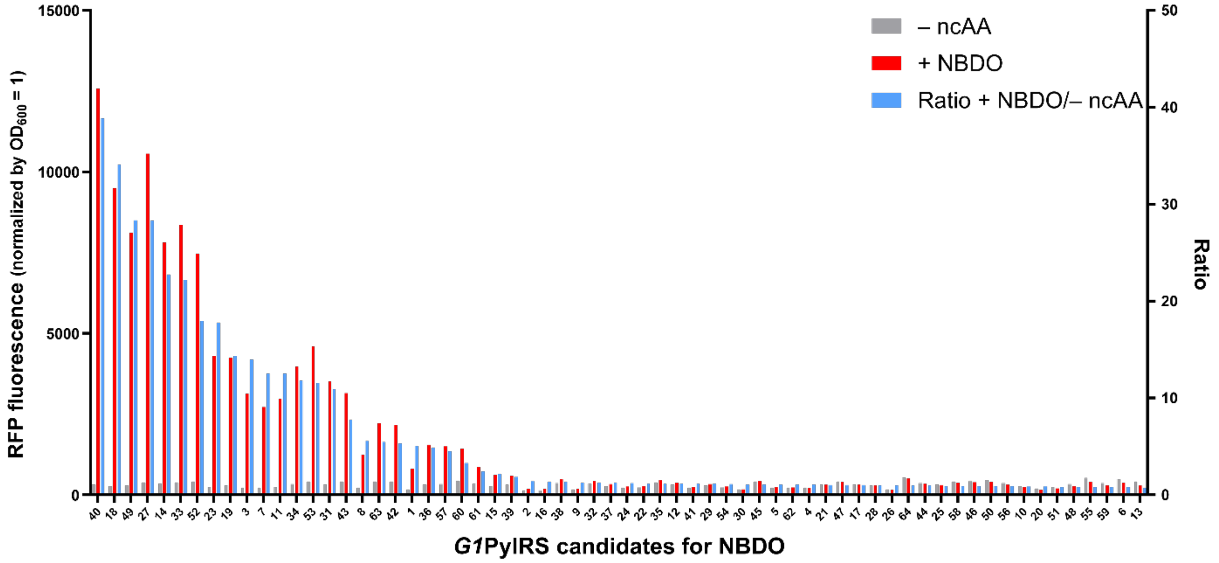

Supplementary Fig. 7. Activity and specificity screen of G1PylRS variants for NBDO incorporation. Cells from the 3.3% fraction with the highest red fluorescence in the final selection round were cultured in 96-well plates with and without 0.5 mM NBDO. Red fluorescence intensity indicative of the readthrough efficiency of the amber-interrupted reporter gene was then measured. The plot presents the colonies ranked in a descending order based on the ratio of red fluorescence in the + NBDO wells compared to the – ncAA wells. This ranking highlights the candidates with the highest activity and specificity for NBDO incorporation.

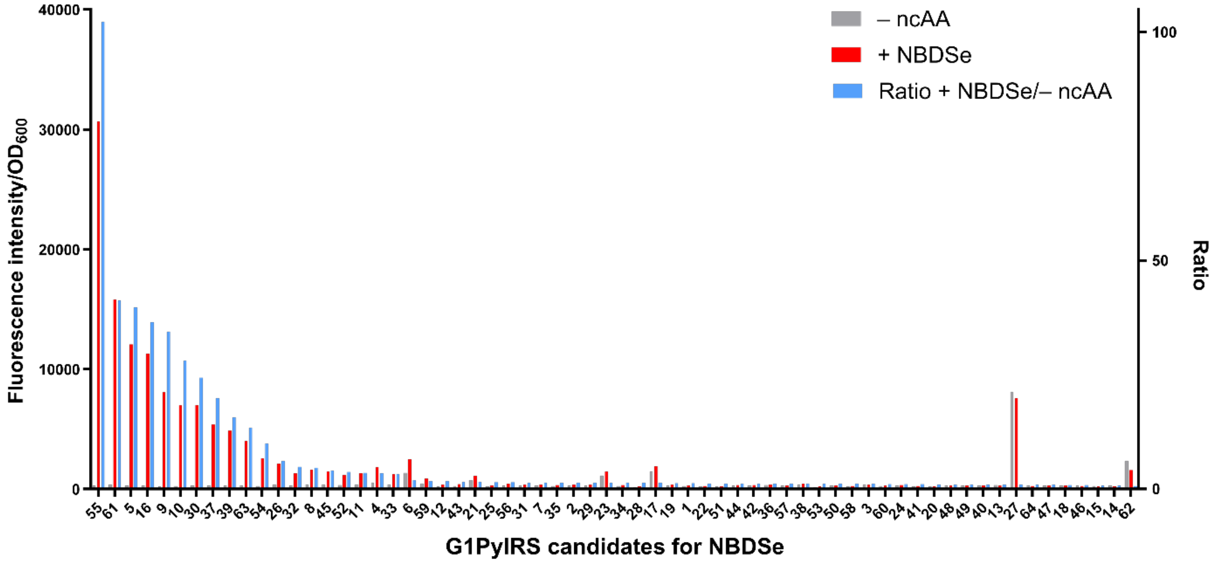

Supplementary Fig. 8. Activity and specificity screen of G1PylRS variants for NBDSe incorporation. Cells from the 3.0% fraction with the highest red fluorescence in the final selection round were cultured in 96-well plates with and without 0.5 mM NBDSe. Red fluorescence intensity indicative of the readthrough efficiency of the amber-interrupted reporter gene was then measured. The plot presents the colonies ranked in a descending order based on the ratio of red fluorescence in the + NBDSe wells compared to the – ncAA wells. This ranking highlights the candidates with the highest activity and specificity for NBDSe incorporation.

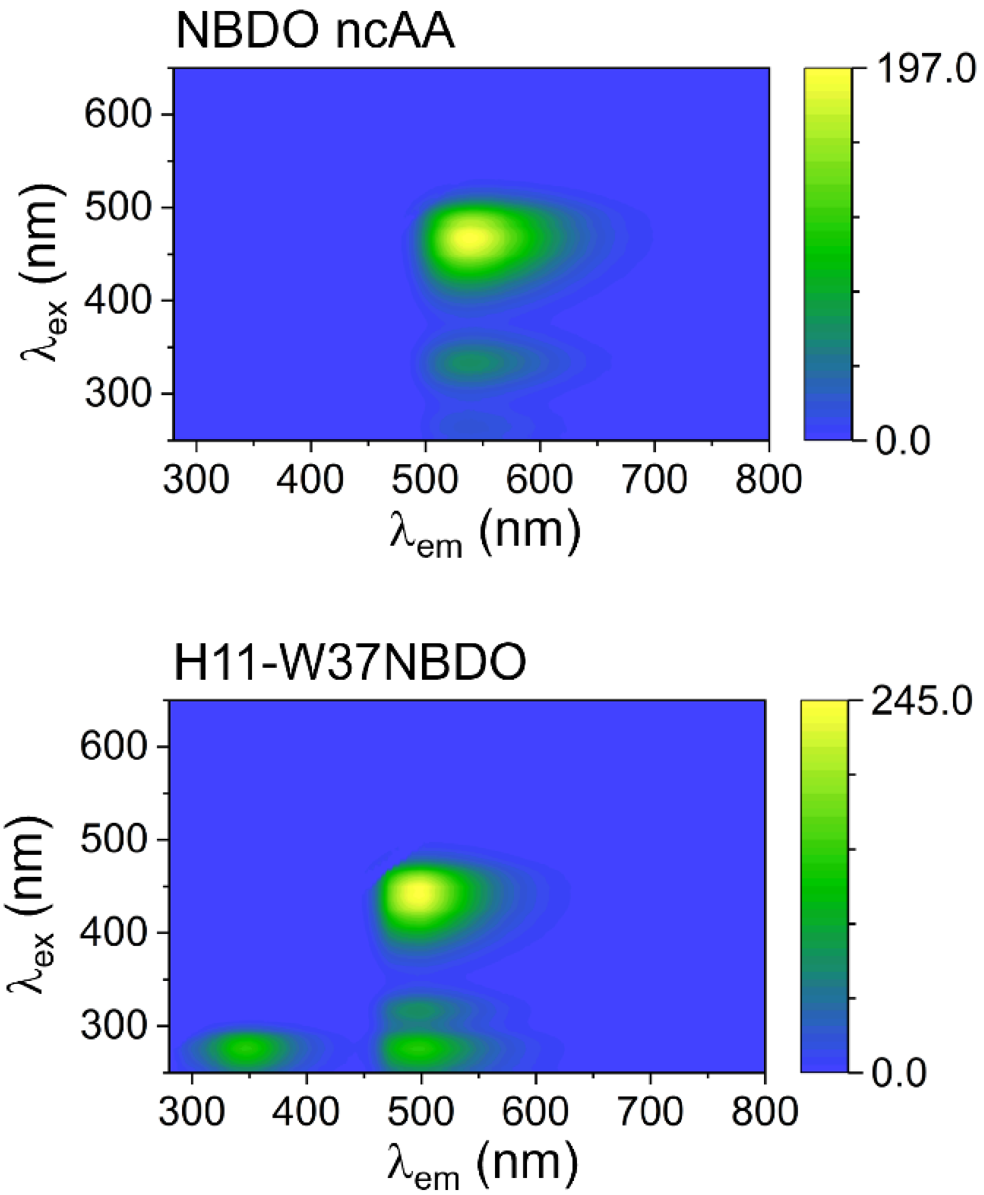

Supplementary Fig. 9. Full spectra of the two-dimensional fluorescence excitation–emission scan of NBDO recorded for the free amino acids and after incorporation in the hydrophobic core of H11, both measured at 10 μM and recorded with the same photomultiplier tube (PMT) detector voltage at 600 V.

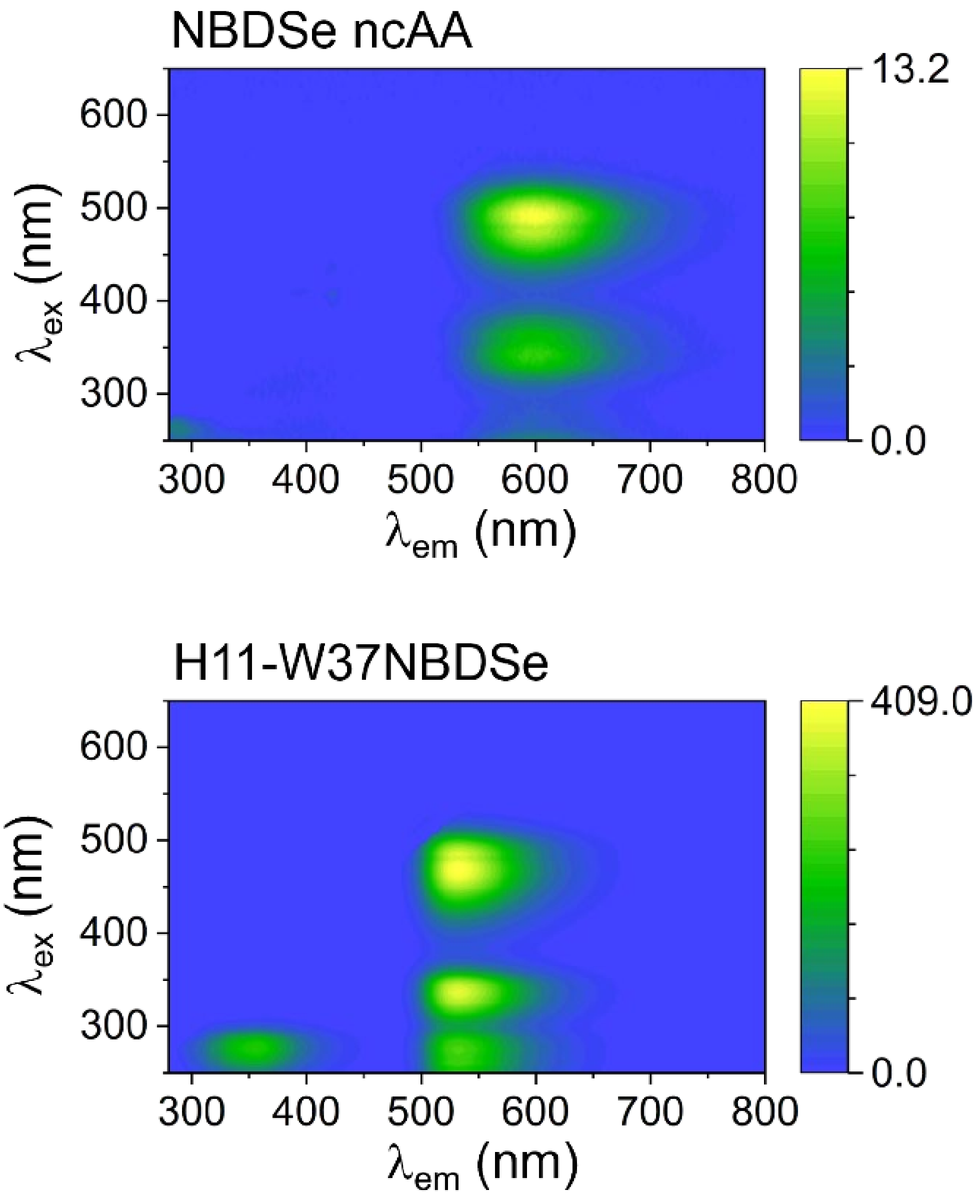

Supplementary Fig. 10. Full spectra of the two-dimensional fluorescence excitation–emission scan of NBDSe recorded for the free amino acids and after incorporation in the hydrophobic core of H11, both measured at 10 μM and recorded with the same PMT detector voltage at 600 V.

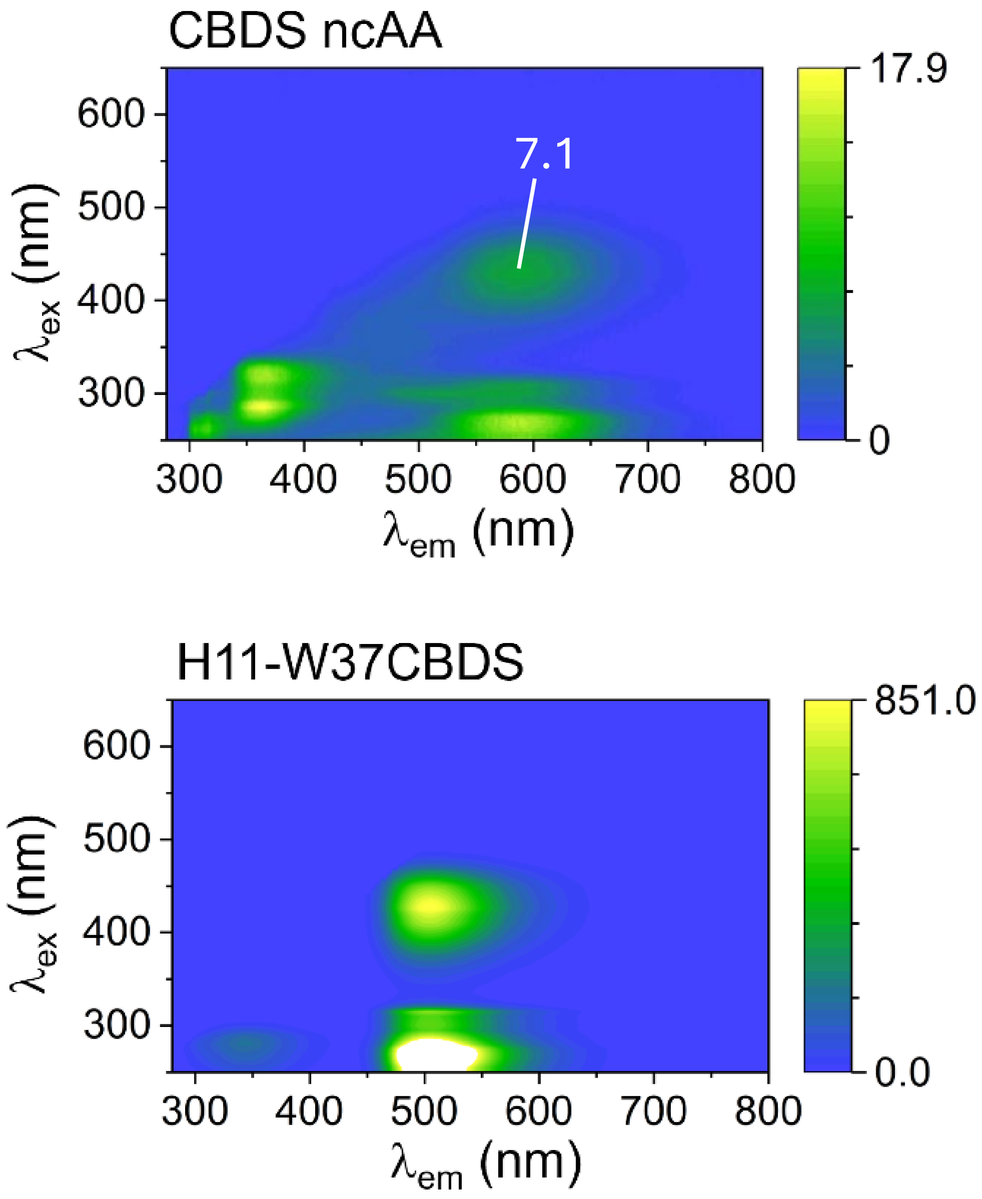

Supplementary Fig. 11. Full spectra of the two-dimensional fluorescence excitation–emission scan of CBDS recorded for the free amino acids and after incorporation in the hydrophobic core of H11, both measured at 10 μM and recorded with the same PMT detector voltage at 600 V. For H11-W37CBDS, the intense excitation feature near 265 nm exceeded the detector range at its emission maximum. For fluorescence intensity comparison, the emission maximum associated with excitation wavelengths above 405 nm was used for both samples.

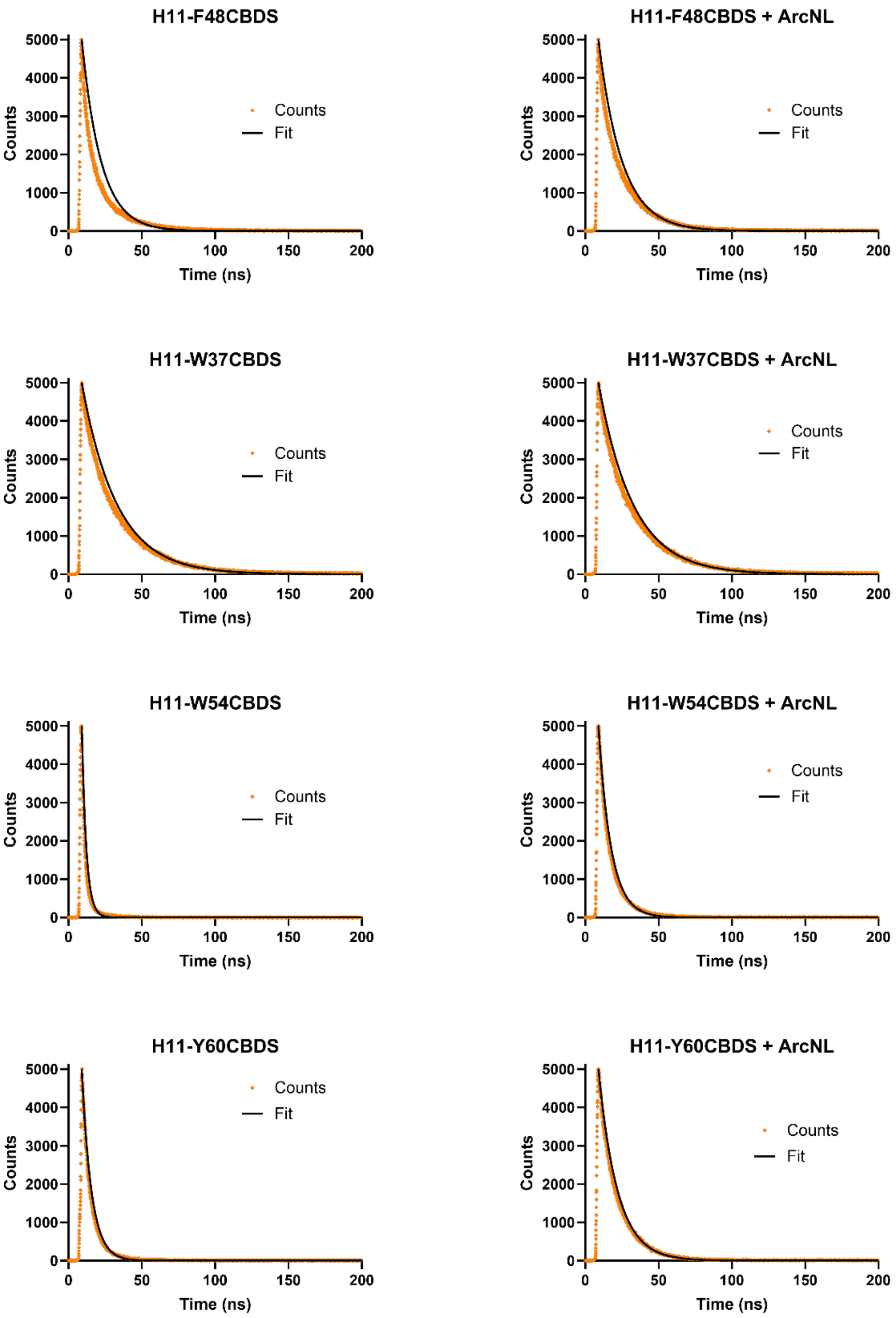

Supplementary Fig. 12. Fluorescence decay histograms of H11 mutants containing CBDS, measured in the presence or absence of ArcNL, are shown as orange symbols together with the corresponding reconvolution fits (black lines). The decay was recorded using TCSPC and fitted using a single-exponential decay model convoluted with the IRF measured under identical experimental conditions.

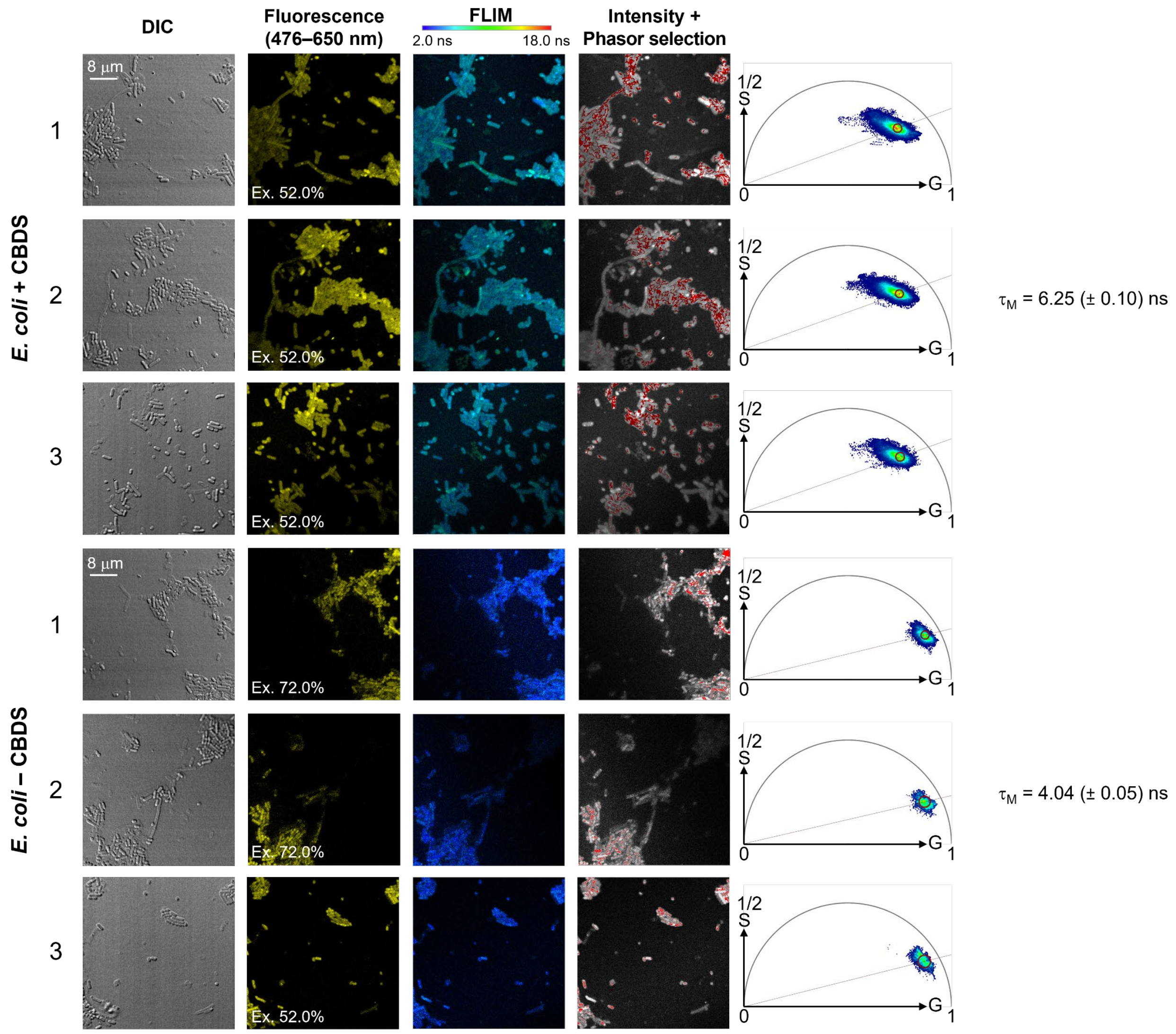

Supplementary Fig. 13. FLIM analysis of *E. coli* B95 cells cultured with or without CBDS supplementation. B95 cells were cultured in the presence (top three rows) or absence (bottom three rows) of CBDS and analyzed by fluorescence lifetime imaging microscopy (FLIM). Three independent imaging experiments were conducted for each condition. From left to right, the columns show transmitted-light images acquired using differential interference contrast (DIC), fluorescence intensity images (yellow), fluorescence lifetime images (pseudocoloured according to the lifetime scale), phasor-selected pixels highlighted in red on the corresponding grayscale images, and the associated phasor plots. Fluorescence was excited at 440 nm using a 20 MHz laser repetition rate, and emission was collected between 476 and 650 nm. The laser excitation power was adjusted independently for each acquisition, as indicated on the corresponding images, to avoid signal saturation during FLIM recording. Fluorescence images were adjusted independently for visualization, with the upper limit of the display range set to the maximum pixel intensity of each image and were not used for quantitative intensity comparison. Phasor populations were selected directly on the phasor plots, and the corresponding photons were mapped back to their spatial positions in the images. The red-highlighted pixels therefore indicate the spatial distribution of the selected phasor population rather than an intensity-based selection. The modulation lifetime ($\tau_M$) was determined independently for each image and averaged across the three images for each culture condition (n = 3 independent imaging experiments). Scale bars, 8 μm; all images are shown at the same magnification.

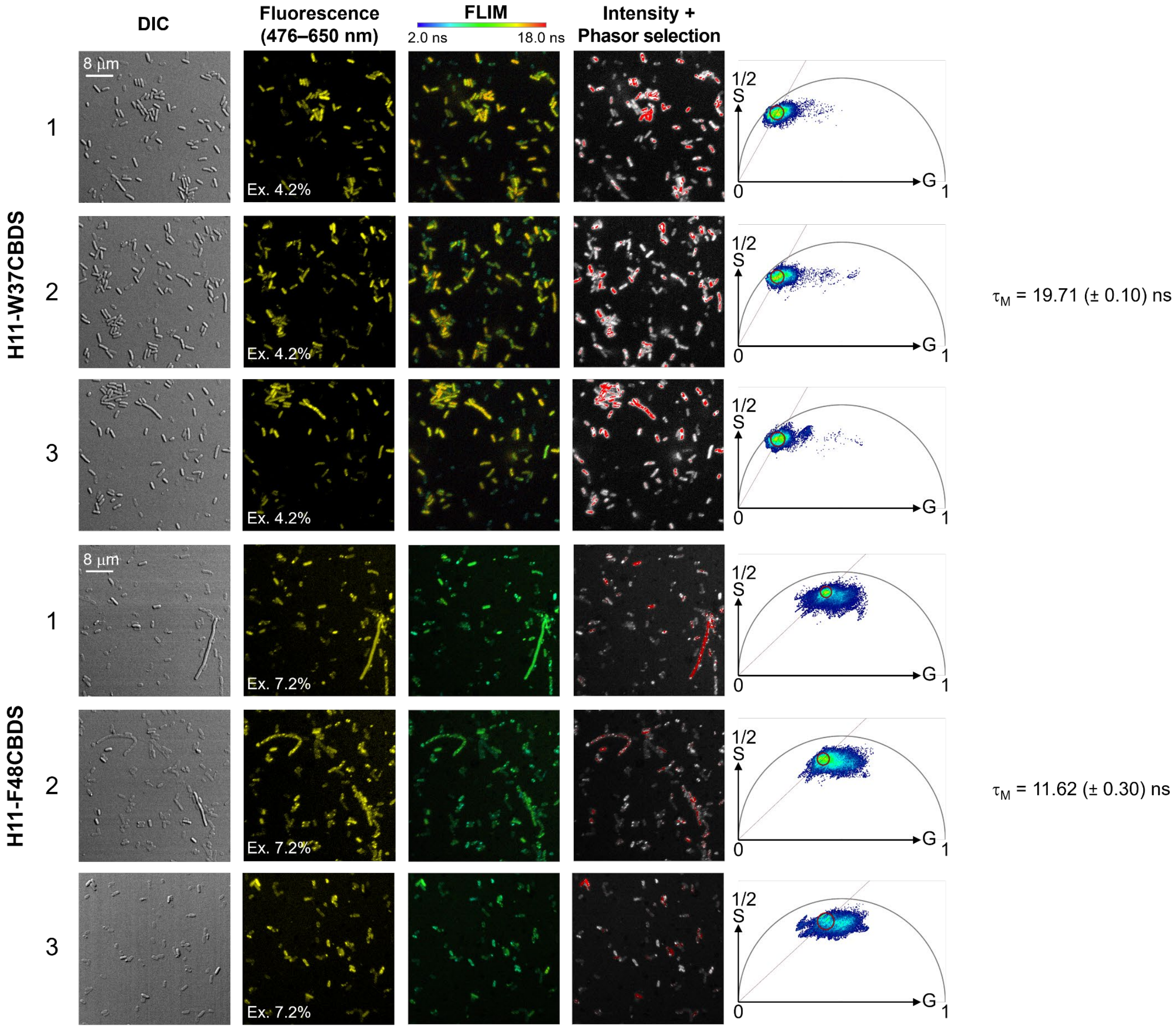

Continued on next page.

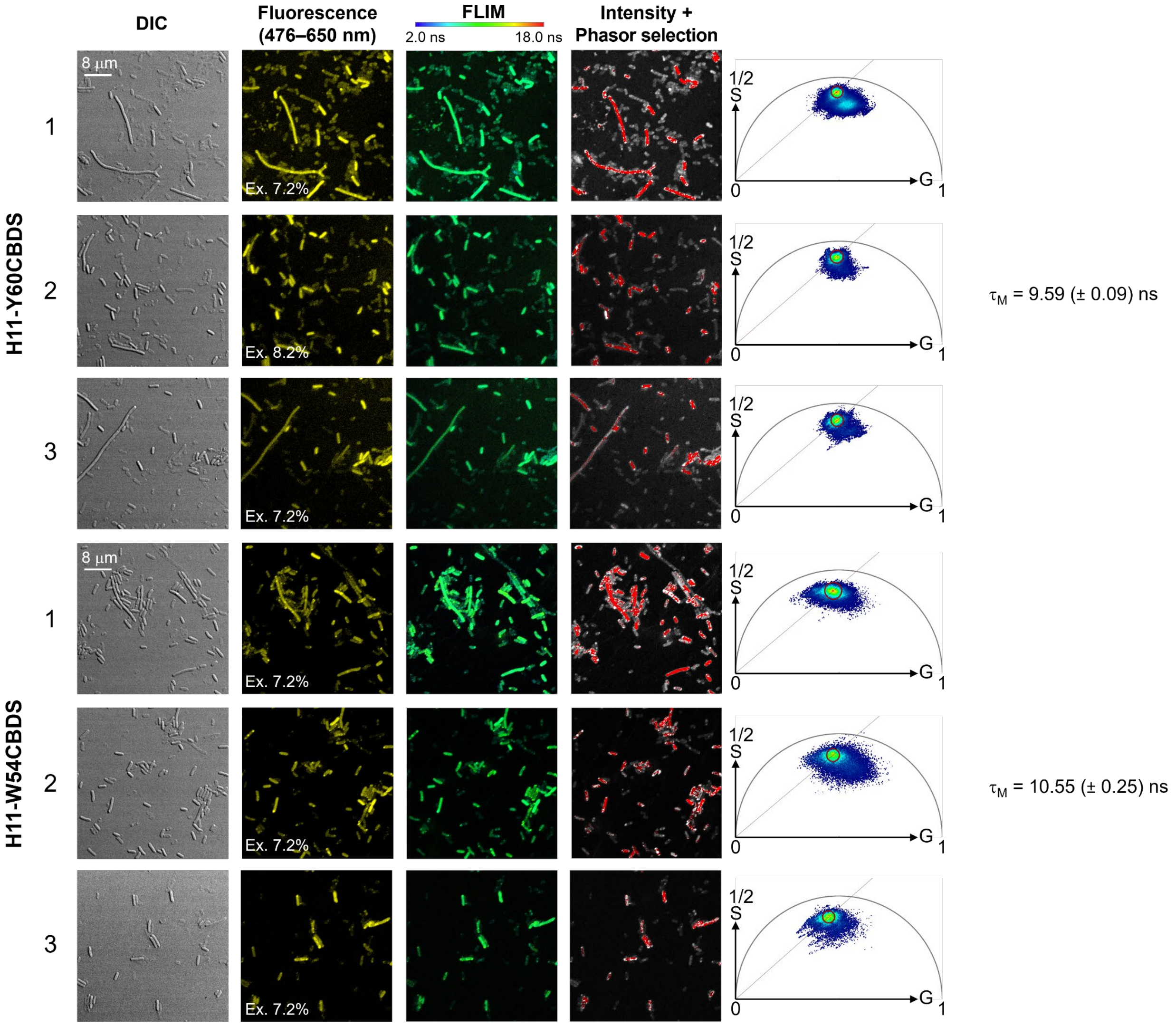

Supplementary Fig. 14. FLIM analysis of *E. coli* B95 cells expressing H11-CBDS mutants. Three independent imaging experiments were conducted for each mutant (H11-W37CBDS, H11-F48CBDS, H11-Y60CBDS, and H11-W54CBDS). Fluorescence was excited at 440 nm using a 20 MHz laser repetition rate, with the laser excitation power adjusted for each acquisition to avoid FLIM detector saturation. Images and phasor analysis are presented as described in Supplementary Fig. 13. $\tau_M$ was determined independently for each image and averaged across the three images for each culture condition (n = 3 independent imaging experiments). Scale bars, 8 µm; all images are shown at the same magnification.

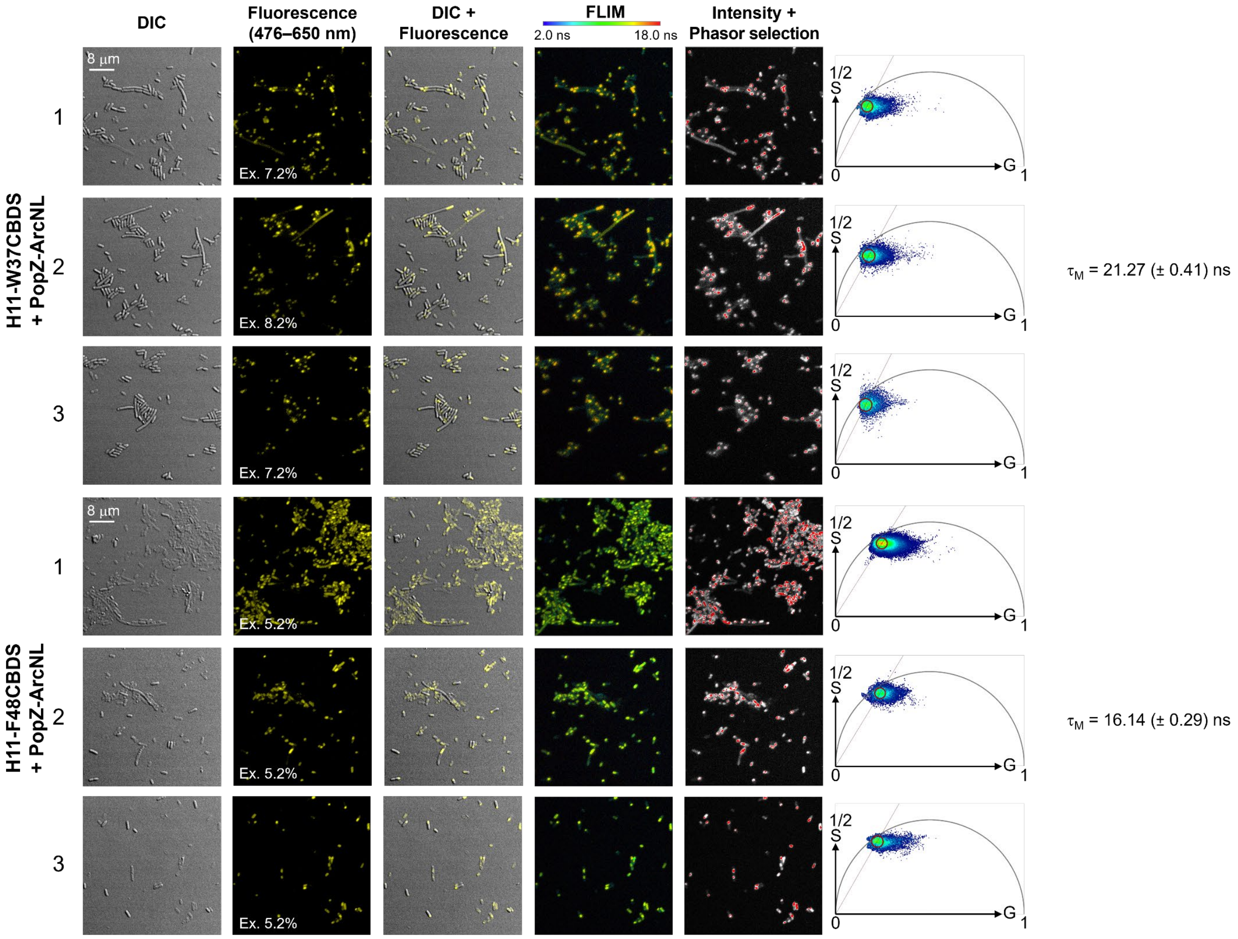

Continued on next page.

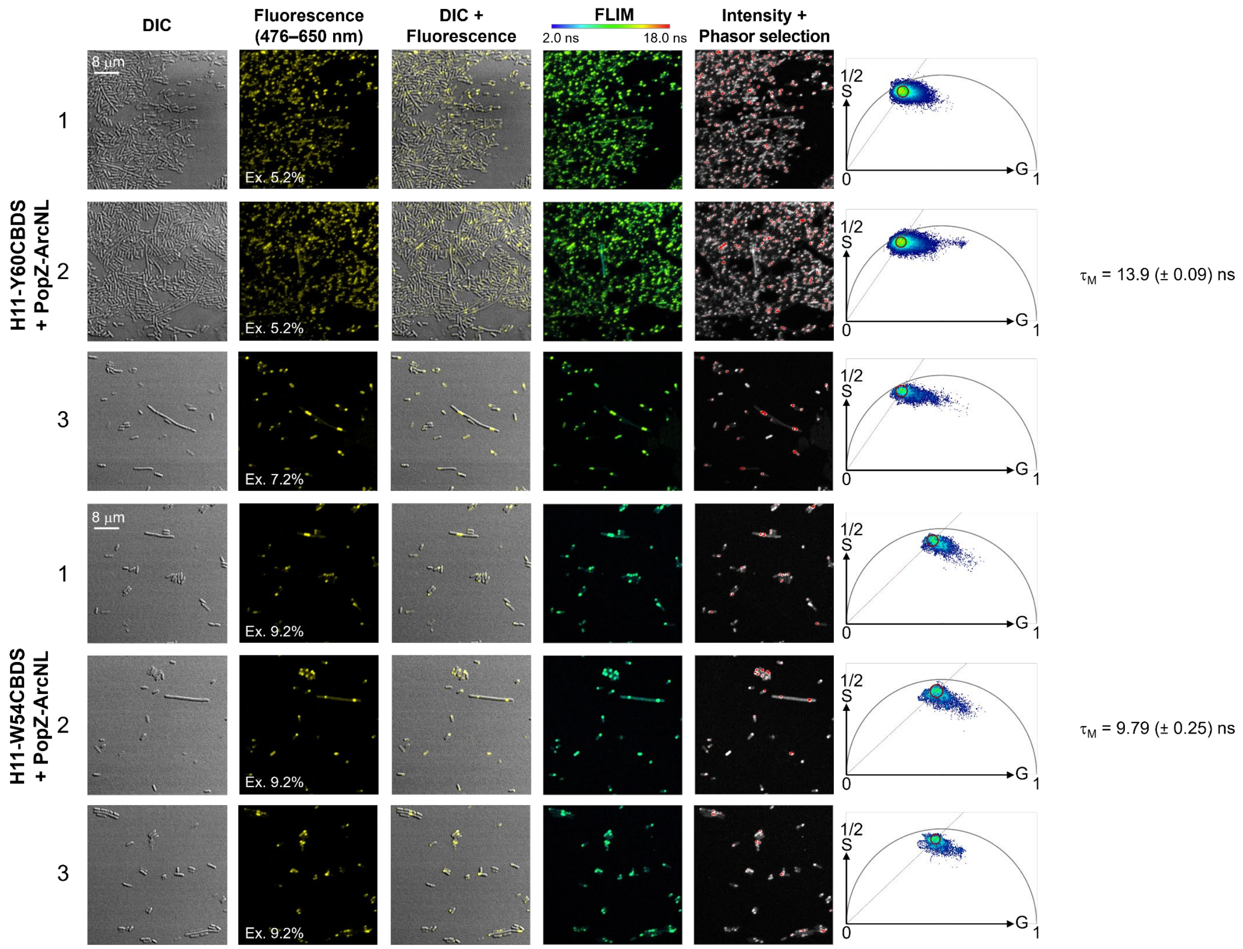

Supplementary Fig. 15. FLIM analysis of *E. coli* B95 cells co-expressing H11-CBDS mutants and PopZ-tagged ArcNL. Three independent imaging experiments were conducted for each mutant (H11-W37CBDS, H11-F48CBDS, H11-Y60CBDS, and H11-W54CBDS). Fluorescence was excited at 440 nm using a 20 MHz laser repetition rate, with the laser excitation power adjusted for each acquisition to avoid FLIM detector saturation. Images and phasor analysis are presented as described in Supplementary Fig. 13. The DIC and fluorescence overlay images show the cellular localization of the fluorescently labelled proteins. $\tau_M$ was determined independently for each image and averaged across the three images for each culture condition (n = 3 independent imaging experiments). Scale bars, 8 µm; all images are shown at the same magnification.

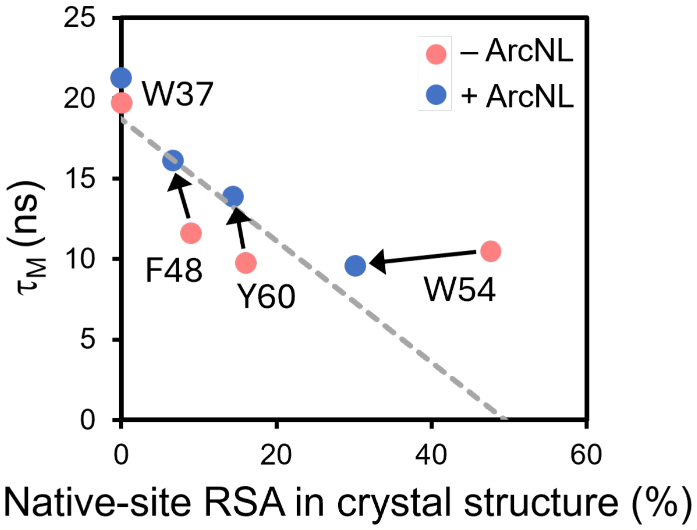

Supplementary Fig. 16. Relationship between fluorescence lifetime and solvent accessibility of H11-CBDS mutants in *E. coli*. $\tau_M$ of H11-CBDS mutants expressed in *E. coli* B95 cells in the absence (red) or presence (blue) of co-expressed PopZ-ArcNL, plotted against the relative solvent-accessible surface area (RSA) of the corresponding native residues, calculated from the crystal structures of H11 (PDB 8QF3 and 8QF4).[3] The plot allows comparison with the corresponding measurements in PBS (Fig. 3f) to assess the effects of the cellular environment on the fluorescence lifetimes of CBDS at different incorporation sites. H11-W54-CBDS deviated from the linear relationship observed for the other mutants in the absence of ArcNL, potentially reflecting nonspecific interactions with cellular components in the absence of its native binding partner. Linear regression of the data excluding H11-W54-CBDS (gray dashed line) yielded $\tau_M = -0.38 \times \text{RSA} + 18.68$ ($R^2 = 0.73$), compared with $\tau_M = -0.21 \times \text{RSA} + 17.27$ ($R^2 = 0.54$) when H11-W54-CBDS was included.

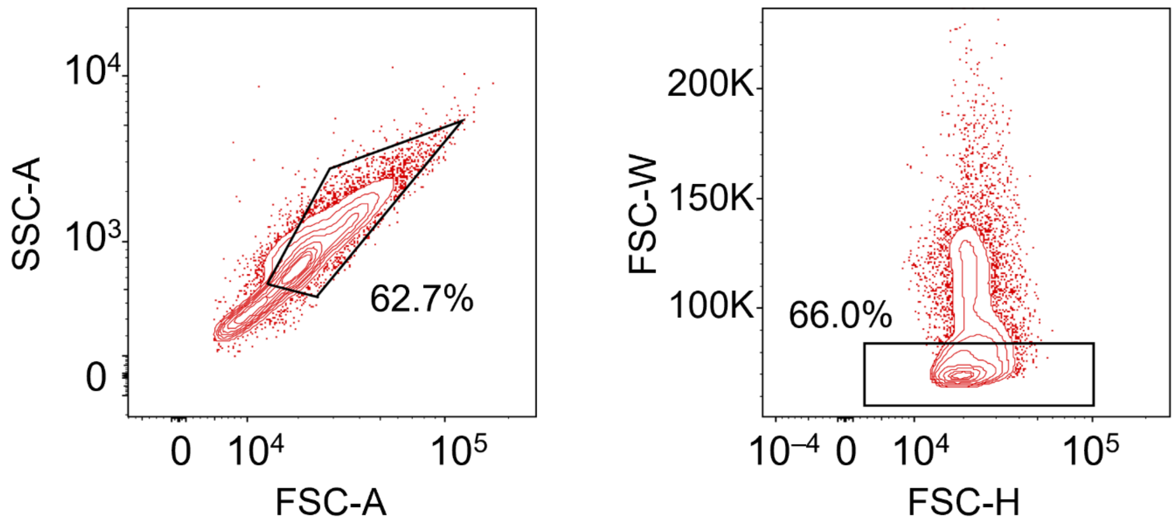

Supplementary Fig. 17. FACS setup for the amber readthrough efficiency test of G1NBDO53 in genome modified *S. cerevisiae* BY4741. Debris was excluded using an SSC-A versus FSC-A density plot, followed by sequential gating based on FSC-W versus FSC-H to exclude budding cells. Blue and red fluorescence were detected using 405 nm and 560 nm excitation lasers and measured using 450/50 and 610/20 nm bandpass filters, respectively.

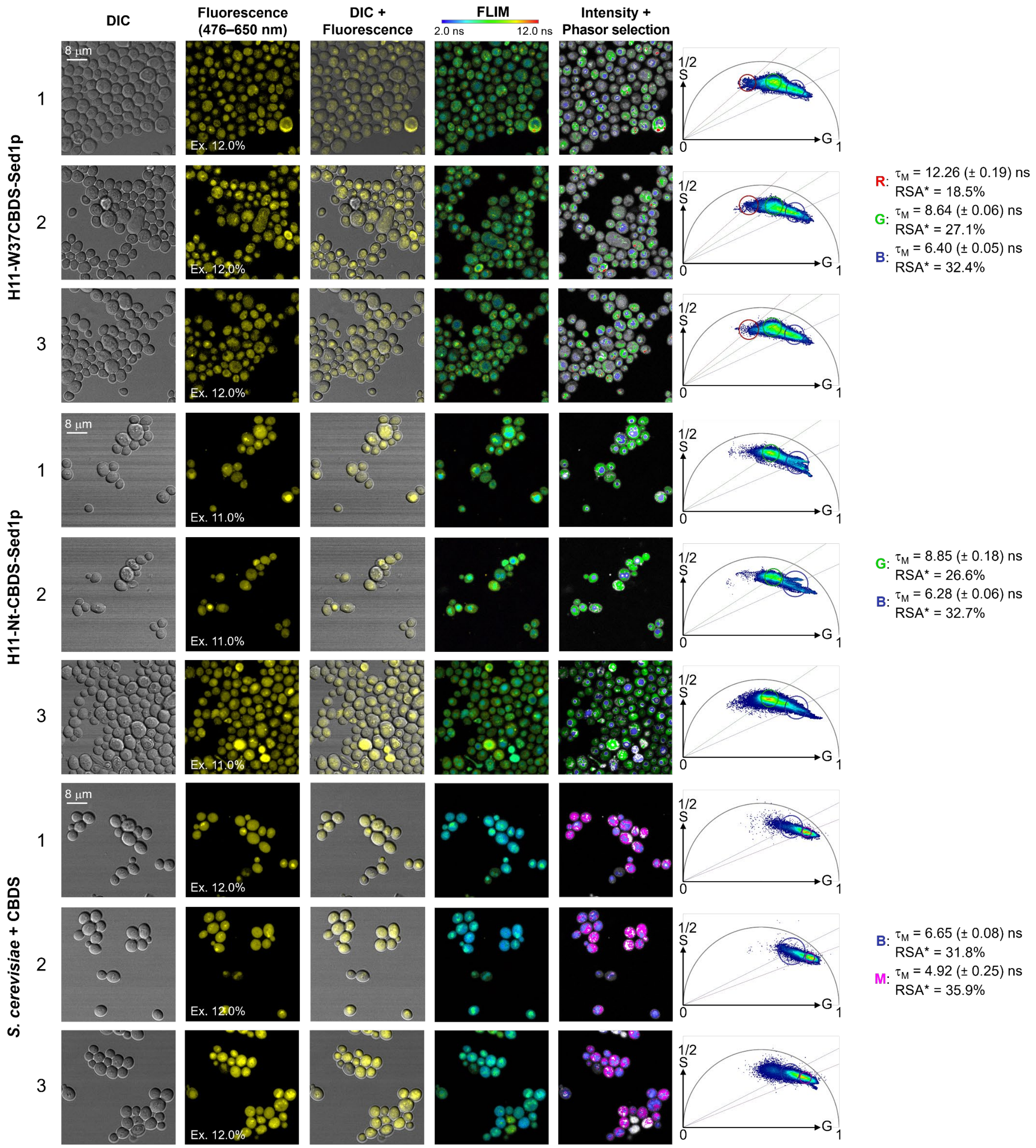

Supplementary Fig. 18. FLIM analysis of *S. cerevisiae* BY4741 cells expressing CBDS-incorporated proteins or cultured with CBDS alone as a control. Fluorescence was excited at 440 nm using a 20 MHz laser repetition rate, with the laser excitation power adjusted for each acquisition. Images and phasor analysis are presented as described in Supplementary Fig. 13–15. $\tau_M$ for each cellular context was determined independently for each image and averaged across three images for each culture condition (n = 3 independent imaging experiments). Scale bars, 8 μm; all images are shown at the same magnification.

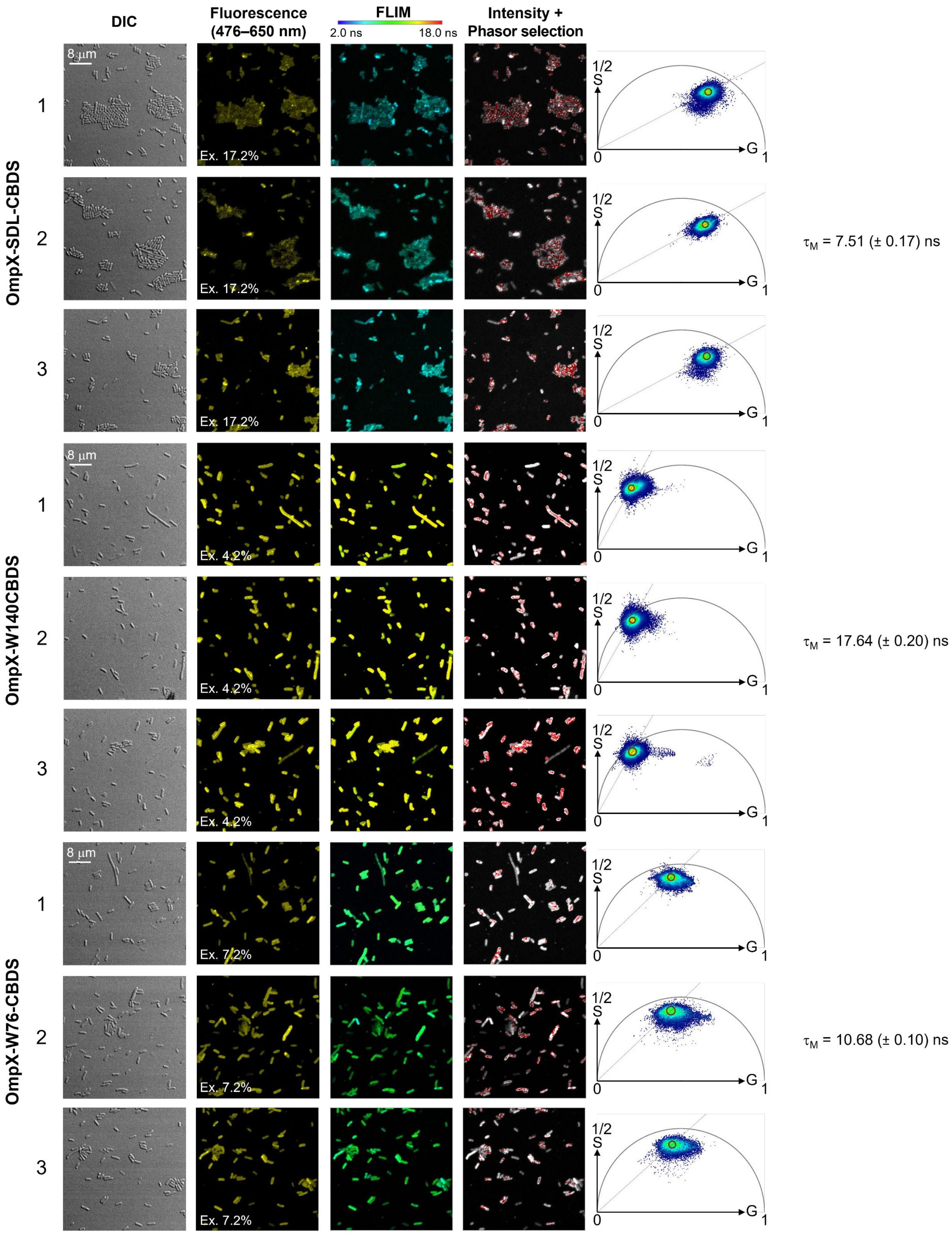
DIC
Fluorescence (476–650 nm)
FLIM
2.0 ns
18.0 ns
Intensity + Phasor selection
8 µm
Ex. 17.2%
Ex. 4.2%
Ex. 7.2%
OmpX-SDL-CBDS
OmpX-W140CBDS
OmpX-W76-CBDS
1
2
3
1/2
S
G
0
1
τ_M = 7.51 (± 0.17) ns
τ_M = 17.64 (± 0.20) ns
τ_M = 10.68 (± 0.10) ns

Supplementary Fig. 19. FLIM analysis of *E. coli* B95 cells expressing OmpX-CBDS mutants. Three independent imaging experiments were conducted for each mutant (OmpX-SDL, OmpX-W140, and OmpX-W76). Fluorescence was excited at 440 nm using a 20 MHz laser repetition rate, with the laser excitation power adjusted for each acquisition to avoid FLIM detector saturation. Images and phasor analysis are presented as described in Supplementary Fig. 13. $\tau_M$ was determined independently for each image and averaged across the three images for each culture condition (n = 3 independent imaging experiments). Scale bars, 8 μm; all images are shown at the same magnification.

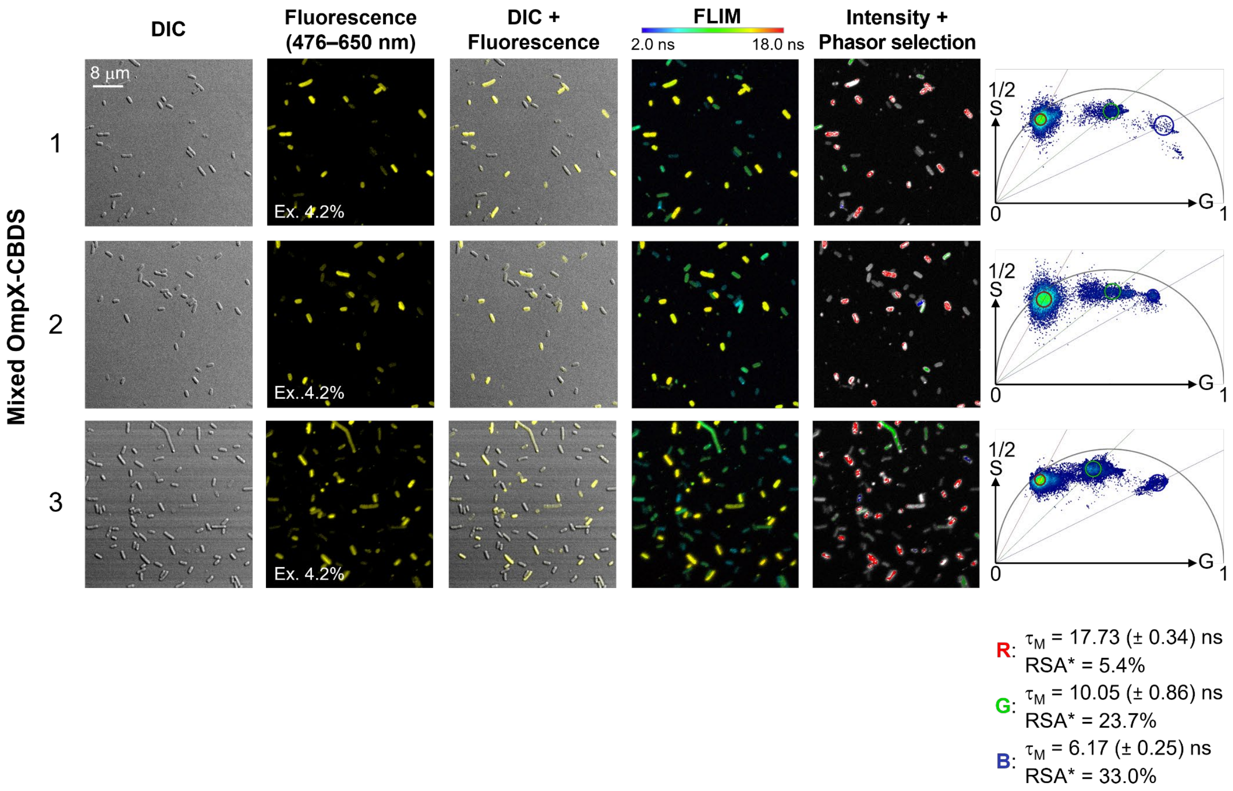

Supplementary Fig. 20. FLIM analysis of *E. coli* B95 cells expressing three different OmpX-CBDS mutants in co-culture. Three independent imaging experiments were conducted. Fluorescence was excited at 440 nm using a 20 MHz laser repetition rate, with the laser excitation power adjusted for each acquisition to avoid FLIM detector saturation. Images and phasor analysis are presented as described in Supplementary Fig. 13. $\tau_M$ for each OmpX-CBDS mutant population was determined independently for each image and averaged across three images for each culture condition (n = 3 independent imaging experiments). Scale bars, 8 μm; all images are shown at the same magnification.

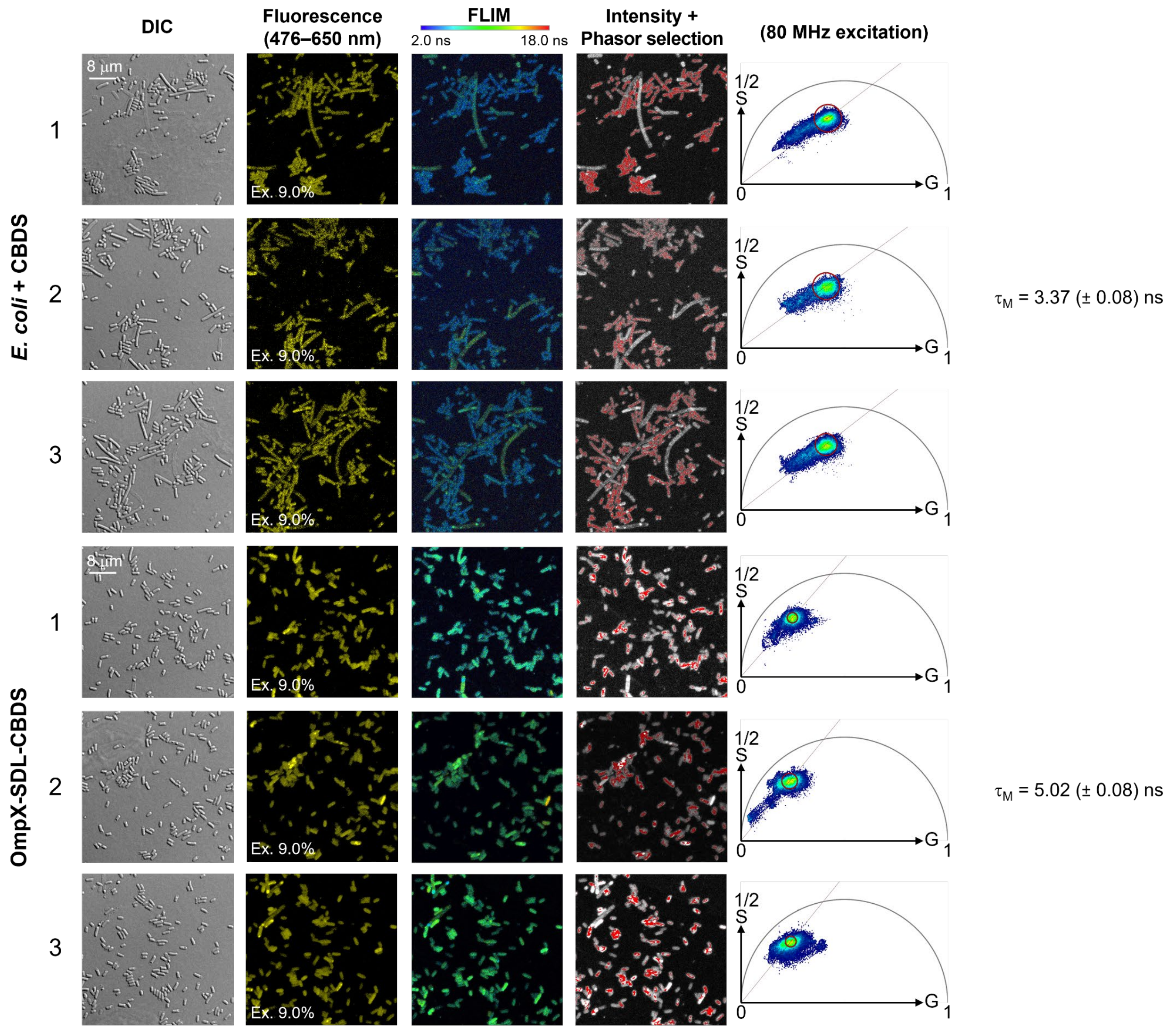

Continued on next page.

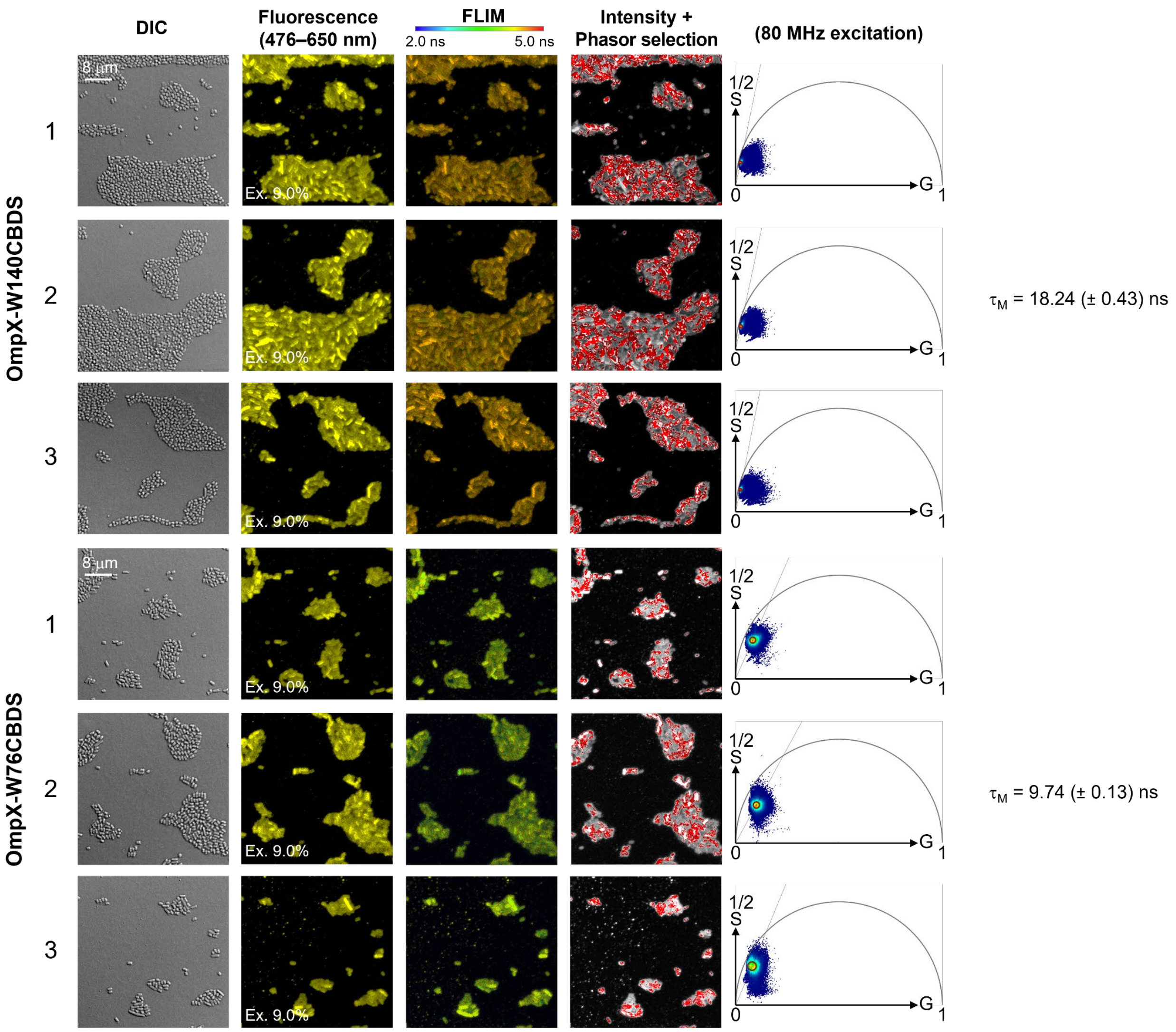

Supplementary Fig. 21. FLIM analysis of *E. coli* B95 cells expressing OmpX-CBDS mutants using an 80 MHz laser repetition rate. Three independent imaging experiments were conducted for each mutant (OmpX-SDL, OmpX-W140 and OmpX-W76) and for cells cultured with CBDS alone as a control. Fluorescence was excited at 440 nm using an 80 MHz laser repetition rate, with the laser excitation power fixed at 9% for all acquisitions to allow direct comparison of fluorescence intensities without saturating the FLIM detector. Images and phasor analysis are presented as described in Supplementary Fig. 13. $\tau_M$ was determined independently for each image and averaged across the three images for each culture condition (n = 3 independent imaging experiments). The complete fluorescence decay of OmpX-W140CBDS could not be captured within the 12.5 ns interpulse interval at an 80 MHz laser repetition rate. Scale bars, 8 µm; all images are shown at the same magnification.

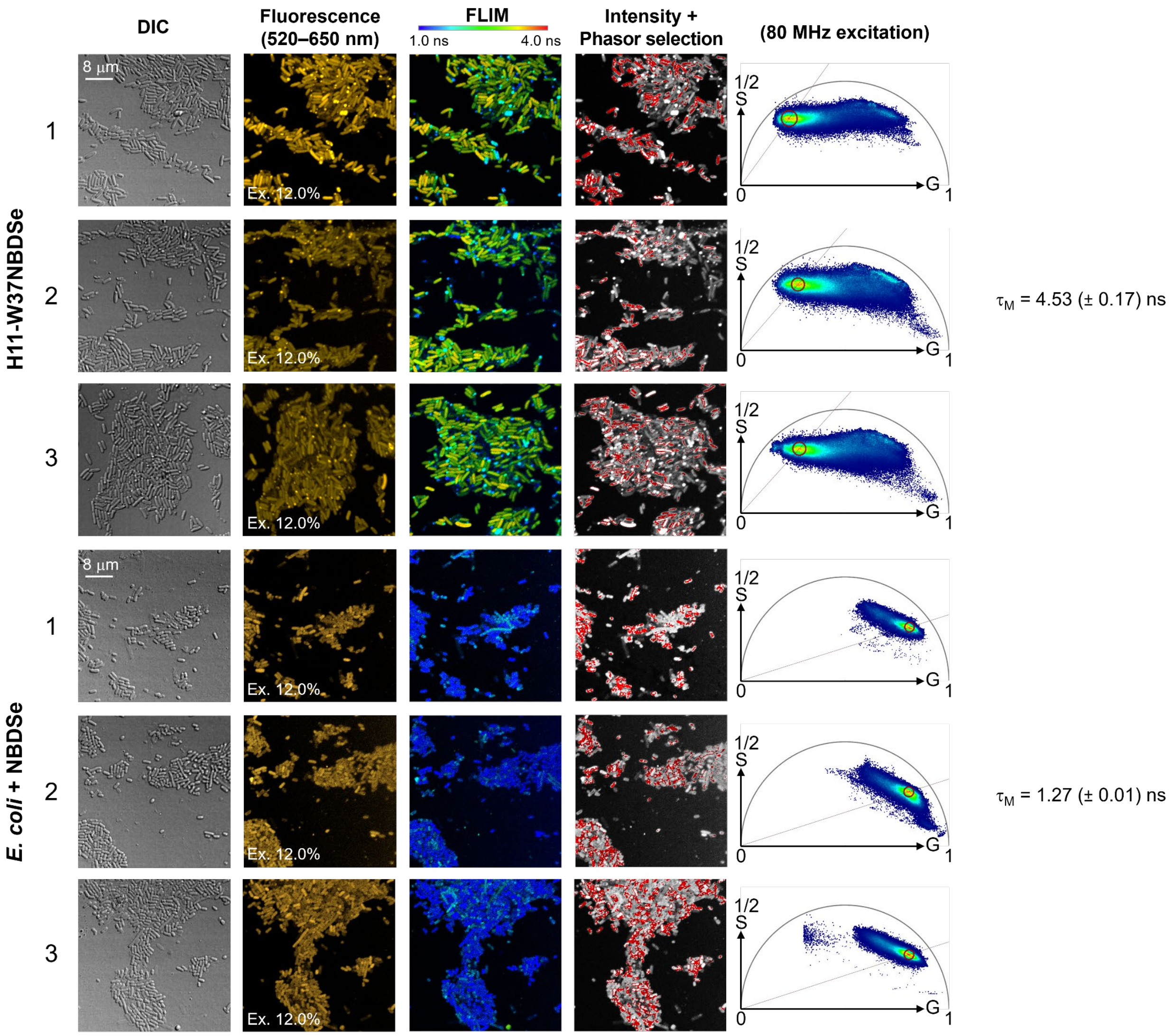

Supplementary Fig. 22. FLIM analysis of *E. coli* B95 cells expressing H11-W37NBDSe using an 80 MHz laser repetition rate. Three independent imaging experiments were conducted for cells expressing H11-W37NBDSe and for cells cultured with NBDSe alone as a control. Fluorescence was excited at 500 nm using an 80 MHz laser repetition rate, with emission collected between 520 and 650 nm. The laser excitation power was fixed at 12% for all acquisitions without saturating the FLIM detector. Images and phasor analysis are presented as described in Supplementary Fig. 13. $\tau_M$ was determined independently for each image and averaged across the three images for each culture condition (n = 3 independent imaging experiments). Scale bars, 8 μm; all images are shown at the same magnification.

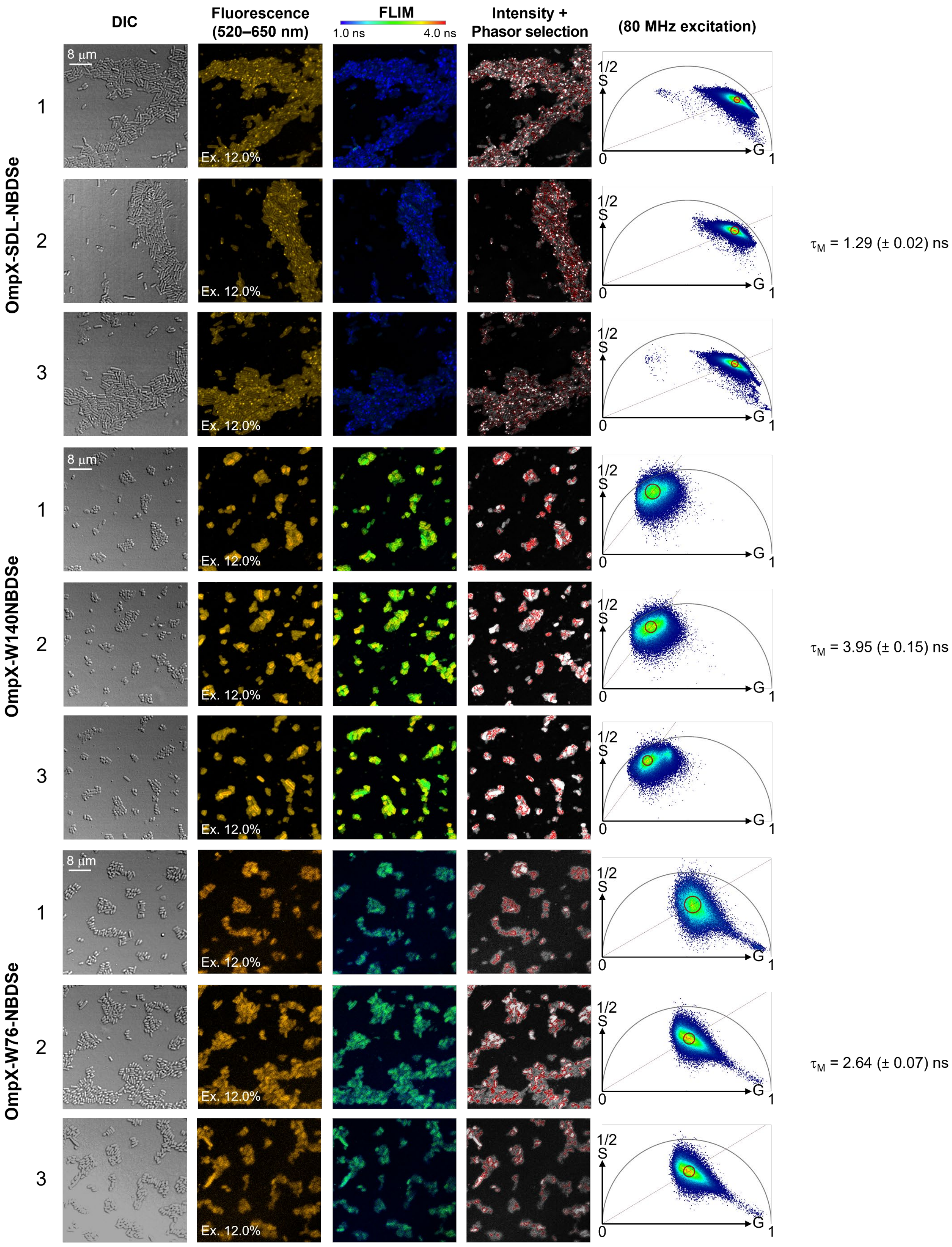

DIC
Fluorescence (520–650 nm)
FLIM
1.0 ns
4.0 ns
Intensity + Phasor selection
(80 MHz excitation)
OmpX-SDL-NBDSe
OmpX-W140NBDSe
OmpX-W76-NBDSe
8 μm
Ex. 12.0%
1/2
S
G
0
1
$\tau_M$ = 1.29 (± 0.02) ns
$\tau_M$ = 3.95 (± 0.15) ns
$\tau_M$ = 2.64 (± 0.07) ns

Supplementary Fig. 23. FLIM analysis of *E. coli* B95 cells expressing OmpX-NBDSe mutants using an 80 MHz laser repetition rate. Three independent imaging experiments were conducted for each mutant. Fluorescence was excited at 500 nm using an 80 MHz laser repetition rate, with emission collected between 520 and 650 nm. The laser excitation power was fixed at 12% for all acquisitions without saturating the FLIM detector. Images and phasor analysis are presented as described in Supplementary Fig. 13. $\tau_M$ was determined independently for each image and averaged across the three images for each culture condition (n = 3 independent imaging experiments). Scale bars, 8 μm; all images are shown at the same magnification.

**Supplementary tables**

Supplementary Table 1. Characteristic properties of fluorescent amino acids.

| | $\lambda_{ex}$ (nm)[a] | $\lambda_{em}$ (nm)[a] | Van der Waals volume of the side chain ($Å^3$)[b] |
|---|---|---|---|
| Trp | 280 | 350 | 123.18 |
| 4-CN-Trp | 305 | 424 | 146.2 |
| 6-CN-Trp | 280 | 380 | 146.2 |
| 7-CN-Trp | 305 | 416 | 146.2 |
| 6-Ac-Trp | 301 | 470 | 163.93 |
| Hco | 340 | 460 | 170.51 |
| Acd | 386 | 420 | 192.29 |
| Anap | 350 | 490 | 195.89 |
| Aad | 420 | 527 | 203.29 |
| DanAla | 340 | 550 | 236.83 |
| AnapTh | 415 | 630 | 269.72 |
| CouLys | 340 | 460 | 282.95 |
| NBDO | 470 | 538 | 145.31 |
| NBDSe | 490 | 600 | 152.09 |
| CBDS | 430 | 592 | 152.11 |

[a] Emission wavelengths listed are the corresponding emission maxima measured in aqueous solution; the excitation wavelengths listed are those used for the measurements;

[b] Van der Waals volume calculated using the Slonimskii method,[4, 5] considering the side chain from the β-position onward.

Supplementary Table 2. Fluorescence lifetimes of CBDS in different solvents measured using a time-resolved spectrofluorometer.

| | $\tau_{fit}$ (ns) | ε (25°C) | $E_T(30)$ (kcal/mol) | Viscosity (25°C; cp) |
|---|---|---|---|---|
| Water | 1.02 | 80.1 | 63.1 | 0.89 |
| Methanol | 4.53 | 32.7 | 55.4 | 0.54 |
| Ethanol | 7.21 | 24.55 | 51.9 | 1.07 |
| DMSO | 10.48 | 46.68 | 45.1 | 1.99 |
| Acetone | 16.03 | 20.7 | 42.2 | 0.31 |
| Chloroform | 16.79 | 4.81 | 39.1 | 0.53 |
| DCM | 21.49 | 8.93 | 40.7 | 0.41 |
| 1:1 EG:Water | 1.89 | - | - | 3.4 |
| 9:1 EG:Water | 3.11 | - | - | 6.4 |

Supplementary Table 3. Mutations found in six G1PylRS variants selected for activity with NBDO and NBDSe.

| RS Variants | Randomized Sites | | | | | | |
|---|---|---|---|---|---|---|---|
| G1PylRS wild-type | L124 | Y125 | N165 | V167 | Y204 | A221 | W237 |
| G1NBDO15 | A | F | N | L | Y | G | Y |
| G1NBDO18 | G | F | N | L | Y | G | Y |
| G1NBDO19 | M | F | N | V | Y | G | Y |
| G1NBDO53[a] | M | Y | N | V | Y | G | Y |
| G1NBDSe09[a] | C | F | N | G | W | G | Y |
| G1NBDSe30 | S | L | N | V | W | G | Y |

[a] The mutants G1NBDO53 and G1NBDSe09 was used for all following applications.

Supplementary Table 4. DNA and corresponding amino acid sequences of the proteins used in the current work.[a]

| Protein | DNA sequence | Amino acid sequence |
|---|---|---|
| G1NBDO53 | ATGGTGGTGAAATTTACCGATAGCCAGATTCAGCATCTGATG<br>GAATATGGTGATAATGATTGGAGCGAAGCCGAATTTGAAGAT<br>GCAGCAGCACGTGATAAAGAATTTAGCAGCCAGTTTAGCAAA<br>CTGAAAAGCGCCAATGATAAAGGCCTGAAAGATGTTATTGCA<br>AATCCGCGTAATGATCTGACCGATCTGGAAAACAAAATTCGC<br>GAAAAACTGGCAGCCCGTGGTTTTATTGAAGTTCATACCCCG<br>ATTTTTGTGAGCAAAAGCGCACTGGCAAAAATGACCATTACC<br>GAAGATCATCCGCTGTTCAAACAGGTGTTTTGGATTGATGATA<br>AACGTGCACTGCGTCCGATGCATGCAATGAATATGTATAAAG<br>TTATGCGTGAACTGCGCGATCATACCAAAGGTCCGGTTAAAA<br>TCTTTGAAATTGGTAGCTGCTTTCGCAAAGAAAGCAAAAGCAG<br>TACCCATCTGGAAGAATTTACCATGCTGAACCTGGTAGAAAT<br>GGGTCCTGATGGTGATCCGATGGAACATCTGAAAATGTATATT<br>GGCGATATCATGGATGCCGTTGGTGTTGAATATACCACCAGT<br>CGTGAAGAATCAGATGTTTATGTTGAAACCCTGGACGTGGAA<br>ATTAATGGCACCGAAGTTGCAAGCGGTGGGGTTGGTCCGCA<br>TAAACTGGATCCGGCACATGATGTGCATGAACCGTATGCAG<br>GTATTGGTTTTGGTCTGGAACGTCTGCTGATGCTGAAAAATGG<br>TAAAAGCAATGCACGCAAAACCGGCAAAAGTATTACCTATCT<br>GAATGGCTACAAACTGGATTAA | MVVKFTDSQIQHLM<br>EYGDNDWSEAEFE<br>DAAARDKEFSSQFS<br>KLKSANDKGLKDVI<br>ANPRNDLTDLENKI<br>REKLAARGFIEVHTP<br>IFVSKSALAKMTITE<br>DHPLFKQVFWIDDK<br>RALRPMHAMNMYK<br>VMRELRDHTKGPV<br>KIFEIGSCFRKESKS<br>STHLEEFTMLNLVE<br>MGPDGDPMEHLK<br>MYIGDIMDAVGVEY<br>TTSREESDVYVETLD<br>VEINGTEVASGGVG<br>PHKLDPAHDVHEP<br>YAGIGFGLERLLML<br>KNGKSNARKTGKSI<br>TYLNGYKLD |
| G1NBDSe09 | ATGGTGGTGAAATTTACCGATAGCCAGATTCAGCATCTGATG<br>GAATATGGTGATAATGATTGGAGCGAAGCCGAATTTGAAGAT<br>GCAGCAGCACGTGATAAAGAATTTAGCAGCCAGTTTAGCAAA | MVVKFTDSQIQHL<br>MEYGDNDWSEAE<br>FEDAAARDKEFSS |

| | | |
|---|---|---|
| | CTGAAAAGCGCCAATGATAAAGGCCTGAAAGATGTTATTGCA<br>AATCCGCGTAATGATCTGACCGATCTGGAAAACAAAATTCGC<br>GAAAAACTGGCAGCCCGTGGTTTTATTGAAGTTCATACCCCG<br>ATTTTTGTGAGCAAAAGCGCACTGGCAAAAATGACCATTACC<br>GAAGATCATCCGCTGTTCAAACAGGTGTTTTGGATTGATGATA<br>AACGTGCACTGCGTCCGATGCATGCAATGAATTGTTTTAAAGT<br>TATGCGTGAACTGCGCGATCATACCAAAGGTCCGGTTAAAAT<br>CTTTGAAATTGGTAGCTGCTTTCGCAAAGAAAGCAAAAGCAGT<br>ACCCATCTGGAAGAATTTACCATGCTGAACCTGGGCGAAATG<br>GGTCCTGATGGTGATCCGATGGAACATCTGAAAATGTATATTG<br>GCGATATCATGGATGCCGTTGGTGTTGAATATACCACCAGTC<br>GTGAAGAATCAGATGTTTGGGTTGAAACCCTGGACGTGGAAA<br>TTAATGGCACCGAAGTTGCAAGCGGTGGGGTTGGTCCGCAT<br>AAACTGGATCCGGCACATGATGTGCATGAACCGTATGCAGG<br>TATTGGTTTTGGTCTGGAACGTCTGCTGATGCTGAAAAATGGT<br>AAAAGCAATGCACGCAAAACCGGCAAAAGTATTACCTATCTG<br>AATGGCTACAAACTGGATTAA | QFSKLKSANDKGL<br>KDVIANPRNDLTD<br>LENKIREKLAARGF<br>IEVHTPIFVSKSALA<br>KMTITEDHPLFKQV<br>FWIDDKRALRPMH<br>AMNCFKVMRELR<br>DHTKGPVKIFEIGS<br>CFRKESKSSTHLEE<br>FTMLNLGEMGPD<br>GDPMEHLKMYIG<br>DIMDAVGVEYTTS<br>REESDVWVETLDV<br>EINGTEVASGGVG<br>PHKLDPAHDVHEP<br>YAGIGFGLERLLML<br>KNGKSNARKTGKS<br>ITYLNGYKLD |
| FKBP12-<br>W59TAG | ATGGGTGTTCAGGTTGAAACCATTAGTCCTGGTGATGGTCGTA<br>CCTTTCCGAAACGTGGTCAGACCTGTGTTGTTCATTACACCG<br>GTATGCTGGAAGATGGCAAAAAGTTTGATAGCAGCCGTGATC<br>GTAATAAGCCGTTTAAATTCGTTCTGGGTAAACAAGAAGTTATT<br>CGCGGTTAGGAAGAGGGTGTTGCACAGATGAGCGTTGGTCA<br>GCGTGCAAAACTGACCATTTCACCGGATTATGCATATGGTGC<br>AACCGGTCATCCGGGTATTATTCCGCCTAATGCAACCCTGAT<br>TTTTGATGTTGAACTGCTGAAACTGGAAGGTGGTAGCCATCAC<br>CATCATCATCATTAA | MGVQVETISPGDG<br>RTFPKRGQTCVVH<br>YTGMLEDGKKFDS<br>SRDRNKPFKFVLG<br>KQEVIRGXEEGVA<br>QMSVGQRAKLTIS<br>PDYAYGATGHPGII<br>PPNATLIFDVELLK<br>LEGGSHHHHHH |
| H11-<br>W37TAG | ATGGGCTCTGAAGTTCAGCTGCTGGAATCTGGTGGTGGTCTG<br>GTTCAGGCGGGTGACTCTCTGCGTCTGTCTTGCGCGGCGTC<br>TGGTCGTACCTTCTCTGCGTACGCGATGGGTTGGTTCCGTCA<br>GGCGCCGGGTAAAGAACGTGAATTCGTTGCGGCGATCTCTT<br>GGTCTGGTAACTCTACCTACTACGCGGACTCTGTTAAAGGTC<br>GTTTCACCATCTCTCGTGACAACGCGAAAAACACCGTTTACC<br>TGCAGATGAACTCTCTGAAACCGGAGGACACCGCGATCTAC<br>TACTGCGCGGCGCGTAAACCGATGTACCGTGTTGACATCTC<br>TAAAGGTCAGAACTACGACTACTGGGGTCAGGGTACCCAGG<br>TTACCGTTTCTTCTCACCACCACCACCACCACTAA | MGSEVQLLESGG<br>GLVQAGDSLRLSC<br>AASGRTFSAYAMG<br>XFRQAPGKEREFV<br>AAISWSGNSTYYA<br>DSVKGRFTISRDN<br>AKNTVYLQMNSLK<br>PEDTAIYYCAARKP<br>MYRVDISKGQNYD<br>YWGQGTQVTVSS<br>HHHHHH |

| | | |
|---|---|---|
| H11-<br>F48TAG | ATGGGCTCTGAAGTTCAGCTGCTGGAATCTGGTGGTGGTCTG<br>GTTCAGGCGGGTGACTCTCTGCGTCTGTCTTGCGCGGCGTC<br>TGGTCGTACCTTCTCTGCGTACGCGATGGGTTGGTTCCGTCA<br>GGCGCCGGGTAAAGAACGTGAATAGGTTGCGGCGATCTCTT<br>GGTCTGGTAACTCTACCTACTACGCGGACTCTGTTAAAGGTC<br>GTTTCACCATCTCTCGTGACAACGCGAAAAACACCGTTTACC<br>TGCAGATGAACTCTCTGAAACCGGAGGACACCGCGATCTAC<br>TACTGCGCGGCGCGTAAACCGATGTACCGTGTTGACATCTC<br>TAAAGGTCAGAACTACGACTACTGGGGTCAGGGTACCCAGG<br>TTACCGTTTCTTCTCACCACCACCACCACCACTAA | MGSEVQLLESGG<br>GLVQAGDSLRLSC<br>AASGRTFSAYAMG<br>WFRQAPGKEREX<br>VAAISWSGNSTYY<br>ADSVKGRFTISRD<br>NAKNTVYLQMNSL<br>KPEDTAIYYCAARK<br>PMYRVDISKGQNY<br>DYWGQGTQVTVS<br>SHHHHHH |
| H11-<br>Y60TAG | ATGGGCTCTGAAGTTCAGCTGCTGGAATCTGGTGGTGGTCTG<br>GTTCAGGCGGGTGACTCTCTGCGTCTGTCTTGCGCGGCGTC<br>TGGTCGTACCTTCTCTGCGTACGCGATGGGTTGGTTCCGTCA<br>GGCGCCGGGTAAAGAACGTGAATTCGTTGCGGCGATCTCTT<br>GGTCTGGTAACTCTACCTAGTACGCGGACTCTGTTAAAGGTC<br>GTTTCACCATCTCTCGTGACAACGCGAAAAACACCGTTTACC<br>TGCAGATGAACTCTCTGAAACCGGAGGACACCGCGATCTAC<br>TACTGCGCGGCGCGTAAACCGATGTACCGTGTTGACATCTC<br>TAAAGGTCAGAACTACGACTACTGGGGTCAGGGTACCCAGG<br>TTACCGTTTCTTCTCACCACCACCACCACCACTAA | MGSEVQLLESGG<br>GLVQAGDSLRLSC<br>AASGRTFSAYAMG<br>WFRQAPGKEREFV<br>AAISWSGNSTXYA<br>DSVKGRFTISRDN<br>AKNTVYLQMNSLK<br>PEDTAIYYCAARKP<br>MYRVDISKGQNYD<br>YWGQGTQVTVSS<br>HHHHHH |
| H11-<br>W54TAG | ATGGGCTCTGAAGTTCAGCTGCTGGAATCTGGTGGTGGTCTG<br>GTTCAGGCGGGTGACTCTCTGCGTCTGTCTTGCGCGGCGTC<br>TGGTCGTACCTTCTCTGCGTACGCGATGGGTTGGTTCCGTCA<br>GGCGCCGGGTAAAGAACGTGAATTCGTTGCGGCGATCTCTT<br>AGTCTGGTAACTCTACCTATTACGCGGACTCTGTTAAAGGTC<br>GTTTCACCATCTCTCGTGACAACGCGAAAAACACCGTTTACC<br>TGCAGATGAACTCTCTGAAACCGGAGGACACCGCGATCTAC<br>TACTGCGCGGCGCGTAAACCGATGTACCGTGTTGACATCTC<br>TAAAGGTCAGAACTACGACTACTGGGGTCAGGGTACCCAGG<br>TTACCGTTTCTTCTCACCACCACCACCACCACTAA | MGSEVQLLESGG<br>GLVQAGDSLRLSC<br>AASGRTFSAYAMG<br>WFRQAPGKEREFV<br>AAISXSGNSTYYAD<br>SVKGRFTISRDNAK<br>NTVYLQMNSLKPE<br>DTAIYYCAARKPMY<br>RVDISKGQNYDY<br>WGQGTQVTVSSH<br>HHHHH |
| RFP-Tev-<br>ArcNL-TST-<br>His$_6$ | ATGGCCAGTAGTGAAGACGTTATCAAGGAGTTTATGCGTTTCA<br>AAGTACGTATGGAGGGTAGTGTTAACGGACACGAATTTGAGA<br>TCGAGGGAGAGGGGGAAGGTCGTCCTTACGAGGGAACTCA<br>AACGGCCAAATTAAAGGTGACCAAAGGTGGCCCTTGCCAT<br>TCGCGTGGGACATCTTGTCACCCCAGTTCCAGTACGGGTCG<br>AAGGCATACGTAAAACACCCAGCGGACATTCCTGACTATCTT<br>AAGTTATCTTTCCCGGAAGGTTTTAAATGGGAACGCGTGATGA<br>ACTTTGAGGATGGGGGGGTTGTTACGGTGACACAAGACTCCT<br>CATTGCAAGATGGAGAGTTTATCTATAAAGTCAAACTTCGCGG<br>CACCAATTTTCCATCTGACGGTCCTGTAATGCAGAAAAAAC<br>AATGGGCTGGGAAGCCTCCACAGAACGTATGTACCCCGAAG<br>ATGGAGCTTTAAAGGGCGAAATTAAAATGCGCTTAAAACTTAA | MASSEDVIKEFMR<br>FKVRMEGSVNGH<br>EFEIEGEGEGRPYE<br>GTQTAKLKVTKGG<br>PLPFAWDILSPQF<br>QYGSKAYVKHPAD<br>IPDYLKLSFPEGFK<br>WERVMNFEDGGV<br>VTVTQDSSLQDGE<br>FIYKVKLRGTNFPS<br>DGPVMQKKTMG<br>WEASTERMYPED |

| | | |
|---|---|---|
| | AGACGGCGGCCATTACGACGCCGAAGTGAAAACGACGTATA<br>TGGCTAAGAAACCCGTCCAGCTTCCGGGAGCCTATAAAACT<br>GACATCAAACTGGATATTACATCACACAACGAAGATTATACTA<br>TTGTCGAACAGTACGAACGCGCCGAAGGCCGCCATTCAAC<br>GGGAGCAGGTGGATCAGAAAACCTGTACTTTCAAGGTGCGA<br>TGGGGCCGGGTGTGGACACGCAAATTTTTGAGGATCCCCGC<br>GAATTTTTAAGTCACTTAGAGGAGTATCTTCGCCAAGTTGGCG<br>GTTCAGAAGAGTACTGGCTGTCTCAGATCCAGAACCACATGA<br>ACGGTCCTGCCAAAAAGTGGTGGGAGTTTAAGCAAGGATCC<br>GTCAAAAACTGGGTCGAATTTAAGAAGGAGTTTTTACAGTACA<br>GCGAGGGATGGTCCCACCCGCAGTTCGAAAAAGGTGGTGG<br>TTCTGGTGGTGGTTCTGGTGGTTCTGCGTGGTCCCACCCGC<br>AGTTCGAAAAACACCACCACCACCACCACTAA | GALKGEIKMRLKLK<br>DGGHYDAEVKTTY<br>MAKKPVQLPGAYK<br>TDIKLDITSHNEDY<br>TIVEQYERAEGRH<br>STGAGGSENLYFQ<br>GAMGPGVDTQIFE<br>DPREFLSHLEEYLR<br>QVGGSEEYWLSQI<br>QNHMNGPAKKW<br>WEFKQGSVKNWV<br>EFKKEFLQYSEGW<br>SHPQFEKGGGSG<br>GGSGGSAWSHPQ<br>FEKHHHHHH |
| OmpX-SDL-TAG[b] | ATGAAAAAGATTGCATGTCTGAGCGCACTGGCAGCAGTTCTG<br>GCATTTACCGCAGGCACCAGCGTTGCAGCAACCAGCACCG<br>TTACCGGTGGTTATGCACAGAGTGATGCACAGGGTCAGATGA<br>ATAAAATGGGTGGCTTTAATCTGAAATATCGCTACGAAGAAGA<br>TAATAGTCCGCTGGGTGTTATTGGTAGCTTTACCTATACCGAA<br>AAATCAAGAACCGCAAGCGCGGCCGCAGCGTAGGCAGCC<br>AGCGGTGATTATAACAAAAATCAGTATTATGGCATTACAGCCG<br>GTCCGGCATATCGTATTAATGATTGGGCAAGCATTTATGGTGT<br>TGTTGGTGTTGGTTATGGCAAATTTCAGACCACCGAATATCCG<br>ACCTATAAACATGATACCAGCGATTATGGTTTTAGCTATGGTG<br>CAGGTCTGCAGTTTAATCCGATGGAAAATGTTGCACTGGATTT<br>CAGCTATGAACAGAGCCGTATTCGTAGCGTTGATGTTGGCAC<br>CTGGATTGCCGGTGTGGGTTATCGTTTTCACCATCACCATCA<br>CCATTAA | MKKIACLSALAAVL<br>AFTAGTSVAATSTV<br>TGGYAQSDAQGQ<br>MNKMGGFNLKYR<br>YEEDNSPLGVIGSF<br>TYTEKSRTASAAAA<br>XAASGDYNKNQYY<br>GITAGPAYRINDW<br>ASIYGVVGVGYGK<br>FQTTEYPTYKHDTS<br>DYGFSYGAGLQFN<br>PMENVALDFSYEQ<br>SRIRSVDVGTWIA<br>GVGYRFHHHHHH |
| OmpX-W76TAG | ATGAAAAAGATTGCATGTCTGAGCGCACTGGCAGCAGTTCTG<br>GCATTTACCGCAGGCACCAGCGTTGCAGCAACCAGCACCG<br>TTACCGGTGGTTATGCACAGAGTGATGCACAGGGTCAGATGA<br>ATAAAATGGGTGGCTTTAATCTGAAATATCGCTACGAAGAAGA<br>TAATAGTCCGCTGGGTGTTATTGGTAGCTTTACCTATACCGAA<br>AAATCAAGAACCGCAAGCGCGGCCGCAGCGGCAGCAGCC<br>AGCGGTGATTATAACAAAAATCAGTATTATGGCATTACAGCCG<br>GTCCGGCATATCGTATTAATGATTAGGCAAGCATTTATGGTGT<br>TGTTGGTGTTGGTTATGGCAAATTTCAGACCACCGAATATCCG<br>ACCTATAAACATGATACCAGCGATTATGGTTTTAGCTATGGTG<br>CAGGTCTGCAGTTTAATCCGATGGAAAATGTTGCACTGGATTT<br>CAGCTATGAACAGAGCCGTATTCGTAGCGTTGATGTTGGCAC<br>CTGGATTGCCGGTGTGGGTTATCGTTTTCACCATCACCATCA<br>CCATTAA | MKKIACLSALAAVL<br>AFTAGTSVAATSTV<br>TGGYAQSDAQGQ<br>MNKMGGFNLKYR<br>YEEDNSPLGVIGSF<br>TYTEKSRTASAAAA<br>AAASGDYNKNQY<br>YGITAGPAYRINDX<br>ASIYGVVGVGYGK<br>FQTTEYPTYKHDTS<br>DYGFSYGAGLQFN<br>PMENVALDFSYEQ<br>SRIRSVDVGTWIA<br>GVGYRFHHHHHH |

| | | |
|---|---|---|
| OmpX-<br>W140TAG | ATGAAAAAGATTGCATGTCTGAGCGCACTGGCAGCAGTTCTG<br>GCATTTACCGCAGGCACCAGCGTTGCAGCAACCAGCACCG<br>TTACCGGTGGTTATGCACAGAGTGATGCACAGGGTCAGATGA<br>ATAAAATGGGTGGCTTTAATCTGAAATATCGCTACGAAGAAGA<br>TAATAGTCCGCTGGGTGTTATTGGTAGCTTTACCTATACCGAA<br>AAATCAAGAACCGCAAGCGCGGCCGCAGCGGCAGCAGCC<br>AGCGGTGATTATAACAAAAATCAGTATTATGGCATTACAGCCG<br>GTCCGGCATATCGTATTAATGATTGGGCAAGCATTTATGGTGT<br>TGTTGGTGTTGGTTATGGCAAATTTCAGACCACCGAATATCCG<br>ACCTATAAACATGATACCAGCGATTATGGTTTTAGCTATGGTG<br>CAGGTCTGCAGTTTAATCCGATGGAAAATGTTGCACTGGATTT<br>CAGCTATGAACAGAGCCGTATTCGTAGCGTTGATGTTGGCAC<br>CTAGATTGCCGGTGTGGGTTATCGTTTTCACCATCACCATCA<br>CCATTAA | MKKIACLSALAAVL<br>AFTAGTSVAATSTV<br>TGGYAQSDAQGQ<br>MNKMGGFNLKYR<br>YEEDNSPLGVIGSF<br>TYTEKSRTASAAAA<br>AAASGDYNKNQY<br>YGITAGPAYRIND<br>WASIYGVVGVGYG<br>KFQTTEYPTYKHDT<br>SDYGFSYGAGLQF<br>NPMENVALDFSYE<br>QSRIRSVDVGTXIA<br>GVGYRFHHHHHH |
| G1NBDO53<br>(for yeast<br>expression) | ATGGTCGTTAAGTTTACAGATTCTCAAATCCAACACTTGATGG<br>AATACGGTGACAACGATTGGTCTGAAGCTGAATTCGAAGACG<br>CCGCTGCCAGAGACAAAGAATTCTCCTCTCAATTCTCCAAAT<br>TGAAATCTGCTAACGACAAGGGTCTAAAGGATGTTATTGCTAA<br>TCCACGTAATGATTTGACCGACTTAGAAAATAAGATTAGAGAA<br>AAGTTGGCTGCCAGAGGTTTTATCGAGGTCCATACTCCAATC<br>TTCGTTTCCAAATCTGCCTTGGCTAAGATGACTATTACTGAAG<br>ACCACCCATTGTTCAAGCAAGTTTTCTGGATTGATGACAAAAG<br>AGCCTTGCGTCCAATGCATGCCATGAACATGTACAAGGTCAT<br>GAGAGAATTGAGAGATCACACTAAGGGTCCAGTCAAGATTTT<br>CGAAATTGGTTCCTGTTTCAGAAAGGAATCTAAGTCCTCTACC<br>CACTTGGAAGAGTTCACTATGTTGAACTTAGTTGAAATGGGTC<br>CAGATGGTGATCCAATGGAACATTTGAAGATGTATATCGGTGA<br>CATTATGGACGCCGTCGGTGTCGAATACACCACTTCTAGAGA<br>GGAATCAGATGTCTACGTCGAAACCTTGGACGTTGAGATCAA<br>CGGCACTGAAGTTGCTTCAGGCGGTGTTGGTCCACACAAGT<br>TAGATCCAGCCCACGACGTTCACGAACCTTACGCTGGTATTG<br>GTTTCGGTTTGGAGAGATTATTGATGTTGAAAAACGGTAAATCT<br>AATGCCAGAAAGACCGGTAAATCTATTACTTACTTGAATGGCT<br>ACAAGTTGGACTAA | MVVKFTDSQIQHL<br>MEYGDNDWSEAE<br>FEDAAARDKEFSS<br>QFSKLKSANDKGL<br>KDVIANPRNDLTD<br>LENKIREKLAARGF<br>IEVHTPIFVSKSALA<br>KMTITEDHPLFKQV<br>FWIDDKRALRPMH<br>AMNMYKVMRELR<br>DHTKGPVKIFEIGS<br>CFRKESKSSTHLEE<br>FTMLNLVEMGPD<br>GDPMEHLKMYIG<br>DIMDAVGVEYTTS<br>REESDVYVETLDVE<br>INGTEVASGGVGP<br>HKLDPAHDVHEPY<br>AGIGFGLERLLMLK<br>NGKSNARKTGKSI<br>TYLNGYKLD |
| Electra2-<br>TAG-<br>mScarlet3<br>(for yeast<br>expression) | ATGCATCACCATCACCATCACGGGTCTGTGAGCAAAGGCGA<br>GGAACTCATCGAAGAAAACATGAGGATGAAAGTGGTTATGGA<br>AGGAAGCGTGAATGGCCACCAATTTAAATGTACTGGGGAGG<br>GCGAGGGAAGACCATACGAGGGAGTTCAGACAATGAGGATT<br>AAGGTAATTGAGGGTGGGCCTCTTCCTTTCGCCTTTGATATAT<br>TGGCTACATCTTTCTTATTCGGCTCCAAGACCTTCATAAAATA<br>CCCTGCTGATATTCCAGACTTTTTTGAACAGTCTTTCCCTGAA<br>GGGTTTACATGGGAACGAGTTACCAGATACGAGGATGGAGG<br>TGTCGTCACTGTCACACAGGACACATCTCTGGAAGATGGGG<br>GATTGGTTTACAACGTGAAGGTCAGAGGTGTCAACTTCCACA<br>GCAAAGGCCCTGTCATGCAAAAAAAGACTGAGGGTTGGGAG<br>CCTTTTACTGAGATGATGTACCCCGCCGACGGGGGCCTGAG<br>GGGCTACACAGACATTGCATTGAAAGTGGATGGAGGCGGCC | MHHHHHHGSVSK<br>GEELIEENMRMKV<br>VMEGSVNGHQFK<br>CTGEGEGRPYEGV<br>QTMRIKVIEGGPLP<br>FAFDILATSFLFGS<br>KTFIKYPADIPDFFE<br>QSFPEGFTWERVT<br>RYEDGGVVTVTQD<br>TSLEDGGLVYNVK<br>VRGVNFHSKGPV<br>MQKKTEGWEPFTE<br>MMYPADGGLRGY |

| | | |
|---|---|---|
| | ACTTGCATGCAAACATTGTAACAACTTACAGATCCAAGAAGA | TDIALKVDGGGHL |
| | CTGTAGGGAACATTAAAATGCCTGGGGTACATGCAGTGGACT | HANIVTTYRSKKTV |
| | ACAGGCTCGAACGAATTGAGGAATCTGATAATGAGACATATG | GNIKMPGVHAVDY |
| | TTGTTCTCAGGGAAGTGGCCGTTGCTAAATACTCCAATCTCG | RLERIEESDNETYV |
| | GTGGGGGAATGGATGAATTGTTTAAGGGTTCTGGTGAAAATTT | VLREVAVAKYSNL |
| | GTATTTTCAATCTATGGGATCCGCTGGCTCCGCTGCTGGTTCT | GGGMDELFKGSG |
| | GGCGAATAGGCAGGTGGGGATTCTACCGAAGCTGTCATTAA | ENLYFQSMGSAG |
| | GGAATTTATGAGATTCAAGGTTCATATGGAAGGTTCTATGAAC | SAAGSGEXAGGD |
| | GGTCACGAGTTCGAAATTGAAGGTGAAGGTGAAGGTCGTCC | STEAVIKEFMRFKV |
| | ATATGAAGGTACCCAAACCGCCAAGTTGAGAGTTACTAAAGG | HMEGSMNGHEFE |
| | CGGTCCATTGCCATTCTCTTGGGATATTTTGTCCCCACAATTT | IEGEGEGRPYEGT |
| | ATGTACGGTTCAAGAGCTTTCACTAAACACCCAGCTGACATC | QTAKLRVTKGGPL |
| | CCTGACTACTGGAAACAATCTTTTCCAGAAGGTTTCAAGTGG | PFSWDILSPQFMY |
| | GAGAGAGTCATGAACTTTGAAGATGGCGGTGCTGTCTCAGTT | GSRAFTKHPADIP |
| | GCTCAAGATACTTCTTTGGAAGATGGTACTTTAATTTATAAGGT | DYWKQSFPEGFK |
| | TAAGTTGAGAGGCACTAACTTCCCTCCAGATGGTCCAGTTAT | WERVMNFEDGGA |
| | GCAAAAAAAGACCATGGGTTGGGAAGCTTCCACCGAAAGAC | VSVAQDTSLEDGT |
| | TATATCCTGAAGACGTCGTTTTGAAGGGTGATATTAAGATGGC | LIYKVKLRGTNFPP |
| | CTTGAGATTGAAGGACGGTGGCAGATACTTGGCTGATTTCAA | DGPVMQKKTMG |
| | GACTACCTACAGAGCTAAAAAGCCAGTTCAAATGCCTGGTGC | WEASTERLYPEDV |
| | TTTTAATATTGATAGAAAGTTGGACATTACCTCACACAACGAAG | VLKGDIKMALRLK |
| | ACTATACCGTTGTCGAACAATACGAGCGTTCTGTTGCTAGAC | DGGRYLADFKTTY |
| | ACTCCACTGGCGGGTCATCCTAA | RAKKPVQMPGAF |
| | | NIDRKLDITSHNED |
| | | YTVVEQYERSVAR |
| | | HSTGGSS |
| MFapp8- | ATGAGATTCCCATCTATCTTCACCGCTGTTTTGTTCGCTGCTT | MRFPSIFTAVLFAA |
| H11- | CTTCTGCTTTGGCTGCTCCAGCTAACACCACCACCGAAGAC | SSALAAPANTTTED |
| W37TAG- | GAAACCGCTCAAATCCCAGCTGAAGCTGTTATCGACTACTCT | ETAQIPAEAVIDYS |
| Sed1p | GACTTGGAAGGTGACTTCGACGCTGCTGCTTTGCCATTGTCT | DLEGDFDAAALPL |
| (for yeast | AACTCTACCAACAACGGTTTGTCTTCTACCAACACCACCATC | SNSTNNGLSSTNT |
| expression) | GCTTCTATCGCTGCTAAGGAAGAAGGTGTTCAATTGGACAAG | TIASIAAKEEGVQL |
| | AGAGAAGCTGATGCTTCTGGTTCCGAAGTCCAATTATTGGAAT | DKREADASGSEVQ |
| | CTGGTGGCGGTTTGGTTCAAGCCGGTGATTCCTTGAGATTGT | LLESGGGLVQAG |
| | CTTGTGCCGCTTCCGGTCGTACCTTTTCCGCTTACGCCATGG | DSLRLSCAASGRT |
| | GTTAGTTCAGACAAGCTCCAGGTAAAGAAAGAGAATTTGTTGC | FSAYAMGXFRQAP |
| | TGCCATTTCATGGTCTGGTAACTCCACTTATTACGCTGACTCT | GKEREFVAAISWS |
| | GTTAAGGGCAGATTTACCATTTCAAGAGACAACGCCAAGAAC | GNSTYYADSVKGR |
| | ACTGTTTACTTGCAAATGAACTCTTTGAAACCTGAAGACACCG | FTISRDNAKNTVYL |
| | CCATTTATTACTGCGCTGCCAGAAAACCAATGTACAGAGTTG | QMNSLKPEDTAIY |
| | ACATTTCCAAAGGTCAAAATTACGATTACTGGGGTCAAGGCA | YCAARKPMYRVDI |
| | CCCAAGTCACTGTTTCTTCCCATCACCATCACCACCACGGAT | SKGQNYDYWGQ |
| | CCGGTGGTGGTGGTTCAGGCGGCGGTGGTTCCTATCCCTAT | GTQVTVSSHHHH |
| | GACGTACCCGACTATGCGAAAGAGAACTTATACTTCCAGAGC | HHGSGGGGSGG |
| | CAGTTCTCTAATTCAACGTCCGCCTCGTCTACAGACGTTACC | GGSYPYDVPDYAK |
| | TCTTCTTCGTCTATATCTACCTCGTCCGGGAGCGTCACGATTA | ENLYFQSQFSNST |
| | CCTCATCCGAGGCACCTGAGAGCGATAATGGCACTTCTACC | SASSTDVTSSSSIS |
| | GCAGCCCCCACGGAGACTAGCACCGAAGCCCCTACAACC | TSSGSVTITSSEAP |

| | | |
|---|---|---|
| | GCCATTCCTACCAATGGCACTTCAACAGAGGCACCCACAAC<br>AGCAATACCCACAAATGGAACTTCAACCGAGGCCCCTACGG<br>ACACAACAACAGAGGCACCTACGACTGCATTACCGACAAAT<br>GGCACCTCGACAGAGGCACCTACCGACACAACCACAGAG<br>GCACCCACTACAGGCTTGCCTACAAATGGAACTACAAGCGC<br>GTTTCCCCCTACGACGTCACTTCCCCCTAGTAATACAACGAC<br>AACACCACCATATAATCCCTCTACAGATTATACTACGGATTAT<br>ACAGTCGTTACAGAGTACACAACATATTGCCCTGAGCCGAC<br>GACCTTTACAACGAATGGCAAAACATATACTGTGACGGAGCC<br>GACAACTTTAACAATTACCGATTGCCCCTGTACTATCGAGAAA<br>CCTACTACTACCTCTACAACAGAGTATACCGTTGTGACAGAAT<br>ATACCACGTATTGCCCTGAGCCTACTACGTTTACTACTAATGG<br>AAAGACCTATACTGTTACAGAGCCGACTACGCTAACAATAAC<br>AGATTGCCCTTGCACGATAGAGAAATCTGAGGCTCCAGAATC<br>AAGCGTGCCGGTGACGGAGAGCAAAGGTACAACGACAAAG<br>GAGACTGGCGTCACAACTAAGCAGACGACCGCTAATCCTTC<br>TTTAACGGTGTCGACGGTGGTTCCCGTGTCTTCTAGTGCATC<br>CTCGCACTCAGTCGTAATTAATAGCAATGGCGCAAATGTAGTT<br>GTACCGGGAGCACTGGGATTAGCAGGCGTGGCAATGCTGTT<br>CTTGTAA | ESDNGTSTAAPTET<br>STEAPTTAIPTNGT<br>STEAPTTAIPTNGT<br>STEAPTDTTTEAPT<br>TALPTNGTSTEAPT<br>DTTTEAPTTGLPTN<br>GTTSAFPPTTSLPP<br>SNTTTTPPYNPSTD<br>YTTDYTVVTEYTTY<br>CPEPTTFTTNGKTY<br>TVTEPTTLTITDCPC<br>TIEKPTTTSTTEYTV<br>VTEYTTYCPEPTTFT<br>TNGKTYTVTEPTTL<br>TITDCPCTIEKSEAP<br>ESSVPVTESKGTTT<br>KETGVTTKQTTANP<br>SLTVSTVVPVSSSA<br>SSHSVVINSNGAN<br>VVVPGALGLAGVA<br>MLFL |
| MFapp8-<br>H11-Nt-<br>TAG-Sed1p<br>(for yeast<br>expression) | ATGAGATTCCCATCTATCTTCACCGCTGTTTTGTTCGCTGCTT<br>CTTCTGCTTTGGCTGCTCCAGCTAACACCACCACCGAAGAC<br>GAAACCGCTCAAATCCCAGCTGAAGCTGTTATCGACTACTCT<br>GACTTGGAAGGTGACTTCGACGCTGCTGCTTTGCCATTGTCT<br>AACTCTACCAACAACGGTTTGTCTTCTACCAACACCACCATC<br>GCTTCTATCGCTGCTAAGGAAGAAGGTGTTCAATTGGACAAG<br>AGAGAAGCTGATGCTTCTTAGGGTTCCGAAGTCCAATTATTGG<br>AATCTGGTGGCGGTTTGGTTCAAGCCGGTGATTCCTTGAGATT<br>GTCTTGTGCCGCTTCCGGTCGTACCTTTTCCGCTTACGCCAT<br>GGGTTGGTTCAGACAAGCTCCAGGTAAAGAAAGAGAATTTGT<br>TGCTGCCATTTCATGGTCTGGTAACTCCACTTATTACGCTGAC<br>TCTGTTAAGGGCAGATTTACCATTTCAAGAGACAACGCCAAG<br>AACACTGTTTACTTGCAAATGAACTCTTTGAAACCTGAAGACA<br>CCGCCATTTATTACTGCGCTGCCAGAAAACCAATGTACAGAG<br>TTGACATTTCCAAAGGTCAAAATTACGATTACTGGGGTCAAGG<br>CACCCAAGTCACTGTTTCTTCCCATCACCATCACCACCACG<br>GATCCGGTGGTGGTGGTTCAGGCGGCGGTGGTTCCTATCCC<br>TATGACGTACCCGACTATGCGAAAGAGAACTTATACTTCCAG<br>AGCCAGTTCTCTAATTCAACGTCCGCCTCGTCTACAGACGTT<br>ACCTCTTCTTCGTCTATATCTACCTCGTCCGGGAGCGTCACG<br>ATTACCTCATCCGAGGCACCTGAGAGCGATAATGGCACTTCT<br>ACCGCAGCCCCCACGGAGACTAGCACCGAAGCCCCTACA<br>ACCGCCATTCCTACCAATGGCACTTCAACAGAGGCACCCAC<br>AACAGCAATACCCACAAATGGAACTTCAACCGAGGCCCCTA<br>CGGACACAACAACAGAGGCACCTACGACTGCATTACCGAC<br>AAATGGCACCTCGACAGAGGCACCTACCGACACAACCACA<br>GAGGCACCCACTACAGGCTTGCCTACAAATGGAACTACAAG | MRFPSIFTAVLFAA<br>SSALAAPANTTTED<br>ETAQIPAEAVIDYS<br>DLEGDFDAAALPL<br>SNSTNNGLSSTNT<br>TIASIAAKEEGVQL<br>DKREADASXGSEV<br>QLLESGGGLVQA<br>GDSLRLSCAASGR<br>TFSAYAMGWFRQ<br>APGKEREFVAAIS<br>WSGNSTYYADSVK<br>GRFTISRDNAKNT<br>VYLQMNSLKPEDT<br>AIYYCAARKPMYR<br>VDISKGQNYDYW<br>GQGTQVTVSSHH<br>HHHHGSGGGGS<br>GGGGSYPYDVPD<br>YAKENLYFQSQFS<br>NSTSASSTDVTSSS<br>SISTSSGSVTITSSE<br>APESDNGTSTAAP<br>TETSTEAPTTAIPTN<br>GTSTEAPTTAIPTN<br>GTSTEAPTDTTTEA<br>PTTALPTNGTSTEA |

| | | |
|---|---|---|
| | CGCGTTTCCCCCTACGACGTCACTTCCCCCTAGTAATACAA | PTDTTTEAPTTGLP |
| | CGACAACACCACCATATAATCCCTCTACAGATTATACTACGG | TNGTTSAFPPTTSL |
| | ATTATACAGTCGTTACAGAGTACACAACATATTGCCCTGAGC | PPSNTTTTPPYNPS |
| | CGACGACCTTTACAACGAATGGCAAAACATATACTGTGACGG | TDYTTDYTVVTEYT |
| | AGCCGACAACTTTAACAATTACCGATTGCCCCTGTACTATCG | TYCPEPTTFTTNGK |
| | AGAAACCTACTACTACCTCTACAACAGAGTATACCGTTGTGA | TYTVTEPTTLTITDC |
| | CAGAATATACCACGTATTGCCCTGAGCCTACTACGTTTACTAC | PCTIEKPTTTSTTEY |
| | TAATGGAAAGACCTATACTGTTACAGAGCCGACTACGCTAAC | TVVTEYTTYCPEPT |
| | AATAACAGATTGCCCTTGCACGATAGAGAAATCTGAGGCTCC | TFTTNGKTYTVTEP |
| | AGAATCAAGCGTGCCGGTGACGGAGAGCAAAGGTACAACG | TTLTITDCPCTIEKS |
| | ACAAAGGAGACTGGCGTCACAACTAAGCAGACGACCGCTAA | EAPESSVPVTESKG |
| | TCCTTCTTTAACGGTGTCGACGGTGGTTCCCGTGTCTTCTAGT | TTTKETGVTTKQTT |
| | GCATCCTCGCACTCAGTCGTAATTAATAGCAATGGCGCAAAT | ANPSLTVSTVVPVS |
| | GTAGTTGTACCGGGAGCACTGGGATTAGCAGGCGTGGCAAT | SSASSHSVVINSN |
| | GCTGTTCTTGTAA | GANVVVPGALGLA |
| | | GVAMLFL |

[a] X indicates the positions of BD-ncAAs;

[b] An artificial surface-display loop consisting of seven alanine residues was inserted between S53 and S54 of OmpX, with the fifth alanine replaced by an amber (TAG) codon.

Supplementary Table 5. Fluorescence emission maxima of CBDS in different solvents and protein environments used for TCSPC measurements.

| Sample[a] | $\lambda_{ex}$, max (nm) |
|---|---|
| Water | 600 |
| 1:1 Ethylene glycol:Water | 600 |
| 9:1 Ethylene glycol:Water | 599 |
| DMSO | 599 |
| Methanol | 597 |
| Ethanol | 590 |
| Acetone | 581 |
| Chloroform | 572 |
| Dichloromethane | 563 |
| H11-W37CBDS | 505 |
| H11-F48CBDS | 550 |
| H11-W54CBDS | 575 |
| H11-Y60CBDS | 565 |
| H11-W37CBDS + ArcNL | 505 |
| H11-F48CBDS + ArcNL | 555 |

| | |
|---|---|
| H11-W54CBDS + ArcNL | 560 |
| H11-Y60CBDS + ArcNL | 560 |

[a] Solvent screen samples contain 0.1% DMSO.

Supplementary Table 6. Crystallographic statistics of nanobody ArcNL H11-W37NBDO and H11-W37NBDSe.

| | ArcNL H11-W37NBDO | ArcNL H11-W37NBDSe |
|---|---|---|
| **PDB ID** | 9PZZ | 9Q00 |
| **Data collection** | | |
| Source | Australian Synchrotron MX2 | Australian Synchrotron MX2 |
| Space group | $C\ 1\ 2_1\ 1$ | $C\ 1\ 2_1\ 1$ |
| Cell dimensions | | |
| *a, b, c* (Å) | 113.0 44.1 88.8 | 113.8 44.3 89.0 |
| α, β, γ (°) | 90.0 92.7 90.0 | 90.0 98.2 90.0 |
| Resolution (Å) | 48.68 – 1.56 (1.58 – 1.56) | 44.65 – 2.84 (3.13 – 2.84) |
| $R_{merge}$ | 0.105 (1.107) | 0.433 (1.107) |
| $R_{pim}$ | 0.066 (0.690) | 0.286 (0.755) |
| $I/\sigma I$ | 11.5 (1.6) | 6.1 (1.7) |
| $CC_{1/2}$ | 0.998 (0.442) | 0.951 (0.434) |
| Completeness (%) | 100.0 (100.0) | 99.9 (99.8) |
| Redundancy | 6.8 (6.9) | 6.1 (5.7) |
| **Refinement** | | |
| Resolution (Å) | 48.68 – 1.56 (1.58 – 1.56) | 44.65 – 2.84 (3.13 – 2.84) |
| No. reflections | 62461 (2704) | 10612 (2634) |
| $R_{work}/R_{free}$ | 0.1856 / 0.2267 (0.2962 / 0.3437) | 0.2156 / 0.2807 (0.2156 / 0.3933) |
| Number of atoms | | |
| Protein | 3138 | 3109 |
| Ligand/ion | 36 | 36 |
| Solvent | 683 | 115 |
| *B*-factors (overall) | 23.8 | 34.7 |
| Protein | 21.8 | 35.2 |
| Ligand/ion | 14.1 | 27.3 |
| Solvent | 33.7 | 21.7 |
| RMS deviations | | |
| Bond lengths (Å) | 0.008 | 0.003 |
| Bond angles (°) | 0.83 | 0.66 |

[a] Values in parentheses are for the highest resolution shell.
